# A new class of human CpG island promoters with primate-specific repeats

**DOI:** 10.1101/2024.01.17.576070

**Authors:** K Naga Mohan, Anuhya Anne, Lov Kumar, J Richard Chaillet

## Abstract

A subset of imprinting control regions (ICRs) in the human and mouse possess CpG islands associated with imperfect tandem repeats (TRs) that were shown to be essential for genomic imprinting through genetic studies. To identify whether this feature is also present in non-imprinted CpG island promoters, we performed extensive dot plot analyses and identified 365 CpG island gene promoters associated with imperfect TRs of ≥ 400nt. These TRs are absent in the orthologous mouse CGI promoters, and most occur as clusters at the human chromosome ends, distinct from the clusters of imprinted genes. These genes showed an enrichment in neurodevelopmental/behavioral disorders and show interindividual variation in methylation levels. A subset of TR-CGIs is highly methylated and remains so during reprogramming to primed iPSCs, but become unmethylated in naïve iPSCs, as has been shown for the ICRs. Transcript levels correlate with methylation levels for some TR-CGI genes suggesting their gene regulatory potential. Orthologs of the subset of methylated TR-CGIs are unmethylated in mouse, suggesting the role of TRs in imparting methylation in humans. The human TR-CGIs accompanied primate evolution after divergence from the rodent lineage with evidence of recent additions in human evolution. In summary, the incorporation of TRs in certain CGI promoters in primate evolution results in the unique ability to acquire methylation during embryonic development and resist reprogramming to a pluripotent stem cell state with a possible effect on gene expression.

## Introduction

About 50% of mammalian gene regulation at the transcription initiation level takes place via CpG-rich promoter regions known as CpG islands (CGIs) that are mostly unmethylated in pre-and post-implantation embryonic development and post-natal life. The others acquire methylation during embryonic development and tend to be transcriptionally silent. A notable exception to the general rule is the parental origin-specific methylation of approximately 20 CG-rich autosomal imprinting control regions (ICRs), ∼1-5 Kb in length. Methylation of these sequences is established during gametogenesis reflecting the parental sex by *de novo* methyltransferases and maintained post-fertilization by the DNA methyltransferase 1 (DNMT1) [1]. As a result, in eutherian mammals, one parental ICR allele is methylated and the counterpart unmethylated. Although there is uncertainty on the sequence requirements for establishing methylation, tandem repeat (TR) sequences have been experimentally shown to be essential for some, but not all, ICRs [3–7]. The absence of sequence similarities between ICRs strongly suggests that repetitiveness majorly contributes to the imprinted state [8].

DNMT1 maintains methylation on all DNA sequences following its replication. This process requires important co-factors such as NP95/UHRF and certain zinc-finger proteins such as ZFP57 and ZNF445 for perpetuation of ICR methylation with cell division. A salient feature of DNMT1 is its distinct roles in maintaining methylation on ICR and non-ICR sequences. When DNMT1 protein is removed from mouse ES cells by blocking *Dnmt1* transcription, both ICR and non-ICR sequences lose methylation. With re-expression of DNMT1 only non-ICR sequences recover their methylation [9,10]. The *in vivo* deletion of DNMT1o, the oocyte isoform of DNMT1, mirrors these *in vitro* findings – half of ICR methylation is permanently lost, but non-ICR methylation is normal post-implantation [11]. These findings suggest two DNMT1-dependent maintenance functions, one acting on ICRs and the other on non-ICR sequences.

We have identified mutations in *Dnmt1* that maintain only non-ICR methylation, consistent with separate ICR and non-ICR maintenance functions encoded in *Dnmt1* itself [9,12]. Overexpression of DNMT1s, the somatic isoform, is lethal, whereas overexpression of DNMT1o in mice is not lethal. This suggests a non-catalytic function of the DNMT1s-specific amino-terminal domain, which when enhanced is lethal [13,14]. In summary, ICR methylation is a type of epigenetic cellular memory, where, once established, is perpetuated with cell division and differentiation. In contrast, non-ICR methylation is part of a gene regulatory process, instructing gene transcription, and whose level changes in sync with changes in transcription rates [15].

A second feature of monoallelic ICR methylation is its longevity; ICR methylation is established during gametogenesis and persists throughout preimplantation development despite genome-wide hypomethylation. Monoallelic methylation ensures transcription of genes in a dose-dependent manner that is required for normal development in mammals, yet generally not required after birth. For example, many imprinted genes are only expressed in the placenta but monoallelic ICR methylation persists for the lifetime of the individual [16]. From this, we reasoned that CGI methylation akin to ICR methylation may occur on non-imprinted sequences. For example, germline genes are targeted for repression in ES cells by the polycomb repressive complex 1.6 and DNA methylation [17]. Methylation of such sequences, initiated as *de novo* methylation at some developmental stage post-fertilization, and thereafter maintained by DNMT1, might differ from ICR methylation because post-zygotic *de novo* methylation may be cell type-specific and can differ between individuals.

## Results

### Identification of direct repeats in human promoter-proximal CpG islands

To test these hypotheses, we first generated and analyzed dotplots of human promoter-proximal CGIs (see Materials and Methods) and identified 365 with tandem repeats (TRs) measuring ≥400bp each, containing more than two copies of a perfect or imperfect repeating unit. We refer to these sequences as TR-CGIs, of which 241 occur in and 124 within 200 bp of the CGIs. A vast majority of the TR-CGIs are autosomal (347) and were investigated in detail in the context of DNA methylation, epigenetic reprogramming, individual-to-individual variation, and its influence on the transcript levels (Fig. 1A). Dotplots of two TR-CGIs and one ICR are shown as examples in Figures 1B and S2. As is the case for repeats in ICRs, DNA sequence of each TR is unique. Thus, if TRs were to provide similar properties to all TR-CGIs, it must be through their repetitiveness rather than due to a sequence.

**Fig. 1.**
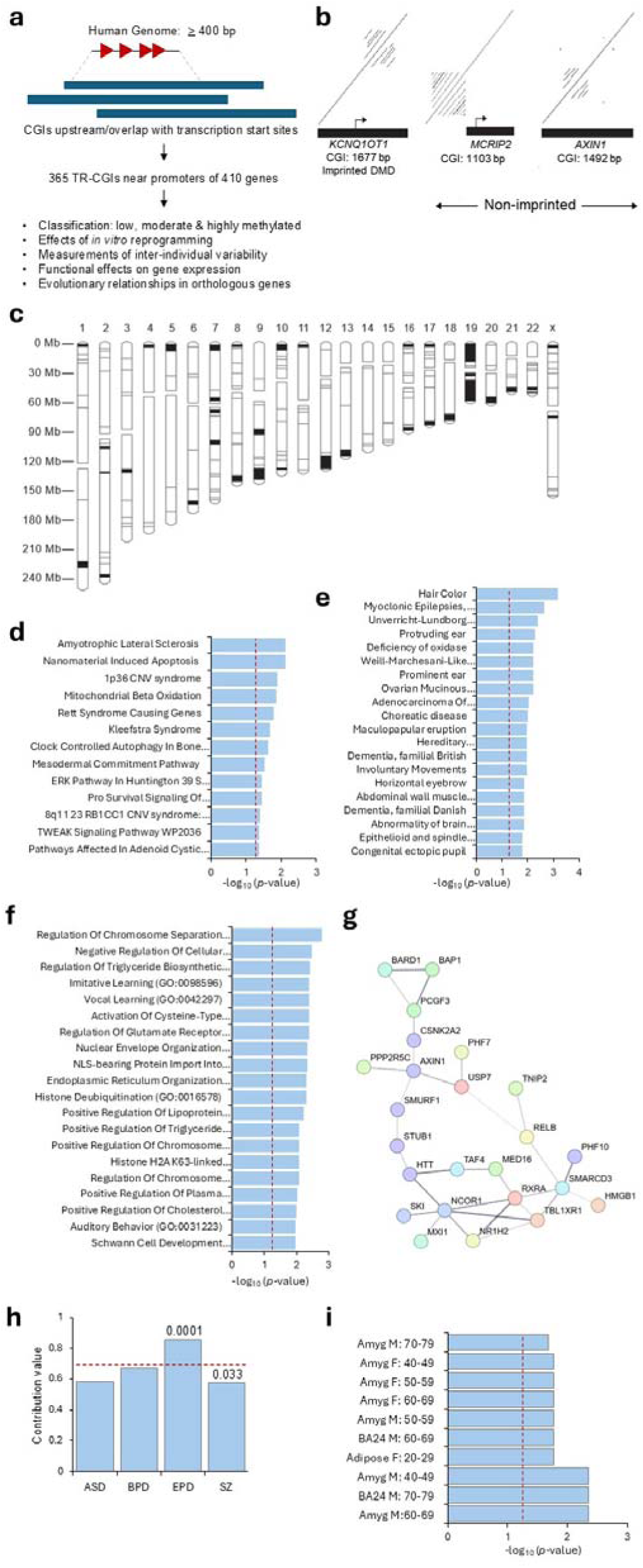
Identification and characterization of human CpG islands (CGIs) upstream or overlapping transcription start sites and containing tandem repeat sequences. **(A)** Schema for determining features of tandem repeat (TR)-containing CGIs at promoter regions. **(B)** Two examples of non-imprinted genes with TRs and their comparison with the *KCNQ1OT1* imprinted region. Lines parallel to the central diagonal indicate the presence of tandem repeats; the number of lines is the number of repeats, and space between lines is the sequence length of a unit copy. **(C)** Chromosomal locations of the identified non-imprinted genes with TRs either in or within 200 bp of CGI promoters. Horizontal lines indicate individual genes whereas filled rectangles indicate gene clusters. **(D-F)** Top twenty most significant terms identified by DisGenet, Wikipathway and Biological process analyses. Vertical dashed red lines represent *p* values of 0.05. **(G)** Protein-protein interaction analysis of genes with TR-CGI promoters. **(H)** Proportions of genes associated with autism spectrum (ASD), bipolar (BPD), epilepsy (EPD) and schizophrenia (SZ). The horizontal dashed line indicates the expected value. The *p* values are given on the top of the histograms. **(I)** GTEx-analysis of genes with TR-CGIs in promoters. Vertical dashed lines represent *p* values of 0.05.

Approximately 60% of TR-CGIs are in 33 autosomal clusters, each with more than two genes, occurring in lengths ∼2.5-23 Mb with ∼0.5-6.0 TR-CGIs/Mb (Fig. 1C and Supplementary Table S1). Many autosomes contain more than one cluster, predominantly at the ends of chromosomes except chromosome 19 that has 51 TR-CGIs (14% of total) intermingled with *ZFP* and *ZNF* genes [18]. CGI density follows that of euchromatin, which is much more broadly distributed than TR-CGIs. This is illustrated in a comparison of a CGI ideogram produced by mapping a library of CGI clones (most single-copy and near TSSs) to the human genome [19,20]. TR-CGI distribution is clearly different from that reported for CGIs in general. An exception to this is chromosome 19, which has a high concentration of CGIs, TR-CGIs, and genes encoding ZNFs. Clustering to chromosome ends is not seen for mouse syntenic regions (Supplementary Table S1). Importantly, TR-CGI clusters do not overlap with the clusters of imprinted genes.

### Biological pathways, diseases and physical associations with TR-CGI genes

Following this preliminary description, we compared wherever necessary, the features and phenomena associated with the 25 ICRs and the remaining 8,332 promoter-associated CGIs that are referred to as non-TR CGIs. Bioinformatic analyses of the autosomal TR-CGI genes by disease and gene ontologies yielded terms that are distinct from those identified for non-TR genes. (Fig. 1D-1F, Supplementary Figure S1). Among the 25 diseases associated with the TR-CGI genes, 11 are neurological, neurodevelopmental, or behavioral disorders (Fig. 1D, 1E). The list of biological processes includes chromosome condensation, chromosome separation, nuclear envelope organization, and regulation of glutamate receptor (Fig. 1F). Protein-protein interaction analysis yielded a single cluster of proteins involved in ubiquitinylation, and nuclear receptor corepressor activities (Fig. 1G). Within neurological/behavioral disorders, there was a significant overrepresentation for epilepsy, but underrepresentation in case of schizophrenia (Fig. 1H and Supplementary Table S2). GTEx analysis confirmed significant association of TR-CGI gene expression in the amygdala with no sex bias (Fig. 1I and Supplementary Table S2).

### A higher proportion of TR-CGIs are methylated than the non-TR CGIs

Given the presence of TRs in the TR-CGIs, the features of these sequences were compared with features of CpG-rich ICRs and non-TR CGIs. GC-content and CpG ratio analysis showed a major overlap between the ICRs and TR-CGIs, whereas non-TRs mostly formed a separate cluster (Figures 2A, S3). The proportions of ICR, TR-CGI and non-TR CGIs showing different levels of methylation were compared using data on 45 human cerebellum samples (Fig. 2B) [21], All ICRs showed methylation levels between 40% and 70%, whereas a majority of the non-TRs showed low methylation levels. In contrast, methylation levels of TR-CGIs are broadly distributed with a significant difference in their proportion among the sequences with 70% or more methylation (Fig. 2B and Supplementary Table S3). Specifically, when 30% methylation was used as a cutoff, there were 79 TR-CGIs out of 347 analyzed (22.77%) whereas there were only 8.55% non-TR CGIs, indicating a significant difference (p = 0.0062). Similar data were obtained in the case of 68 normal prefrontal cortex samples [22]. Analysis of 47 TR-CGIs with >40% methylation is shown in Fig. 2C. Nearly 60% of the methylated TR-CGIs showed very high levels of methylation (∼75-95%). Among the TR-CGIs with >30% methylation, there were 12 that showed methylation difference of at least 20% in the frontal cortices of different individuals, suggesting a degree of methylation variation in normal people. In contrast, ICRs are not reported to show any such variation among different individuals. Examples of a few TR-CGIs that showed methylation variation in the prefrontal cortex samples in different individuals are shown in Fig. 2C.

**Fig. 2.**
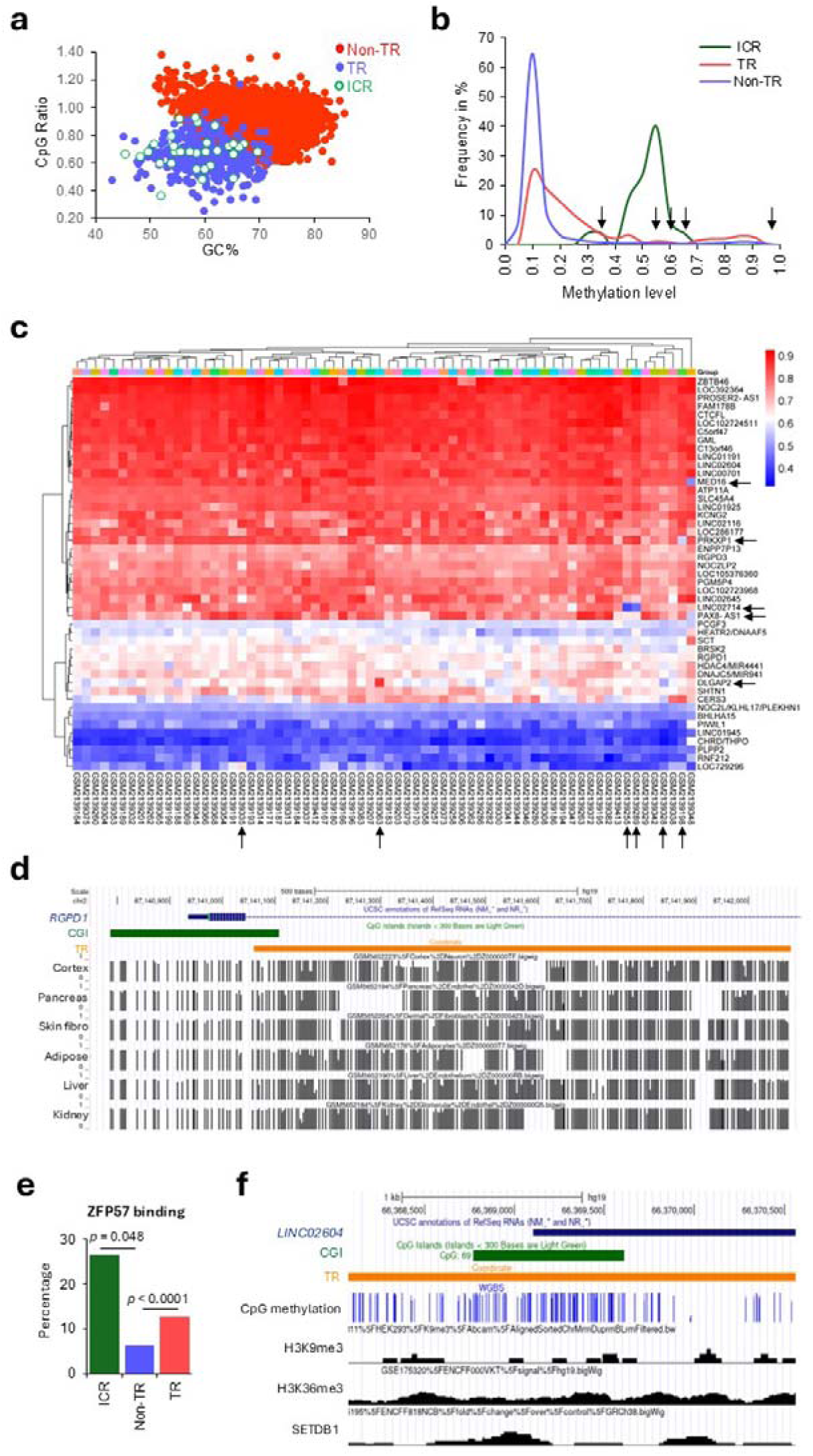
Sequence features, methylation levels and chromatin modifications associated with the TR-CGIs. (**A**) GC content and CpG ratio analyses of imprinting control regions (ICRs), TR-CGIs and non-TR & non-ICR CGIs (Non-TR). (**B**) Methylation levels (X-axis) of the three categories of CGIs in human cerebellum. Y-axis: percentages of CGIs at different methylation levels. 0.0 and 1.00 correspond to 0% and 100% methylation, respectively. Arrows indicate absence of significant difference in the proportion of TR-CGIs and non-TR CGIs with the indicated methylation values. (**C**) Heatmap of methylation levels of 47 TR-CGIs with 30% or more methylation in 68 frontal cortex specimens [22]. Horizontal arrows denote the sequences showing a difference in the level of methylation (± 20% from the average levels) whereas vertical arrows indicate the samples in which the difference in methylation level was observed. The heatmap was generated using average beta values of the indicated samples and genes employing the SR plot online tool [35]. (**D**) UCSC browser screenshot showing the methylation levels of the *RGPD1* 5’-most TR-CGI in six different tissues from different individuals. Sample identities are given above each track with black vertical lines. The height of each vertical line represents the methylation value (0.0-1.0) of the CpG site. **(E)** Analysis of ZFP57 binding sites in ICR, Non-TR and TR-CGIs. **(F)** UCSC browser screenshot of TR-CGI promoter of *LINC02604* showing DNA methylation levels from whole genome bisulfite sequencing and enrichment data of the indicated histone modifications and SETDB1 binding using HEK293 cells. Refer to Figure S4 for more detailed examples. Data are taken from publicly available datasets: <u>GSE200839</u>, <u>GSE175320</u>, <u>GSE175195</u>, <u>GSE80970</u>, <u>GSE129548</u>, <u>GSE247551</u>.

To investigate whether the TR-CGIs methylated in one tissue show similar levels of methylation in other tissues and to compare their levels of methylation in the CGI regions with TR regions, whole genome bisulfite sequencing (WGBS) data was obtained for different tissues from the publicly available databases. A set of 14 TR-CGIs that contain TRs at different locations relative to the CGIs were taken. We observed no drastic difference in the levels of methylation between the CGIs and the associated TRs (Fig. 2D and Supplementary Fig. S4). In addition, among the 14 TR-CGIs studied, there was no significant difference in the levels of methylation in the six tissues used for analysis.

### TR-CGIs, as in cases of ICRs, show a significant enrichment of ZFP57 binding sites

Previously published data suggested that ICR regions have a significantly preferential binding to proteins such as ZFP57, ZNF445 and SETDB1 [23,24]. Sequence data on ICRs, TR-CGIs and non-TRs were analyzed for the presence of binding sites of ZFP57 as consensus sequences for the latter two proteins are not available. One-fourth of the ICRs contain ZFP57 binding sites whereas the corresponding numbers were one in sixteen (∼6%) and one in eight for non-TR CGIs and TR-CGIs, respectively indicating a two-fold enrichment (p < 0.0001) in the latter (Fig. 2E). When enrichment for H3K9me3 and H3K36me3 modifications and binding sites of SETDB1 were investigated in 30 each of the TR-CGIs and non-TR CGIs and the 25 ICRs, a significant enrichment was observed in case of ICRs but there was no difference between the remaining two categories of sequences. Examples of TR-CGIs with their DNA methylation, histone modifications, and SETDB1 binding tracks are shown in Fig. 2F and Supplementary Fig. 1.

### Effects of reprogramming of differentiated tissues to iPSCs on methylation levels of TR-CGIs

An important characteristic of all ICRs is their exemption from epigenetic, i.e. methylation, reprogramming *in vitro* in iPSC generation and *in vivo* in preimplantation. To determine if TR-CGIs also share this feature, we analyzed methylation data on six different embryonic tissues (brain, muscle, skin, lung, kidney and pancreas) and two isogenic primed iPSC lines (iPSC1 and iPSC2 for each tissue) derived from them [25]. Between different embryos, there was no difference in the levels of methylation among the six tissues for ICRs, but there was a significantly lower variation for TR-CGIs than non-TR CGIs (p = 0.0069) (Fig. 3A). This difference between TR-and non-TR CGIs could be because of tissue-specific differences in methylation in the latter category. Although there was a higher degree of variation between different iPSCs, the difference was significant in the case of non-TR CGIs and ICRs, but not the TR-CGIs. The effects of reprogramming accompanying the iPSC generation was particularly evident in the case of ICRs when the proportion of sequences was plotted against the methylation levels (Fig. 3B). PCA analysis of the three categories of sequences revealed the effects of reprogramming on the methylation levels and that the two iPSC lines derived from the corresponding tissues (iPSC1 and iPSC2) showed the higher degree of variation than the tissues (Fig. 3C-E).

**Fig. 3.**
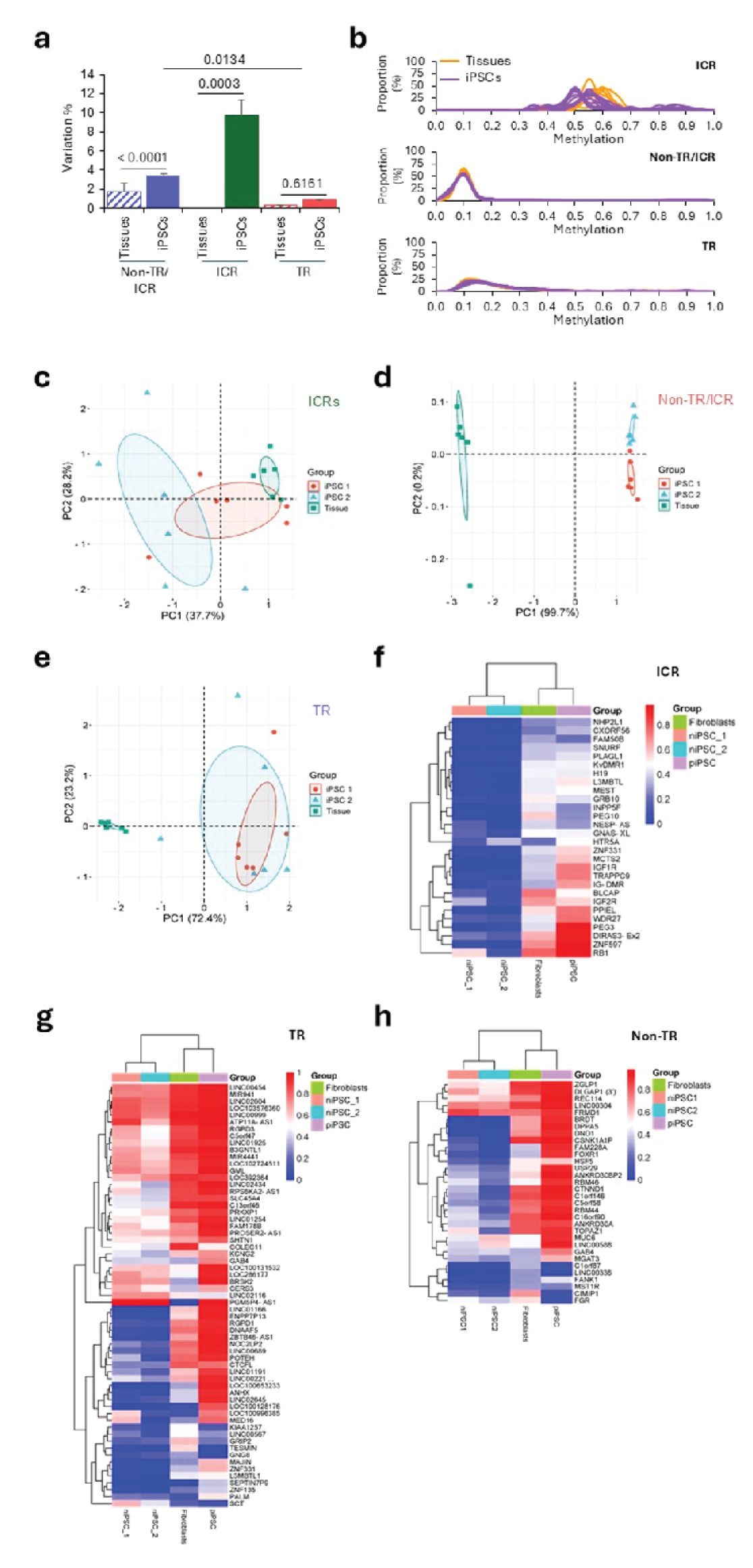
Effects of *in vitro* reprogramming on methylation levels of TR-CGIs of terminally differentiated tissues from the same embryo and their corresponding iPSCs. (**A**) Proportion of sequences showing differences (Variation %) in methylation in the DNAs from different tissues of the same embryo and their corresponding iPSCs. Methylation values of the tissues were used to calculate variation based on ≥ 20% difference in the levels of methylation for each tissue from the average value. **(B)** Proportion of sequences in percentage with different levels of methylation before (orange lines) and after reprogramming (purple lines). **(C-E)** PCA analysis of ICRs, TR-CGIs and Non-TR CGIs based on their methylation levels in the multiple tissues and their iPSCs. The dots represent different samples. **(F-H)** Hierarchical clustering of sequences showing ≥ 30% methylation in any of the fibroblasts or primed or naïve iPSCs derived from them. The heatmaps were generated using average beta values of the indicated samples and genes employing the SR plot online tool [35]. Data are taken from publicly available datasets: <u>GSE200834, GSE76640, GSE76970, GSE110366.</u>

A second important feature reported in case of methylation levels for ICRs during *in vitro* reprogramming was their maintenance of methylation in primed, but not the naïve iPSCs [26,27]. When data on fibroblasts and primed as well as naïve iPSCs derived from them were analyzed by hierarchical clustering, there was an obvious similarity between the fibroblasts and primed iPSCs whereas the naïve iPSCs formed a separate cluster (Fig. 3F). A similar pattern was observed for TR-CGIs (Fig. 3G; Supplementary Table S5). Thus, the TR-CGIs are similar to the ICRs by their low-level variation among the different tissues of the same individual (embryo), and more similar methylation patterns between the tissues and the primed iPSCs derived from them.

We used these same fibroblasts, primed-and naïve iPSCs, and a different approach to compare TR-CGIs and non-TR CGIs. 200 non-TR CGIs overlapping TSSs were randomly chosen from the human genome. Five out of the 200 have >30% methylation, a much lower frequency than that of TR-CGIs, and thus a low number to compare with the methylated TR-CGI set. To increase the yield of methylated non-TR CGIs, we first identified genes containing CGIs with >30% methylation in the human cerebrum and cerebellar samples described above, and then identified 32 non-TR CGIs with >30% methylation in fibroblasts (Supplementary Table S9). 28 out of the 32 retained their methylation in primed iPSCs (Fig. 3H), a feature of ICRs and TR-CGIs described above. Interestingly, 12 of these 28 were closely linked (within 2.0 Mb, the lower limit of TR-CGI density within clusters) to a TR-CGI (many of these in TR-CGI clusters). Whether the proximity to TR-CGIs has any influence on the methylation of the non-TR CGIs is not known.

### Spermatogenesis specifically effects the methylation levels of ICRs and TR-CGIs

An additional approach to assessing the stability of methylation among the three classes of sequences is to compare the data on gametes and somatic tissues from the same individual. This is because gametogenesis is one form of *in vivo* reprogramming wherein demethylation in the primordial germ cell genome is followed by the establishment of gamete-specific methylation patterns. To enable these analyses, methylation data on five males whose DNAs from somatic cells (blood and saliva) and semen were used (Fig. 4A-E). Average methylation values of blood and saliva from each person were used as reference for identifying sequences with > 20% difference in the methylation levels in both the somatic tissues and semen of the same individual. The data suggest that non-TR CGIs show significantly higher variation in somatic tissues than TR-CGIs or ICRs. There was no difference in the proportion of sequences in this category showing variation in methylation levels between semen and other tissues however, significant differences between the two categories of cell types were observed in case of TR-CGIs and ICRs (Fig. 4A-B). The closer relatedness between the somatic tissues over their distant similarity to semen was also obvious in the PCA analysis of the three categories of sequences.

**Fig. 4.**
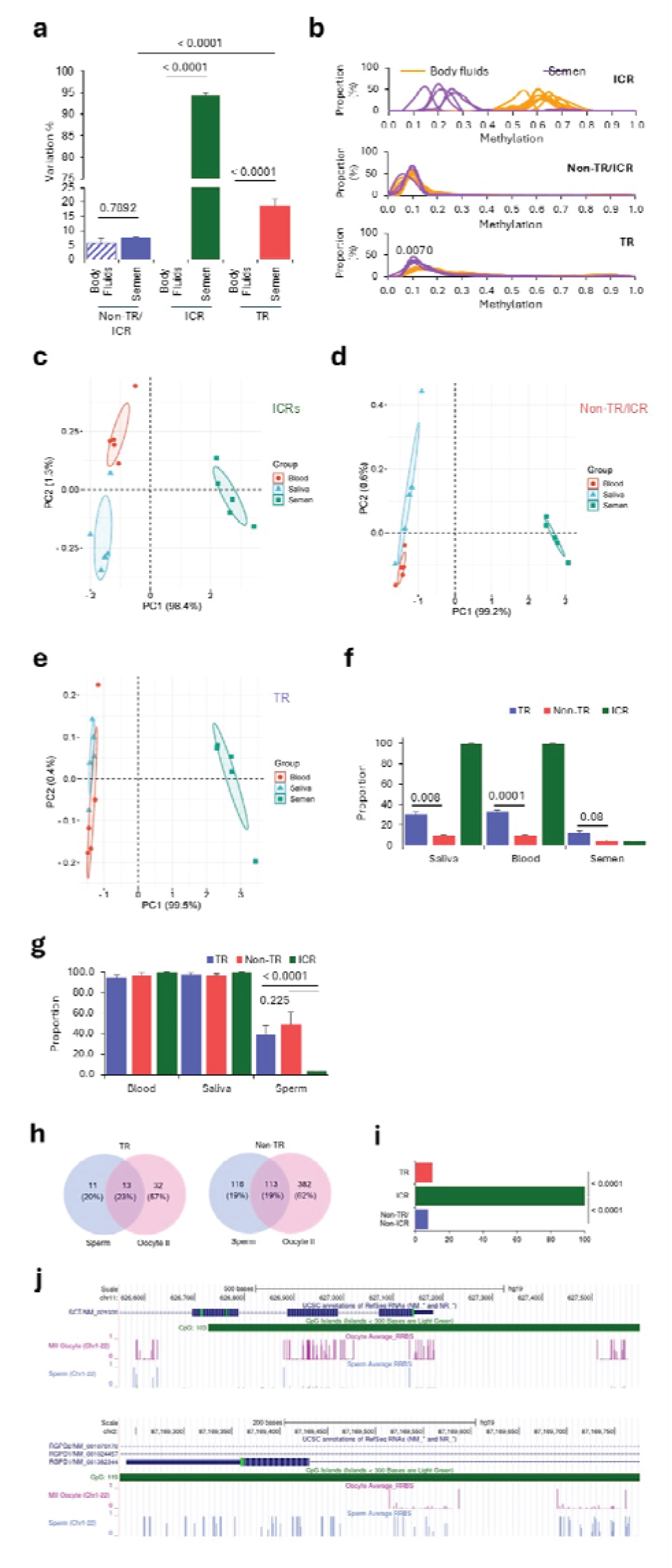
Effects of spermatogenesis-associated reprogramming on the methylation levels of TR-CGIs. **(A)** Proportion of sequences of the three categories mentioned on the X-axis showing differences in methylation levels between the same samples from five different individuals. Average methylation levels in the blood and saliva (body fluids) are used as reference. **(B)** Proportion of sequences in percentage with different levels of methylation before (orange) and after reprogramming (purple). **(C-E)** PCA analyses of the three different categories of CGIs based on their methylation levels in blood, saliva and semen. **(F)** Proportion of sequences (in percentage on Y-axis) showing methylation levels of ≥ 30% in saliva, blood and semen. **(G)** Proportion of methylated sequences with ≥ 30% levels in saliva, blood and semen. (H) Venn diagrams showing the number of TR-CGI and non-TR sequences with ≥ 30% methylation in sperm and MII oocytes. **(I)** Proportion of sequences (in percentage) showing differences in methylation levels between the sperm and the MII oocytes. **(J)** Examples of TR-CGIs showing high levels of methylation in the MII oocytes and low levels in sperm (top panel); low levels in the MII oocytes and high levels in the sperm (bottom panel). Refer to Figure S1 for more detailed information. Data are taken from publicly available datasets: <u>GSE49828</u>, <u>GSE51239</u>.

### Sperm and MII oocytes share some methylated TR-CGIs and non-TR CGIs

A comparison of methylation levels of TR-CGIs and non-TR CGIs between oocytes and sperm would shed light on the fraction of shared and unique sets of sequence. For this purpose, methylation data on sperm and oocyte were used to determine the proportion of sequences that show > 30% methylation in oocytes only, sperm only or both (Fig. 4H-4J). The fraction of sequences showing differences in methylation levels in the gametes was, as expected, highest in case of ICRs and was nearly similar in TR-CGIs (∼10%) than the non-TR CGIs (∼8%). (p = 0.8056) (Fig.4H). Overall, ∼60% and ∼20% of the TR-CGIs with ≥ 30% methylation showed sperm-and oocyte-specific methylation, respectively whereas the remainder were methylated in both types of gametes (Fig. 4I). Similar results were obtained with non-TR CGIs. An example each of the TR-CGIs with increased methylation in oocyte relative to sperm and *vice versa* is shown in Fig. 4J and Supplementary Figure S6.

### Methylation of TR-CGIs in preimplantation

Preimplantation development, i.e., the progression of the zygote to the blastocyst is another phase of reprogramming that naturally occurs in eutherian mammals, a phase in which there is genome-wide loss of methylation by active as well as passive processes. However, ICRs are resistant to this reprogramming process and the methylated states established in the gametes are maintained in preimplantation [28,29]. TR-CGI genes are not among lists of known imprinted genes, so we do not expect that TR-CGI methylation is inherited from gametes. Specifically, we expect TR-CGIs to be unmethylated in gametes, or if methylated in one or both gametes, unmethylated in one or more preimplantation stages. To examine the methylation levels of the TR-CGIs, the same subset used to comparatively study the CGIs and the associated TRs (Fig. 2D) was studied. Analysis of stages from zygote to the recently implanted embryo showed varying methylation patterns for different TR-CGIs (Fig.5). In one category (Fig. 5A), high methylation levels persisted throughout all stages whereas in a second category there was an overall loss of methylation till the blastocyst stage, but a gain in methylation after implantation (Fig. 5B). In the third category, the methylation levels were moderate in preimplantation stages and after implantation there was complete methylation (Fig. 5C). The fourth category of TR-CGIs had low level methylation even after implantation, but became methylated to different levels in different tissues (Fig 5D-5E). Time courses of developmental TR-CGI methylation such as the Fig. 5A example leave open the possibility that these TR-CGIs inherit their methylation.

**Fig. 5.**
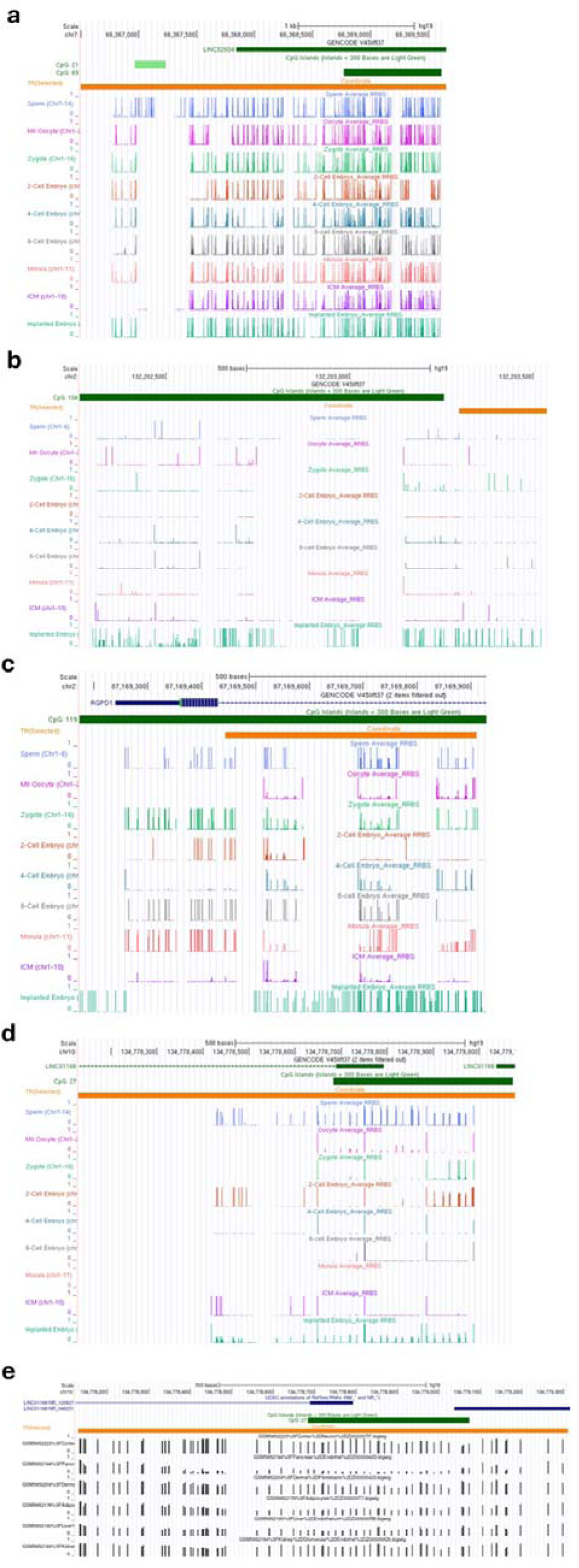
Methylation levels of TR-CGIs during preimplantation development. **(A)** *LINC02064*-associated TR-CGI showing high levels of methylation throughout preimplantation. **(B-E)** Example of a sequence showing low levels of methylation during preimplantation but high levels after implantation (B; NOC2LP2), moderate levels of methylation but high after implantation (C), low levels of methylation in preimplantation as well as peri implantation in embryonic development (D), but gain methylation in some tissues during subsequent stages of development (E). Data are taken from publicly available datasets: <u>GSE49828</u>, <u>GSE51239</u>.

### A subset of TR-CGI associated gene transcription correlates with the level of TR-CGI methylation

As described earlier, a small number of TR-CGIs showed inter-individual differences in the levels of methylation in frontal cortex (Fig. 2C). These results prompted us to investigate whether there is an association of the methylation differences with transcript levels for the TR-CGI-associated genes. For this purpose, we took advantage of the observation that TR-CGIs showed differences in methylation levels between the naïve and primed iPSCs. Here, we used UCLA1, a primed human embryonic stem cell line and its naïve derivative obtained after 5iLAF treatment. Both transcript and methylation data were available for 23 genes wherein associated TR-CGIs showed > 20% difference between the two cell types (Supplementary Table S6). Of these, five genes showed significant correlation between the methylation and transcript levels (Fig. 6). In the case of *BRSK2*, there was a positive correlation, whereas in the case of *RGPD1*, *NOC2L*, *RGPD3*, and *RGPD4*, there was a negative correlation. A loss of TR-CGI methylation may directly cause the transcription increase, or alternatively, the increase may be due to genome-wide methylation loss.

**Fig. 6.**
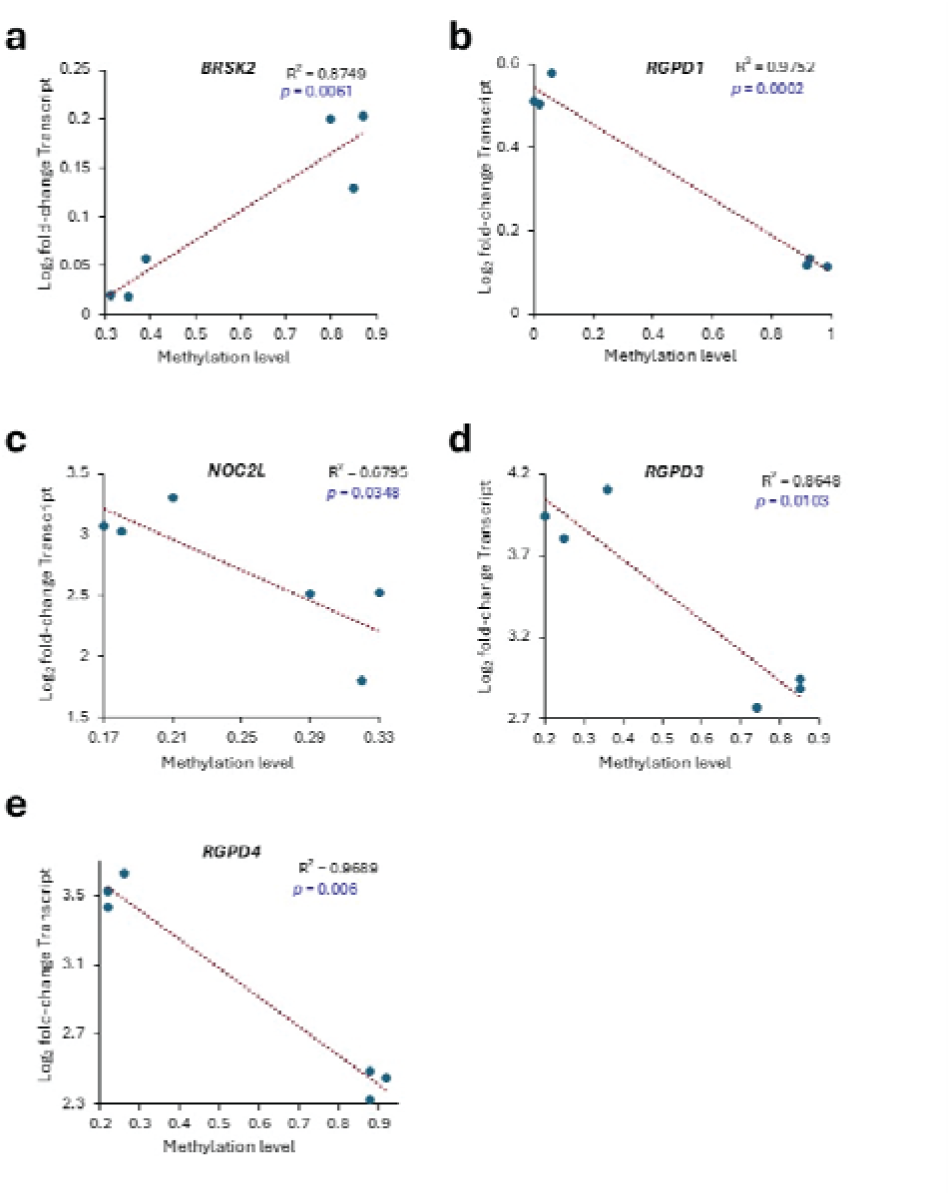
Relationship between the levels of methylation of TR-CGI promoters and their transcripts. (A-E) Correlation plots showing methylation levels on the X-axis and expression levels on the Y-axis. Data was taken from three samples each of naïve and primed hESCs. Data are taken from publicly available dataset: <u>GSE76970</u>.

### Acquisition of tandem repeats accompanied human evolution with evidence of recent additions

We compared orthologous sequences among primates and mouse to gain insights into the origins of the identified human TR-CGIs (Fig. 7A, and Supplementary Table S1). Whereas some sequences (e.g., *TOAK3* TR-CGI) acquired tandem repeats at the evolutionary divergence of mouse and primates, many (e.g., *AXIN1*) were present after the divergence of lower primates or strepsirhines (e.g., lemurs) and higher primates or platyrhines (e.g., New and Old World monkeys). Although, lesser in number, there was evidence of gain of the TRs at the divergence of New World and Old World monkeys (e.g., *GEMIN4*) as well as at hominid and non-hominid divergence ∼29.6 million years ago (e.g., *PRKXP1*) [30].

**Fig. 7.**
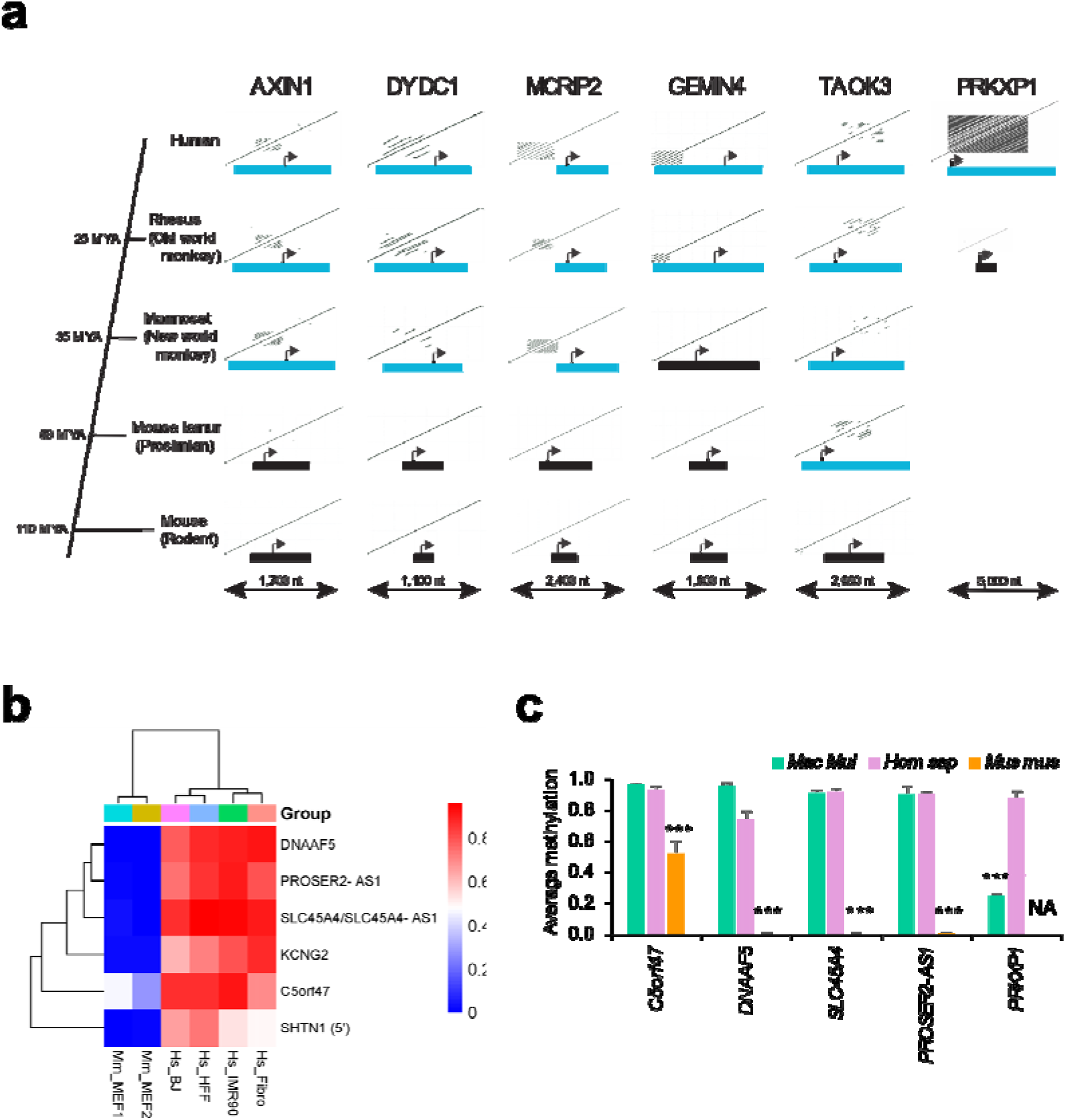
Evolutionary origins of TRs in TR-CGI genes. **(A)** For each orthologous gene, dot plots of the CGI-promoters and surrounding sequences from the five species are displayed. Blue rectangles are TR-CGIs and black rectangles are CGIs without TRs. Arrows are transcriptional start sites and directions of transcription. **(B)** Heatmap of methylation levels of orthologous genes in human (Hs) and mouse (Mm) fibroblasts. The corresponding genes in mouse lack tandem repeats. The heatmap was generated using average beta values of the indicated samples and genes employing the SR plot online tool [25]. **(C)** Comparisons of methylation profiles of orthologous genes in human (*Hom sap*), rhesus macaque (*Mac mul*) and mouse (*Mus mus*). *** indicates *p* values < 0.0001 estimated by Fisher’s paired *t*-test. NA: Not applicable; ortholog absent in mouse. Data are taken from publicly available datasets: <u>GSE124708</u>, <u>GSE110366</u>, <u>GSE233417</u>, <u>GSE120137</u>.

To relate whether the presence of tandem repeats increases the methylation levels in the TR-CGIs, methylation levels of a few orthologous genes were studied in four different human fibroblast lines and two mouse embryonic fibroblast lines. Hierarchical clustering analysis clearly segregated the mouse cell lines from the humans on the basis of higher methylation levels in the latter (Fig. 7B). As a second line of evidence, methylation data from adipose tissues on five orthologs of human, Rhesus macaque, and mouse were examined (Fig. 7C) [31–33]. Of these, four (*C5ORF47, DNAAF5, SLC45A4*, and *PROSER2-AS1*) have TRs in both human and Rhesus macaque but not in mouse. GC content and CpG ratio analysis showed a rough segregation of mouse CGIs from human and rhesus CGIs (Supplementary Fig. S9). These sequences showed higher levels of methylation in the two primates than the rodent. Importantly, *PRKXP1* that has no ortholog in the mouse is devoid of TRs in the Rhesus macaque and showed significantly lower methylation levels than in humans. The human *PRKXP1* TR-CGI was methylated in all somatic tissues examined (Supplementary Fig. S9). Further analysis revealed the association of TRs with the *PRKXP1* promoter CGIs of chimpanzee and gorilla, much shorter length (300 bp) in orangutan and their absence in gibbon and the African green monkey.

## Discussion

The identification and analyses of TR-CGI promoters revealed certain similarities and differences with ICRs. The occurrence of TRs itself, clustering of the genes possessing the TR-CGI promoters, GC-contents, CpG ratios, methylated states in multiple tissues, enrichment of ZNF57 binding sites and substantial effects on methylation due to spermatogenesis-associated reprogramming are some of the common features. Some distinctive features include broad ranges of methylation levels observed, variation in methylation levels of a small subset of the TR-CGIs in the same tissues between individuals, tissue-specific differences in methylation, loss of methylation in both types of gametogenesis, gain of methylation after implantation and lack of significant enrichment of H3K9me3 and H3K36me3 (SETDB1-binding sites).

Based on the methylation patterns observed in a sample of TR-CGIs in preimplantation stages, gametes and multiple adult tissues, we postulate that TR-CGI methylation is mainly acquired post-fertilization and can be stably maintained for the entire lifespan of the individual. These methylation patterns are preserved in case of primed iPSCs but not naïve iPSCs, suggesting their lack of sensitivity to the former type of *in vitro* reprogramming. Given the evidence that methylation levels can influence gene expression in five out of the 23 genes studied, the association of primate-specific acquisition of TRs and their increased potential of attaining methylated states that are more evident after implantation suggest that the TR-CGIs also play a role in the hypothetical epigenetic bifurcation events creating canals in the epigenetic landscape of development proposed by Waddington [15,34].

As mentioned above, the common structural feature associated with the 365 genes with TR-CGI promoters and ICRs is the presence of tandem repeats that do not share sequences. It appears that the repetitive nature is more likely an explanation for the ability to acquire methylation. Since methylation of certain TR-CGI promoters is associated with significant alterations in gene expression, it is reasonable to expect that there are some commonly shared functional features with the imprinted ICRs. However, a unique feature of ICRs is their invariant difference in the methylation levels between the male and female gametes of eutherian mammals. In the case of the TR-CGIs, only three out of the 15 sequences tested showed such difference. However, methylation data on more samples or analysis in families is needed to establish whether this gamete-specific difference or parental allele-specific methylation persists.

An interesting feature observed for TRs among the TR-CGIs identified is their gradual appearance at evolutionary branch points that separate strepsirhines from platyrhines, old world from the new world monkeys and hominids from non-hominids. For example, even among the great apes *PRKXP1* CGI gained TRs in gorilla (1,100 bp), chimpanzee (2,200 bp) and humans (4,400 bp), but not in the more distant gibbon. In orangutan, there is an indication of the presence of a TR, but the length (300 bp) did not meet the threshold value of 400 bp to be included. In support of the hypothesis that the TRs have a role in imparting methylation to the CGIs they are associated with, methylation levels of the *PRKXP1* CGI in humans are much higher than rhesus wherein the ortholog is devoid of TRs. In this context, this hypothesis needs to be tested by a careful examination of the TR-CGIs that have acquired TRs recently in the hominid evolution among the great apes.

In summary, the TR-CGI – associated genes are a unique subset in the human genome with distinctive properties that are similar in some respects but different in others from the ICRs and appear to have a functional role on gene regulation. More detailed analyses are needed to establish their evolutionary significance, if any.

## Materials and Methods

### Identification of TR-CGIs

Promoter-proximal CGIs were identified from the annotated CGI track in the UCSC browser of hg19 and hg38 assemblies using visual inspection of dotplots. In case of an annotation in just one track, or different CGI lengths in the two tracks, the sole or longer CGI was used to deduce the coordinates of the other track ‘s CGI via sequence comparisons. The BLASTN (default) variables for generating dotplots that are then scored for presence of tandem repeats in hg19 and hg38 assemblies: somewhat similar sequences; expect threshold 0.05; word size 11; match/mismatch score 2,-3; existence: 5 extension: 2. TRs within 200 bp of promoter-proximal CGIs ranged in size from <100bp to ≥400bp. We chose to limit our studies to the promoter-proximal TRs ≥400bp (defined as TR-CGIs) because of the length similarities to ICRs (example shown in Figure 1B).

### Determining methylation levels in TR-CGI, ICR and non-TR & non-ICR sequences

CGI methylation values as percents or fractions were calculated from human 450K and EPIC Illumia microarrays, reduced representation bisulfite sequencing (RRBS), and whole genome bisulfite sequencing (WGBS) datasets using annotated CGI coordinates (see above for details concerning TR-CGI coordinates). For comparisons, coordinates for human imprinting control regions (ICRs; ref. 27) and for the remaining (non-TR-CGI & non-ICR -associated CGIs), coordinates were derived from the manifest files of the 450K and EPIC array manifest files.

### Estimation of the proportions of sequences with different methylation levels (Fig2B)

Fishers exact two-tailed *t*-test was performed to determine in the differences among the TR-and non-TR CGIs, and ICRs. In each case, the proportion of sequences with a specific range of methylation values were taken and the reminder as the proportion that have values different from the specific range. For example, the proportion of non-TR CGIs with methylation levels between 0 and 0.5 was ∼8% whereas in case of TR-CGIs, it was ∼1.0%. This would give ∼92% and ∼99% as proportions with methylation levels outside this range. The *p* value of this difference was 0.035 and taken as significant.

### Evolutionary appearance of TRs in CGI

Promoter-proximal CGI sequences from non-human primate and rodent species, corresponding to human TR-CGIs, were studied to determine the latest evolutionary appearance of each human TR-CGI. Given the limited number of available annotated vertebrate genome sequences, we approximated latest evolutionary appearance to all primates, new-world monkeys (NW), old-world monkeys (OW), apes or humans.

## Author contributions

Conceptualization: KNM, JRC

Design of the work: KNM, LK, JRC

Acquisition of data: KNM, AA, LK, JRC

Analysis of data: KNM, LK, JRC

Interpretation of data: KNM, LK, JRC

Writing – original draft: KNM, JRC

## Additional Information

Authors declare that they have no competing interests.

## Data and materials availability

Data is provided within the manuscript or supplementary information files. The following publicly available GEO DataSets were analyzed: <u>GSE49828</u>, <u>GSE51239, GSE76640,</u> <u>GSE76970, GSE80970, GSE110366</u>, <u>GSE120137, GSE124708, GSE129548</u>, <u>GSE175195</u>, <u>GSE175320, GSE200834, GSE200839</u>, <u>GSE233417</u>, <u>GSE247551</u>.

## Supplementary Materials

Supplementary Tables S1 to S9

Supplementary Figures S1 to S9

