## Supplementary Figures for "A new class of human CpG island promoters with primate-specific repeats"

**Supplementary Information: Supplementary Figures**

**Title:** Human CpG island methylation that is resistant to iPSC reprogramming accompanies evolutionary acquisition of tandem repeat sequences

**Authors:** K Naga Mohan[1,3], Anuhya Anne[1], Lov Kumar[2], J Richard Chaillet[3]

**Supplementary Figure Legends**

**Suppl. Fig S1. Bioinformatic analysis of non-TR/ICR CGI (non-TR) promoter sequences. (A)** Disease ontology analysis. **(B)** Pathway analysis. **(C)** Analysis of biological processes. **(D)** GTex analysis.

**Suppl. Fig. S2. Dotplots of 14 human TR-CGIs with hg19 chromosome coordinates of CGI and TR sequences and alignments of two TR copies.**

**Suppl. Fig. S3. Comparison of sequence features of ICR, TR and non-TR CGI sequences. (A)** Abundance of sequences in percentage on (Y-axis) showing different methylation levels in percentage (X-axis). **(B)** Abundance of sequences in percentage (Y-axis) showing different CpG ratios (X-axis).

**Suppl. Fig. S4. UCSC screenshots showing methylation levels of 14 selected TR-CGIs.** In each panel, sample identities are given above track.

**Suppl. Fig. S5. UCSC screenshots of methylation levels and enrichment of H3K9me3 and H3K36me3 and SETDB1 binding sies for a few selected TR- and non-TR CGIs in HEK 293 cells.** In eachpanel, sample identities are given above the track.

**Suppl. Fig. S6. UCSC screenshots of methylation levels of 14 selected TR-CGIs (as in Suppl. Fig. 3.) in spermatocytes (blue horizontal bars) and MII oocytes (pink horizontal bars).**

**Suppl. Fig. S7. UCSC screenshots of methylation levels of 14 selected TR-CGUs (as in Suppl. Fig. 3.) in different stages of preimplantation development and after post-implantation.**

**Suppl. Fig. S8. Mouse and human dotplots accompanying Fig. 7B**

**Suppl. Fig. S9. GC content & CpG ratio (left) and tissue methylation (right) of CGIs shown in Fig. 7B and 7C.**

Suppl. Fig. 1

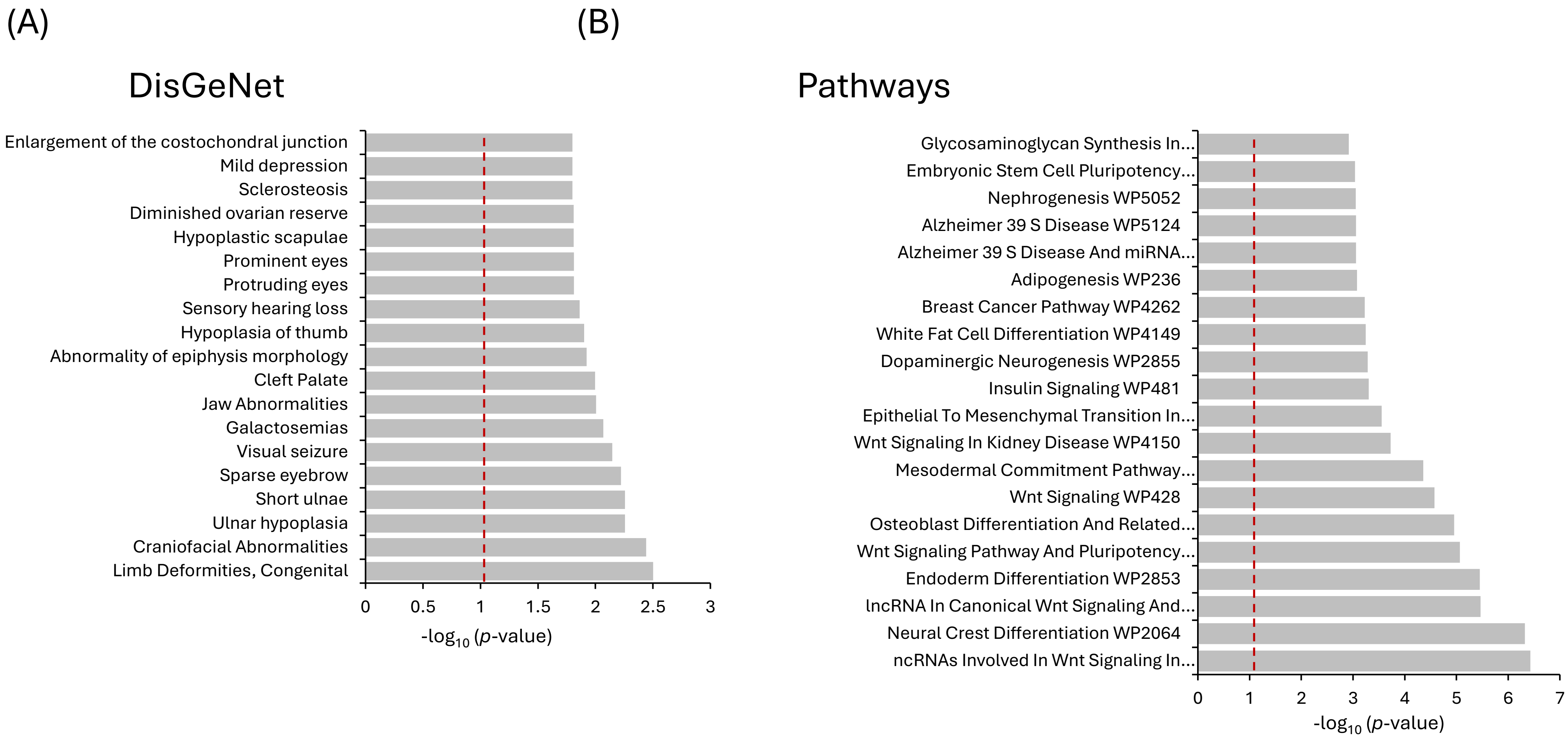
(A)
DisGeNet
Enlargement of the costochondral junction
Mild depression
Sclerosteosis
Diminished ovarian reserve
Hypoplastic scapulae
Prominent eyes
Protruding eyes
Sensory hearing loss
Hypoplasia of thumb
Abnormality of epiphysis morphology
Cleft Palate
Jaw Abnormalities
Galactosemias
Visual seizure
Sparse eyebrow
Short ulnae
Ulnar hypoplasia
Craniofacial Abnormalities
Limb Deformities, Congenital
0
0.5
1
1.5
2
2.5
3
-log10 (p-value)
(B)
Pathways
Glycosaminoglycan Synthesis In...
Embryonic Stem Cell Pluripotency...
Nephrogenesis WP5052
Alzheimer 39 S Disease WP5124
Alzheimer 39 S Disease And miRNA...
Adipogenesis WP236
Breast Cancer Pathway WP4262
White Fat Cell Differentiation WP4149
Dopaminergic Neurogenesis WP2855
Insulin Signaling WP481
Epithelial To Mesenchymal Transition In...
Wnt Signaling In Kidney Disease WP4150
Mesodermal Commitment Pathway...
Wnt Signaling WP428
Osteoblast Differentiation And Related...
Wnt Signaling Pathway And Pluripotency...
Endoderm Differentiation WP2853
lncRNA In Canonical Wnt Signaling And...
Neural Crest Differentiation WP2064
ncRNAs Involved In Wnt Signaling In...
0
1
2
3
4
5
6
7
-log10 (p-value)

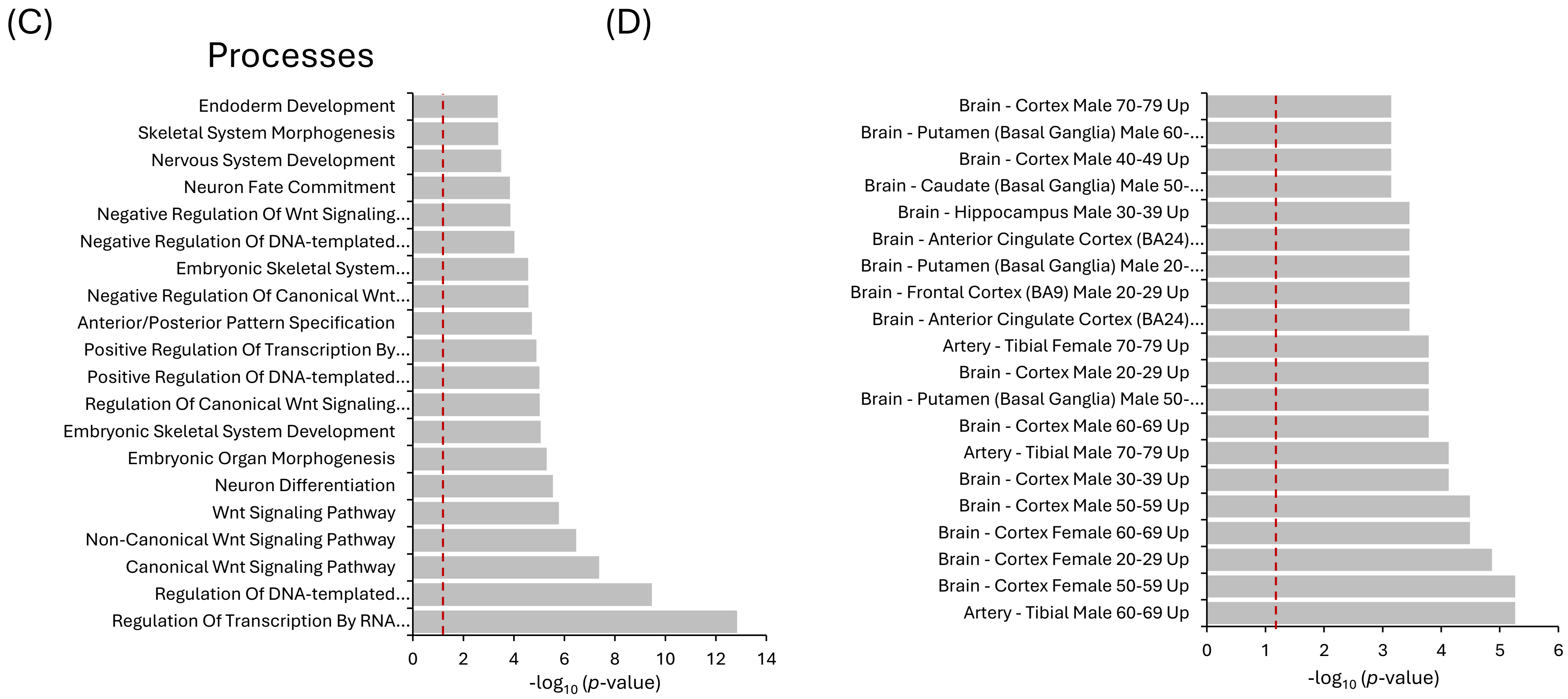

(C)
Processes
Endoderm Development
Skeletal System Morphogenesis
Nervous System Development
Neuron Fate Commitment
Negative Regulation Of Wnt Signaling...
Negative Regulation Of DNA-templated...
Embryonic Skeletal System...
Negative Regulation Of Canonical Wnt...
Anterior/Posterior Pattern Specification
Positive Regulation Of Transcription By...
Positive Regulation Of DNA-templated...
Regulation Of Canonical Wnt Signaling...
Embryonic Skeletal System Development
Embryonic Organ Morphogenesis
Neuron Differentiation
Wnt Signaling Pathway
Non-Canonical Wnt Signaling Pathway
Canonical Wnt Signaling Pathway
Regulation Of DNA-templated...
Regulation Of Transcription By RNA...
0 2 4 6 8 10 12 14
-log10 (p-value)
(D)
Brain - Cortex Male 70-79 Up
Brain - Putamen (Basal Ganglia) Male 60-...
Brain - Cortex Male 40-49 Up
Brain - Caudate (Basal Ganglia) Male 50-...
Brain - Hippocampus Male 30-39 Up
Brain - Anterior Cingulate Cortex (BA24)...
Brain - Putamen (Basal Ganglia) Male 20-...
Brain - Frontal Cortex (BA9) Male 20-29 Up
Brain - Anterior Cingulate Cortex (BA24)...
Artery - Tibial Female 70-79 Up
Brain - Cortex Male 20-29 Up
Brain - Putamen (Basal Ganglia) Male 50-...
Brain - Cortex Male 60-69 Up
Artery - Tibial Male 70-79 Up
Brain - Cortex Male 30-39 Up
Brain - Cortex Male 50-59 Up
Brain - Cortex Female 60-69 Up
Brain - Cortex Female 20-29 Up
Brain - Cortex Female 50-59 Up
Artery - Tibial Male 60-69 Up
0 1 2 3 4 5 6
-log10 (p-value)

Suppl. Fig. 2

### ATP11A-AS1

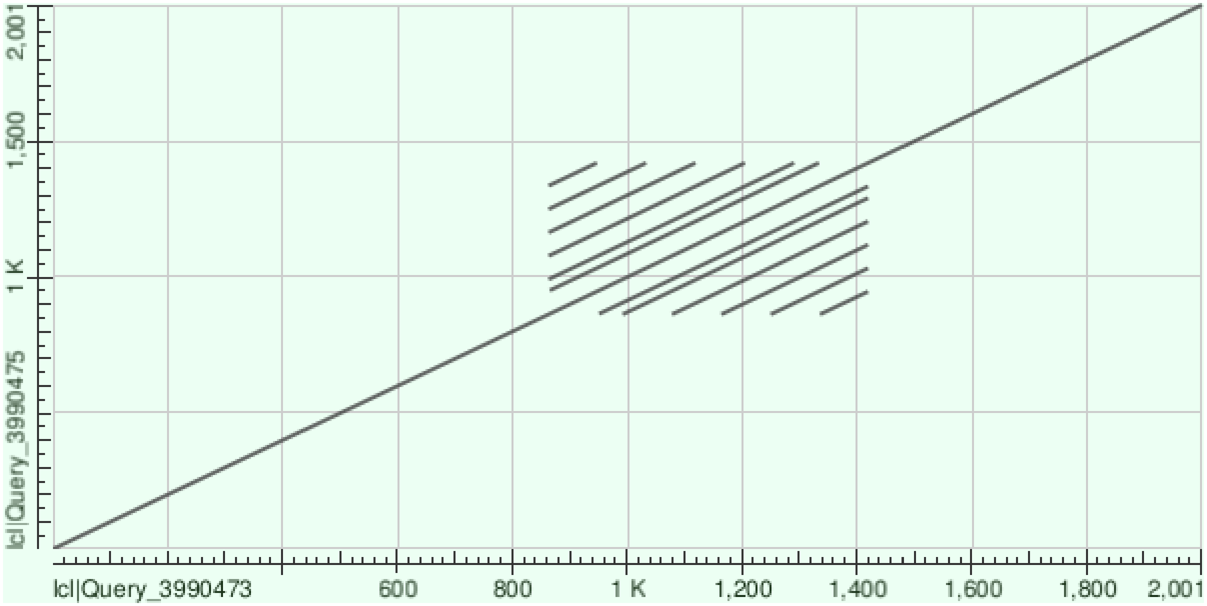


**CGI: Chr13:113,410,005-113,410,362**

**Alignment within TR (Chr13:113,409,860-113,410,420): 85% identity**

```
Query  4    CCAGAACCATTTCCAGTCACGGAACCGTCACTCAGAGCAGCTCCCAGAACCATTTCCAAT  63
            |||||||||||||| ||||||||| || |||||||||| ||||||||||||||||||  |
Sbjct  391  CCAGAACCATTTCCGGTCACGGAAGCGGCACTCAGAGCGGCTCCCAGAACCATTTCCGGT  450

Query  64   CATGGAAGCGTCACTCAGAGCGGCTCCTAGAACCATTTCCAGTCAGGGAAGCATCAATCA  123
            || |||| || |||||||||||||||| |||||||||||| |||| |||||| ||| |||
Sbjct  451  CACGGAAACGGCACTCAGAGCGGCTCCCAGAACCATTTCCGGTCACGGAAGCGTCACTCA  510

Query  124  GAGCGGCTCCCAGAACCATTTCCGGTCACGGAACCATCACTCAGAGCGGC  173
            |||| | | | |||| ||||||| |||| |||| ||| ||| ||||| ||
Sbjct  511  GAGCAGATTCTAGAATCATTTCCAGTCATGGAAGCATTACTTAGAGCAGC  560
```

### C5orf47

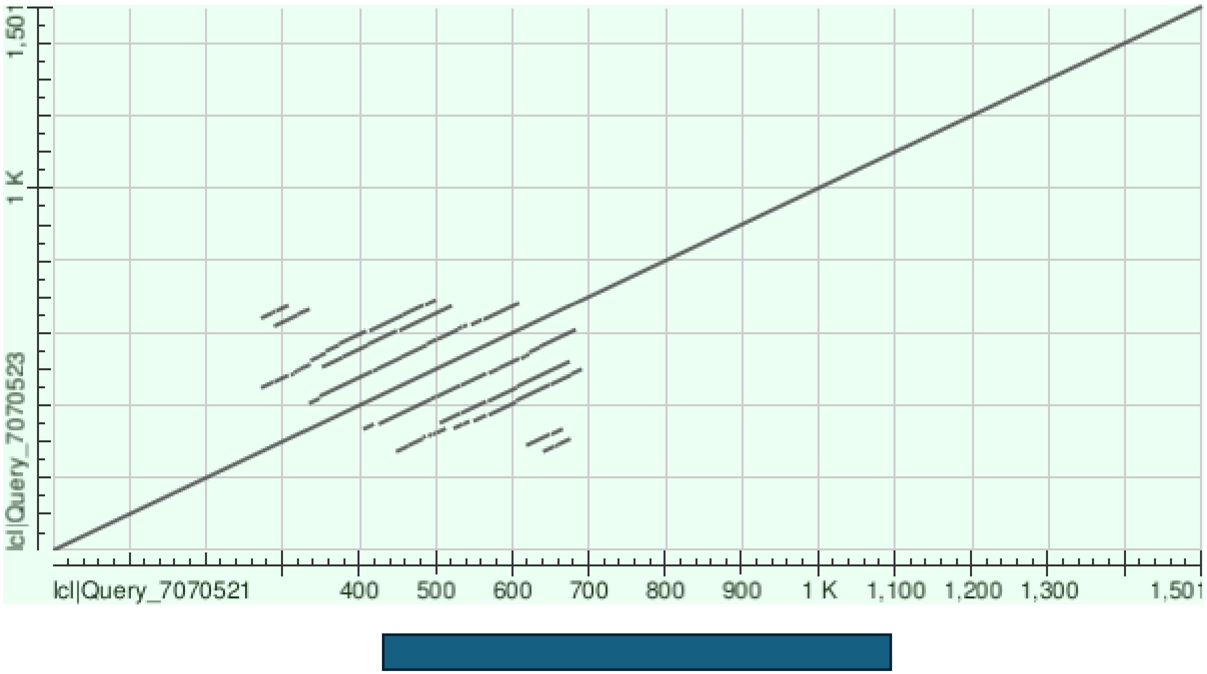


CGI:Chr5:173,415,925-173,416,586

**Alignment within TR (Chr5:173,415,780-173,416,200): 77% identity**

```
Query  74   TGATCCGGCCTTAACCCCGT---CACCCCATCCCTACTGGGTGGTCCTGCCCTAACCCCA  130
            || ||||||| ||||||  |   | ||||||||| || ||||||||  | |||||||||
Sbjct  267  TGGTCCGGCCCTAACCCTTTGTCCCCCCCATCCCCACAGGGTGGTCTGGTCCTAACCCC-  325

Query  131  CCACCCCCATCCCTGCGAGGTGGTCCGGCCGTAACCCCCTCCCCCCA-TCCCCACGCGGT  189
               |||| ||||   || ||||||| |||| |||| ||||||||||| ||||| |  |||
Sbjct  326  ---CCCCAATCCTCACGCGGTGGTCTGGCCCTAACTCCCTCCCCCCAATCCCCGCAGGGT  382

Query  190  GGTCTGGCCCTAACCCTGTCACCGCATCCCT  220
            |||||||||||||||  |   ||||||||||
Sbjct  383  GGTCTGGCCCTAACC--GCTCCCGCATCCCT  411
```

#### FAM178B

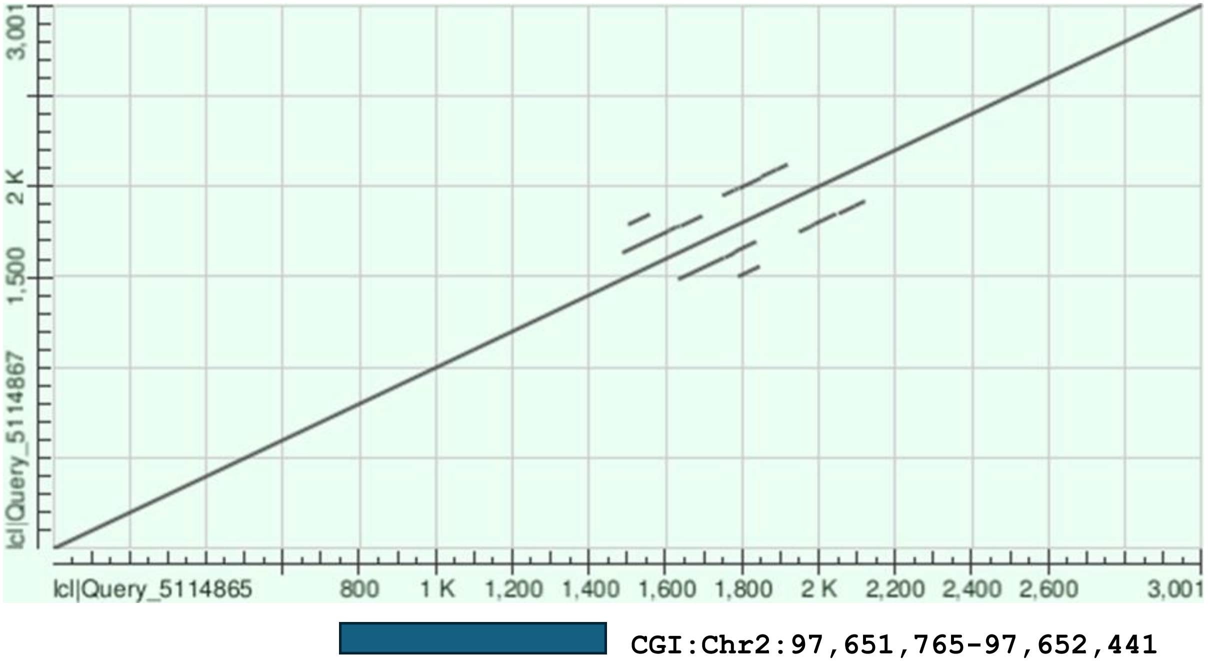


CGI:Chr2:97,651,765-97,652,441

**Alignment within TR (Chr2:97,652,485-97,653,120): 77% identity**

```
Query  3    GGCGCGGTGTCTCATGCCTATAATCCCAGCACTTTGGGAGGCTGAGACGGGACGATTGCT  62
            ||||||||| |||| |||| |||||||||||||||||||||| ||| ||||   ||||| 
Sbjct  149  GGCGCGGTGGCTCACGCCTGTAATCCCAGCACTTTGGGAGGCCGAGGCGGGCGAATTGCC  208

Query  63   TGAGCCCAGGAGTGAGAGACCTGCCTGGGCGACATGGCGAAACCCCGTCTCTACTAAAAA  122
            ||||  |||||||   ||||| | |||| | |||||| ||| ||| ||||||||||||||
Sbjct  209  TGAGGTCAGGAGTTCCAGACCAGTCTGGCCAACATGGTGAAGCCCTGTCTCTACTAAAAA  268

Query  123  T--ACAAAAATTAGCCTGGCGTGGGCAGGGCGCGGTGGCTCACGCCTGTAATCCCAGCAC  180
            |  | ||| |||| || ||| |||    |||    ||||   |||||||||||||||   
Sbjct  269  TTAAAAAACATTAACCGGGCTTGG---TGGC----TGGCGTGCGCCTGTAATCCCAGTTG  321

Query  181  TTTGGGAGGCCGAGGCGGGCGAATTGCCTGA  211
               ||||||| ||||| || ||||||| |||
Sbjct  322  CGCGGGAGGCTGAGGCAGGGGAATTGCTTGA  352
```

### GML

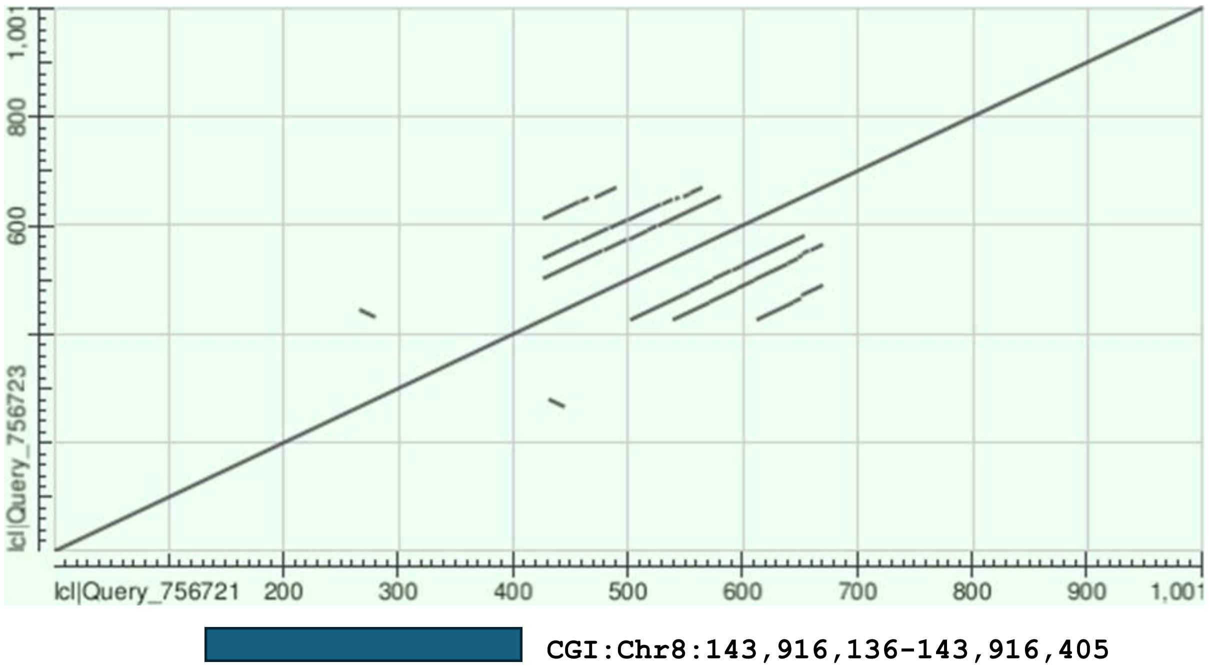


CGI:Chr8:143,916,136-143,916,405

**Alignment within TR (Chr8:143,916,425-143,916,670): identity 78%**

```
Query  2    CCACTCCCCTCGGAGCCCCAGGGAGACCCCCCGAACTCAGCTCCTCTCAGGGGTGCCAGG  61
            |||||||| || | | |||||||||||||||| ||||  |     ||||||||| |||||
Sbjct  188  CCACTCCCATCAGGGTCCCAGGGAGACCCCCC-AACTATG-----CTCAGGGGTCCCAGG  241

Query  62   GGGA  65
            | ||
Sbjct  242  GAGA  245
```

#### LINC01166/LINC01168

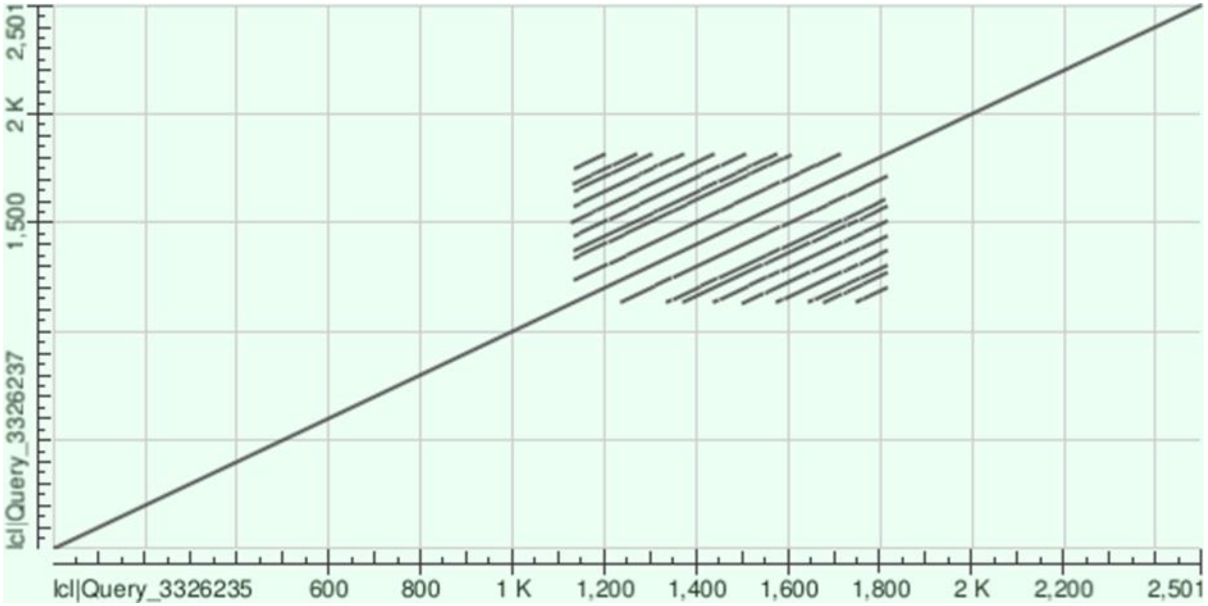


**CGI:Chr10:134,778,684-134,779,073**

**Alignment within TR (Chr10:134,778,630-134,779,310): 79% identity**

```
Query  2    CTGTTCCCACTTCTCACCGGTGTCACCTTCCCACCGTTCCCA---CTCCTCACTGGCGTT  58
            ||||||||||||||||| || || ||||||   | ||||| |   ||||||| || | ||
Sbjct  547  CTGTTCCCACTTCTCACTGGCGTTACCTTCATGCTGTTCCTACTTCTCCTCA-TGCCATT  605

Query  59   ACCTTCCCTCCGTTCCCACTTCTCCTCAACGGCGTTACATTCCTGCGGTTCCCACTTCTC  118
            |||||||  || ||||||||||||||||   || |||| ||||||| ||||||| |||||
Sbjct  606  ACCTTCCTGCCATTCCCACTTCTCCTCA--CGCATTACCTTCCTGCTGTTCCCAATTCTC  663

Query  119  ACCGGTGTCACCTTCC  134
             | || || |||||||
Sbjct  664  TCTGGCGTTACCTTCC  679
```

#### LINC01925

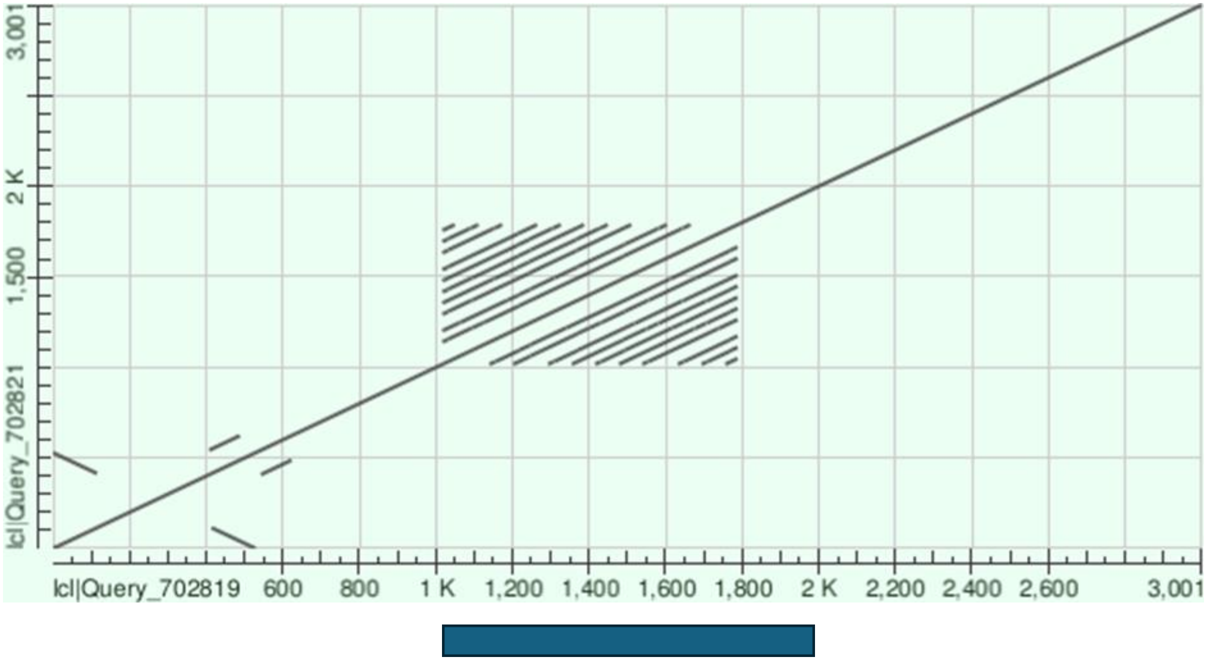


**CGI:Chr18:514,511-515,477**

**Alignment within TR (Chr18:514,520-515,290): 95% identity**

```
Query  1    ACCTCCGGACCC-TCCTCGGACCTCGGCCAGACCTCCAGACCCCTCCTCGGACCTCGGCC  59
            |||||||| ||| |||||||||||||||||||||||| | ||||||||||||||||||||
Sbjct  616  ACCTCCGGGCCCCTCCTCGGACCTCGGCCAGACCTCCGGGCCCCTCCTCGGACCTCGGCC  675

Query  60   AGACCTCCGGACCCCTCCTCGGACCTCGGCCAGACCTCCGGACCCCTCCTCGGACCTCGG  119
            |||||||||| |||||||||||||||||||||||||||||| ||||||||||||||||||
Sbjct  676  AGACCTCCGGGCCCCTCCTCGGACCTCGGCCAGACCTCCGGGCCCCTCCTCGGACCTCGG  735

Query  120  CCAGACCTCCGGACCCCTCCTCGGACCTCGGCCA  153
            ||||||||||||  ||||||||||||||||||||
Sbjct  736  CCAGACCTCCGG-GCCCTCCTCGGACCTCGGCCA  768
```

#### LINC02604

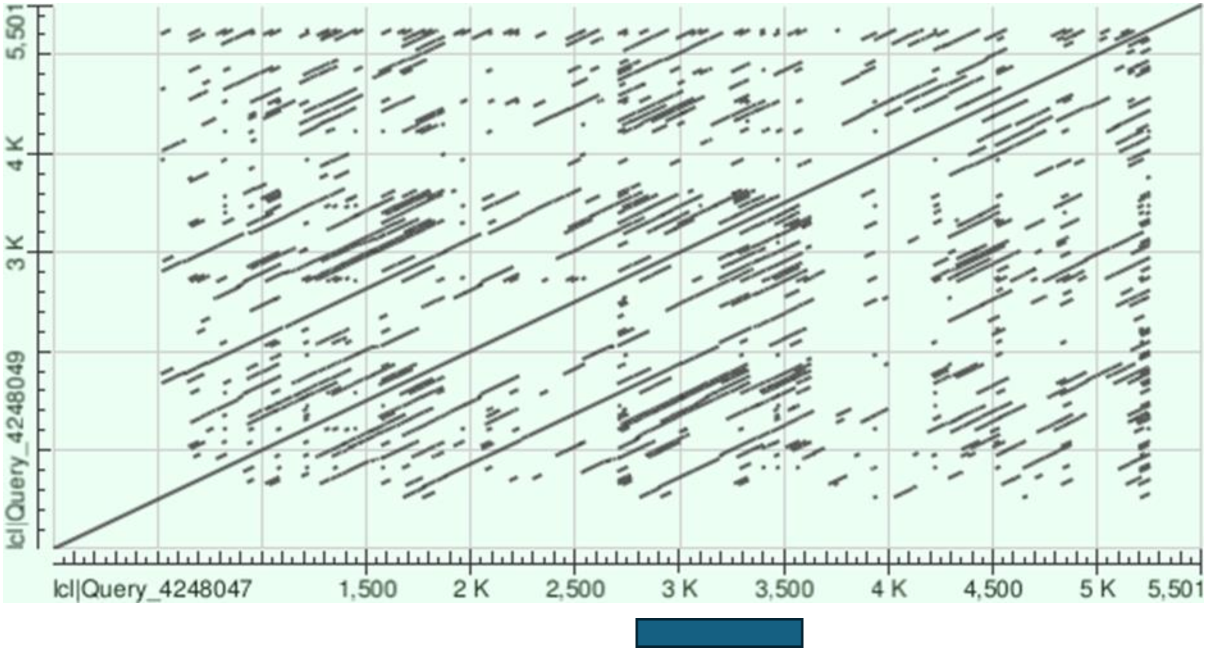


**Alignment within TR (Chr7:66,366,500-66,371,250): 69% identity**

```
Query  271   CCGGCCTCTTCTCCAGTCCCAGTCCTCTCTCGCAGCTGTGTCTACAGGCCCAGCTCCTGC  330
             |||||||||   |||| ||||   ||| ||| |||||||| || | |||||||||||| |
Sbjct  2030  CCGGCCTCT---CCAGGCCCAACTCTCCCTCTCAGCTGTGCCTGCCGGCCCAGCTCCTAC  2086

Query  331   CTCCAAAGAGCCTCTTTTGACTCGGCTCCTACCCAGCTCCTGGCAGCCTTTGTA------  384
             |||  || |||| | || | | | |||||| |||||||||||||||||||||||
Sbjct  2087  CTCGCAAAAGCCACGTTCGGCCCAGCTCCTGCCCAGCTCCTGGCAGCCTTTGTAAACCCC  2146

Query  385   -GG--CCTGAAA-TCT---CTTCCAGTCCA--------GCACTCC------------ATA  417
              ||  |||  || ||    ||| ||| ||         || | ||            |
Sbjct  2147  AGGATCCTCTAAGTCAGGCCTTTCAGGCCCTGCCTTTGGCTCCCCGGTGGCATGGAGAGG  2206

Query  418   CCCAGTCTCC--CCTCACAGCGGCCTTCCCAGGCCCAGCTTTTGCCTCACAGCGGCCTTC  475
             ||||| ||||  ||| ||||||||||  |||||||||||| |||||||||   ||||| |
Sbjct  2207  CCCAG-CTCCTGCCTGACAGCGGCCTCTCCAGGCCCAGCTCTTGCCTCACGTTGGCCTCC  2265

Query  476   CCCGGCCATTTTCTAGCCGGCCTCGCGGTAGCCTCAACAAGCCCAGCTCCCGCCTCACAC  535
             |  |||||  ||   ||| ||||||||| |||| | |||| ||| ||||| ||||| |
Sbjct  2266  CTGGGCCACGTTTCCGCCTGCCTCGCGGCAGCCCCGACAATCCCGGCTCCTGCCTCCCGA  2325

Query  536   TGGCCTCTCTAGGCCCAGCTCAGGCGTCACAGTGGCGT-CTCCAGGCCCAGCTCCCGCC  593
             |||| ||| ||||| || |||  || ||||   ||| | | |||||||   |||| |||
Sbjct  2326  TGGCATCTTTAGGCTCATCTCGTGCCTCACCACGGCCTGCACCAGGCCACACTCCTGCC  2384
```

### NOC2LP2

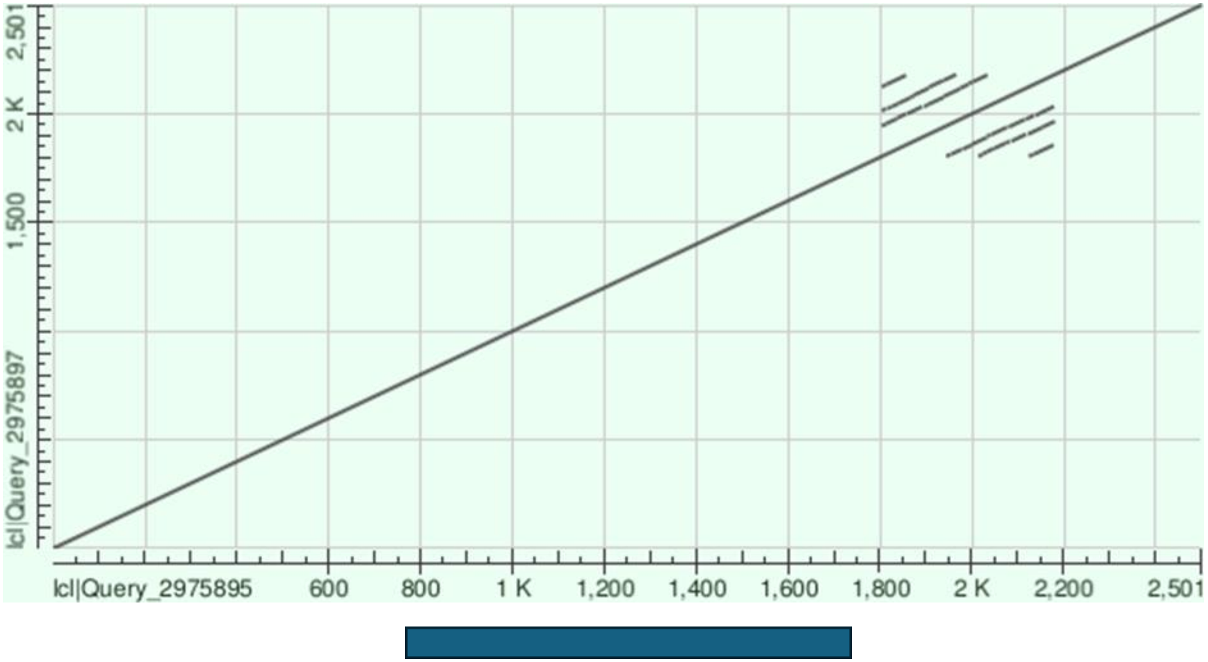


CGI:Chr2:132,202,266-132,203,257

**Alignment within TR (Chr2:132,203,300-132,203,675): 83% identity**

```
Query  5    GGCTTCCGGGGGAGGAGGAGGGCGAGGCCTAGGGGG-CTTCTCGGAGGAGGAGGAGGGCG  63
            ||||||| ||  ||||| ||| |||||||||||||| |||| |||||||||||||||||
Sbjct  215  GGCTTCCCGG--AGGAGAAGG-CGAGGCCTAGGGGGGCTTCCCGGAGGAGGAGGAGGGCT  271

Query  64   AGG-CTTAGGGG---GCTTCCCGGGGGAGGAGGAGGGCGAGGCCTGG----GGGGCTTCT  115
            ||| || |||||   ||||||||| ||||||||||||| ||||||||    ||||||||
Sbjct  272  AGGCCTGAGGGGGGAGCTTCCCGGAGGAGGAGGAGGGCTAGGCCTGGGGGCGGGGCTTCC  331

Query  116  CGGAGGAGGAGGAGGGCGAGGCCTGGGGGGGCTTCCTGGAGGAGGA  161
            | ||||| ||||| |||| ||||  ||||||||||  |||||||||
Sbjct  332  CAGAGGAAGAGGATGGCGGGGCC-AGGGGGGCTTCTCGGAGGAGGA  376
```

#### PROSER2-AS1

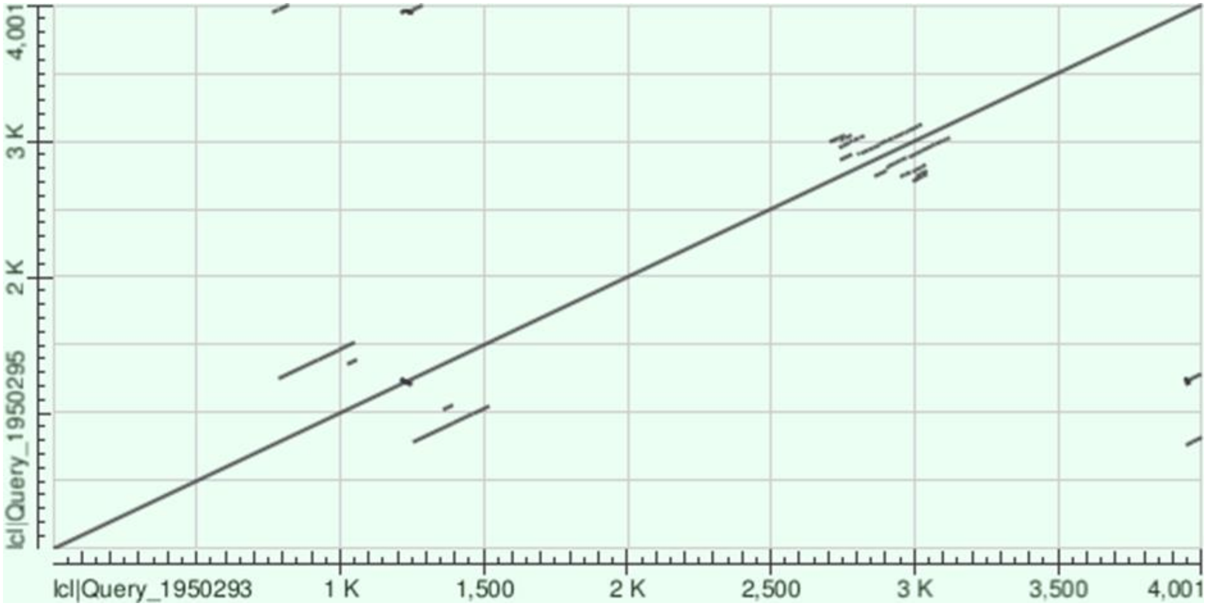


CGI:Chr10:11,936,170-11,937,060

**Alignment within TR (Chr10:11,936,700-11,937,145): 72% identity**

```
Query  38   GGAGGGAGGAGGCCC-GCCGGAAGAAGGGGGAGGGGG------CCCACCGGAAGTGGGGG  90
            |||||||||| |||| || | |||  ||| |||||||      ||||||||||    | |
Sbjct  251  GGAGGGAGGAAGCCCAGCAGAAAGCGGGGCGAGGGGGGGCATACCCACCGGAAA---GCG  307

Query  91   GAGCTTGTCCTCCGGAAAGCGGGGGAGGAGGCCCGC  126
            ||||  | |  || |||   ||||||||||||||||
Sbjct  308  GAGCACGCCAGCCAGAA---GGGGGAGGAGGCCCGC  340
```

### RGPD1(3’)

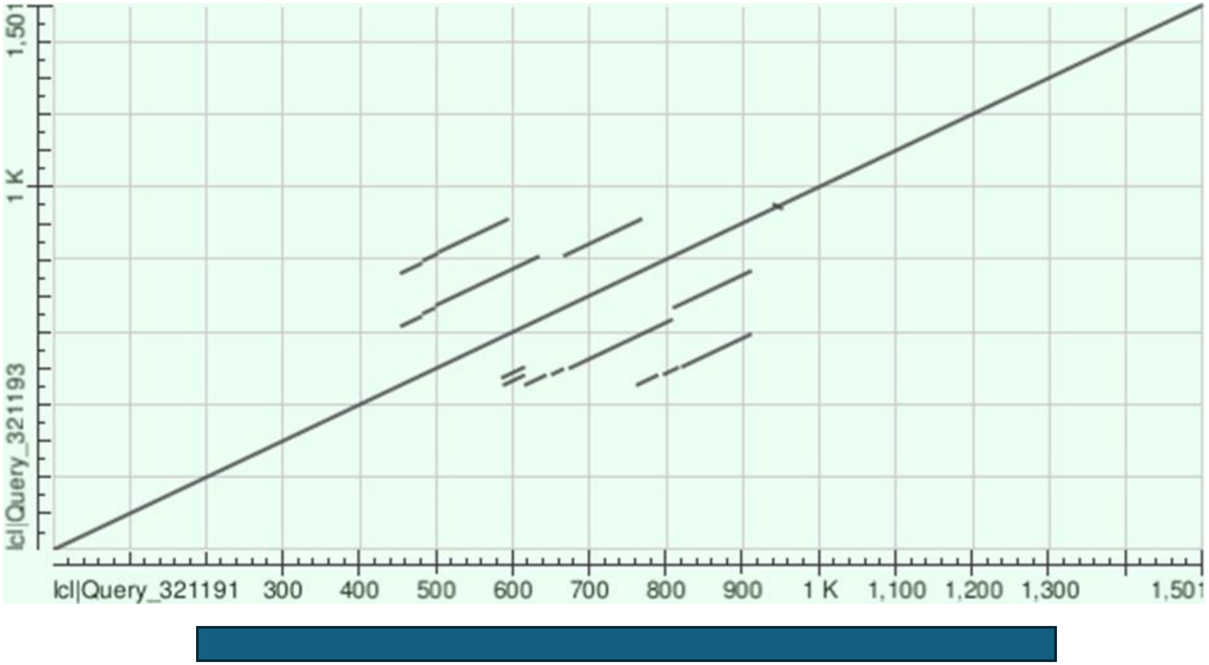


CGI:Chr2:87,169,174-87,170,303

**Alignment within TR (Chr2:87,169,445-87,169,910): 91% identity**

```
Query  9    CGACGGCCTCGACCTGGCCGGGCGGCGGC------CTCGACCTGGCCGGGCGGCGG---C  59
            || ||||||||||||||||||||||||||      |||||||||||||||||||||   |
Sbjct  317  CGCCGGCCTCGACCTGGCCGGGCGGCGGCGGCGGCCTCGACCTGGCCGGGCGGCGGTGGC  376

Query  60   CTCGACCTGGCCGGGCGGCGGCCTCGATGGCTCAGGCGTCATGGCTCCCGACGGGCGCTG  119
            |||||| ||||| |||||||||||||||||||||||||||||||||||| |||||||||||
Sbjct  377  CTCGACGTGGCCCGGCGGCGGCCTCGATGGCTCAGGCGTCATGGCTCCTGACGGGCGCTG  436

Query  120  CTCCCTGGCGCGCTCTGTTGAGGCGCCGGC  149
            ||||||||||||||||||||||||||||||
Sbjct  437  CTCCCTGGCGCGCTCTGTTGAGGCGCCGGC  466
```

**RGPD1(5’)**

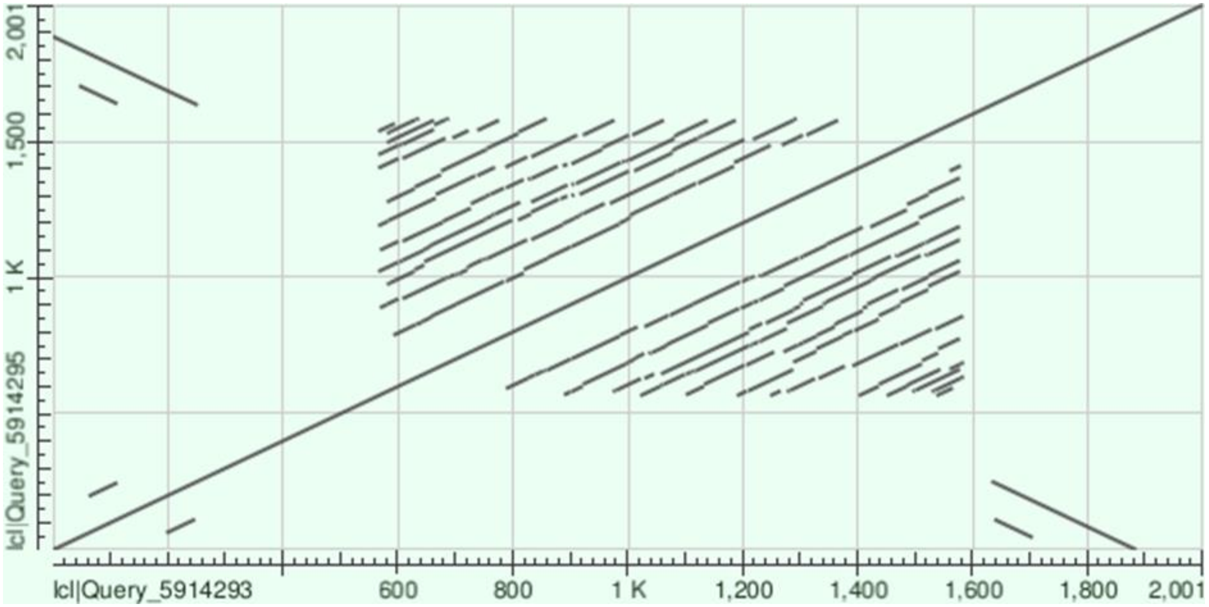


CGI:Chr2:87,140,787-87,141,107

**Alignment within TR (Chr2:87,141,060-87,142,080): 83% identity**

```
Query  582    GGCGG-GGCGGTGGCCTCGACCTGGCCGGGCGGCGGCGGCGGCCTCGGC------CTCGG  634
              ||||| ||||| |||||||||||||||||||||||||||||||  ||||      ||||
Sbjct  1278   GGCGGCGGCGGCGGCCTCGACCTGGCCGGGCGGCGGCGGCGGCGGCGGCGGCCTCGA  1337

Query  635    CCTGGCCGGGCGGCGGCGGCGGCGGCGGCGGCGGCGGCGGC-------------GGCGGC  681
              |||||||||||||||||||||||||||||||||||||||||             ||||||
Sbjct  1338   CCTGGCCGGGCGGCGGCGGCGGCGGCGGCGGCGGCGGCGGCCTCGACCTGGCCGGGCGGC  1397

Query  682    CTCGGCCTCGGCCCCGGCCTGGCCGGGCGGCGGCGGCGGCGGCGGCGGCGGCGGCCTCGG  741
                ||||  ||||| |  ||||| ||||||||||||||||||||||||||||||||  |||
Sbjct  1398   GGCGGCGGCGGCCTCCACCTGGACGGGCGGCGGCGGCGGCGGCGGCGGCGGCGGCGGCGG  1457

Query  742    CCCCGGCCTGGCCG---GGCGGCGGCGGCGGCGGCGGCGGCGGCGGCCTCGGCCTCGGCC  798
              || || ||||||||   ||||||||||||||||||||||||||||||  |||||||| ||
Sbjct  1458   CCTCGACCTGGCCGGGCGGCGGCGGCGGCGGCGGCGGCGGCGGCGGCGGCGGCCTCGACC  1517

Query  799    TGGCCGGACGGC------GGCGGCGGCCTCGGCCTGGCCGGGCGGCGGCGGCGGCCTCGG  852
              ||||||| ||||      ||||||||||||| |||||||||||||||||||||||  |||
Sbjct  1518   TGGCCGGGCGGCGGCGGAGGCGGCGGCCTCGACCTGGCCGGGCGGCGGCGGCGGCGGCGG  1577

Query  853    CCTGGCC  859
              |||||||
Sbjct  1578   CCTGGCC  1584
```

### RGPD3

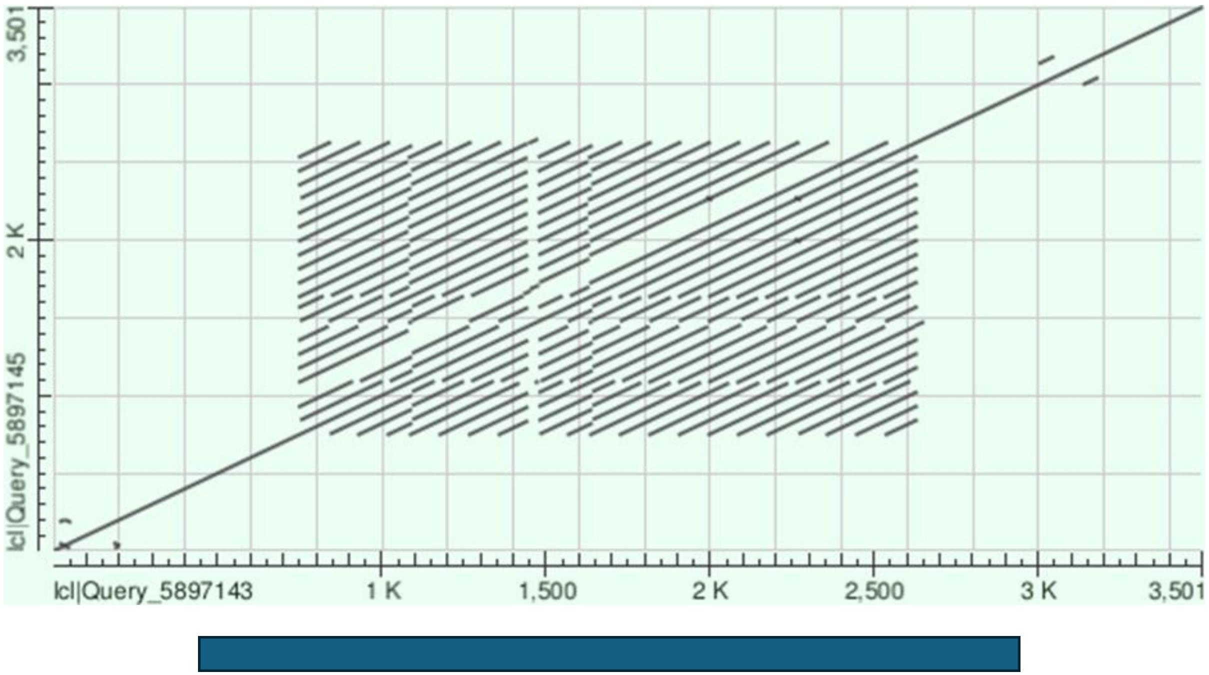


**CGI:Chr2:107,082,329-107,084,936**

**Alignment within TR (Chr2:107,082,750-107,084,645): 94% identity**

```
Query  1     GCGCCGCAACAGAGCGCGCCAGGGAGCAGCGCCCGTCAGGAGCCATGACGCCTGAGCCAT  60
             ||||| ||||||||||||||||||||||||||| ||| ||||||||||||||||||||||
Sbjct  1698  GCGCCTCAACAGAGCGCGCCAGGGAGCAGCGCCGGTCGGGAGCCATGACGCCTGAGCCAT  1757

Query  61    CGAGGCCGCCGCCGGGCCGGGTCCAGGCCACCGCCTCAACAGAGCGCGCCAGGGAGCAGC  120
             |||||||||||| ||||| |||| ||||| ||||||| ||||||||||||||||||||||
Sbjct  1758  CGAGGCCGCCGCAGGGCCAGGTCGAGGCCGCCGCCTCCACAGAGCGCGCCAGGGAGCAGC  1817

Query  121   GCCCGTCGGGAGCCATGACGCCTGAGCCATCGAGGCCGCCGCTGGCCGGGTCGAGGCCG  180
             || ||||||||||||||||||||||||||||||||||||||| |||||||||||||||||
Sbjct  1818  GCTCGTCGGGAGCCATGACGCCTGAGCCATCGAGGCCGCCGCCGGGCCGGGTCGAGGCCG  1877

Query  181   GCGCC  185
              ||||
Sbjct  1878  CCGCC  1882
```

### SHTN1

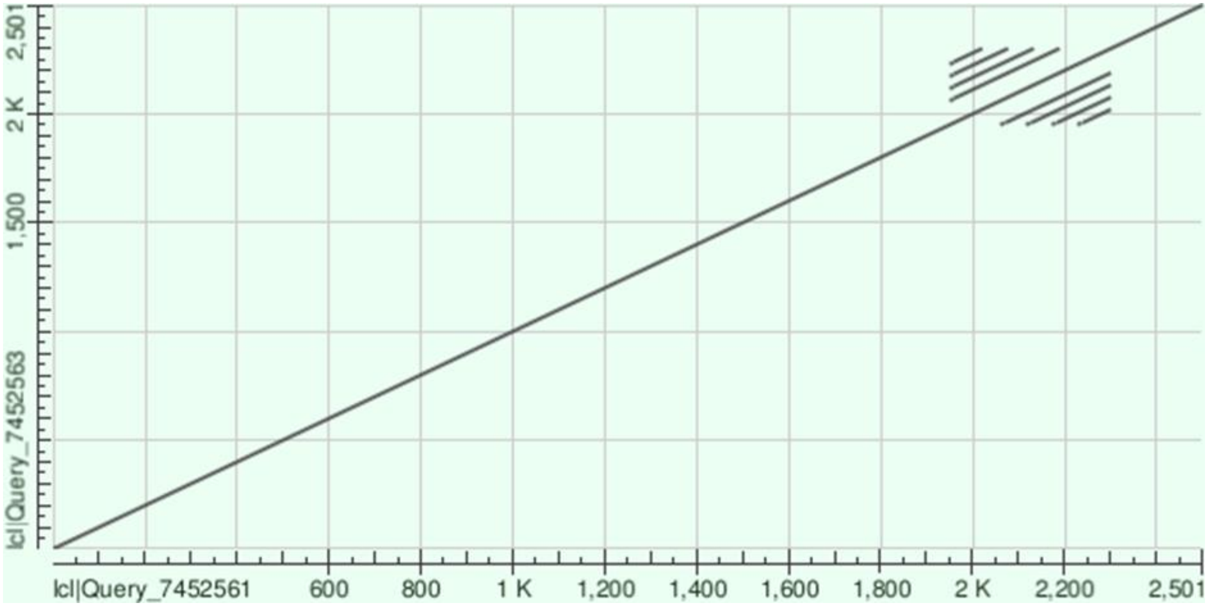


CGI:Chr10:118,885,566-118,885,884

**Alignment within TR (Chr10:118,885,945-118,886,300): 97% Identity**

```
Query  6    CCTAGCACC--TGGCCCTCGCCTCGCCTCCTAGCACGCATGCCCCTCGCCTCGCCTCCTA  63
            ||||||||   || ||||||||||||||||||||||||||||||||||||||||||||||
Sbjct  228  CCTAGCACGCATGCCCCTCGCCTCGCCTCCTAGCACGCATGCCCCTCGCCTCGCCTCCTA  287

Query  64   GCACGCATGCCCCTCGCCTCGCCTCCTAGCACGCATGCCCCTCGCCTCGCCTCCTAGCAC  123
            ||||||||||||||||||||||||||||||||||||||||||||||||||||||||||||
Sbjct  288  GCACGCATGCCCCTCGCCTCGCCTCCTAGCACGCATGCCCCTCGCCTCGCCTCCTAGCAC  347

Query  124  GCATGCCCC  132
            |||||||||
Sbjct  348  GCATGCCCC  356
```

#### SLC45A4/SLC45A4-AS1

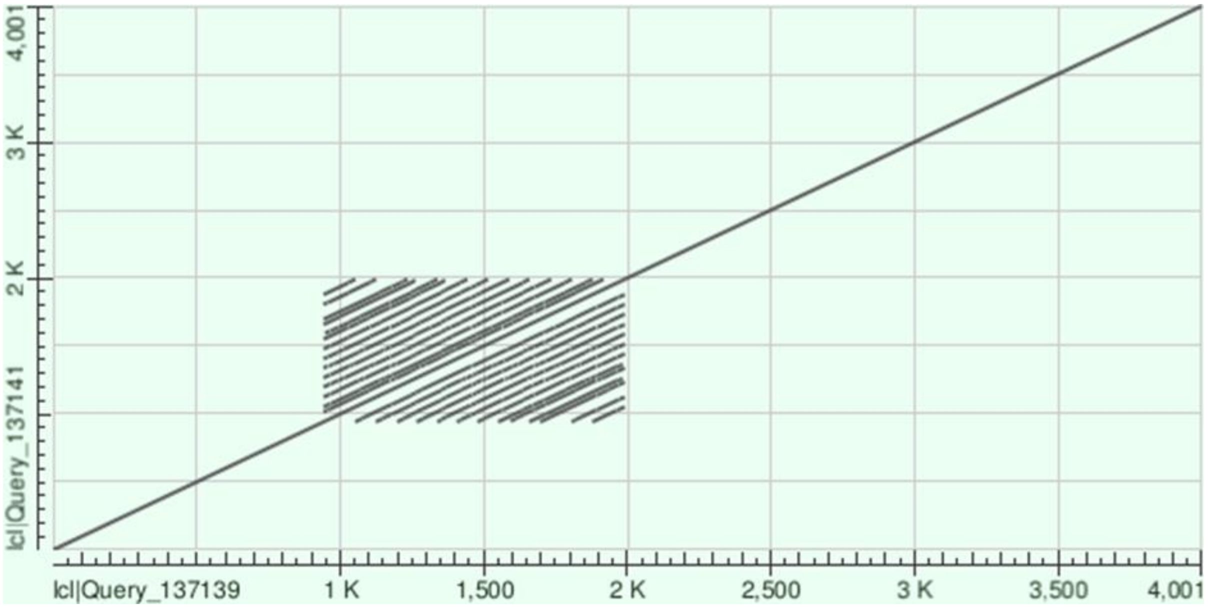


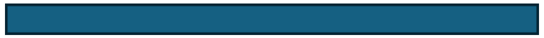

CGI:Chr8:142,262,457-142,264,315

**Alignment within TR (Chr8:142,262,430-142,263,500): 89% identity**

```
Query  13    TCATGCCAACCTGTGCGTCTCTGTTAATTTCCGTGTTTTCACGCCCACCTGCGCGTCTGT  72
             ||| ||| ||||| | || ||||| |||||||||||||||||||||||||||  |||| |
Sbjct  767   TCACGCCCACCTGCGAGTGTCTGTGAATTTCCGTGTTTTCACGCCCACCTGCAAGTCTAT  826

Query  73    GAATTTCCGTGTTTTCACGCCCACCTGCGTGTCTGTGAATTTCCATGTTTTCACGCCCAC  132
             ||||||||||||||||| ||||||||||| |||||||||||||| |||||||| ||||||
Sbjct  827   GAATTTCCGTGTTTTCATGCCCACCTGCGGGTCTGTGAATTTCCGTGTTTTCATGCCCAC  886

Query  133   CTGTGCGTCTGTGAATTTCCGTGTTTTCACGCCCACCTGTGCGTCTGTGAATTTCCGTGT  192
             ||| | ||||||||||||||||||||||| ||||||||||| ||||||||||||||||||
Sbjct  887   CTGCGTGTCTGTGAATTTCCGTGTTTTCATGCCCACCTGTGTGTCTGTGAATTTCCGTGT  946

Query  193   TTTCACGCCCACCTG--TGCGTCTGTGAATTTCCGTGTTTTCACGCCCACCTGCGAGTGT  250
             ||||| |||||||||  || ||||||||||||| ||||||||| ||||||||| | ||  
Sbjct  947   TTTCATGCCCACCTGCGTGTGTCTGTGAATTTCTGTGTTTTCATGCCCACCTGTGCGT--  1004

Query  251   CTGTGAATTTCCGTGTTTTCATGCCCACCTGTGCAT--CTGTGAATTTCCGTGTTTT  305
             |||||| ||||||||||||||||||||||||||| |  |||||||||||| || |||
Sbjct  1005  CTGTGATTTTCCGTGTTTTCATGCCCACCTGTGCGTGCCTGTGAATTTCCATGCTTT  1061
```

### Suppl. Fig. 3

(A)

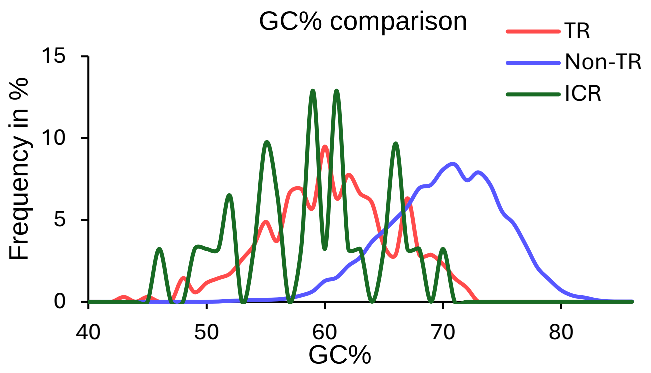


(B)

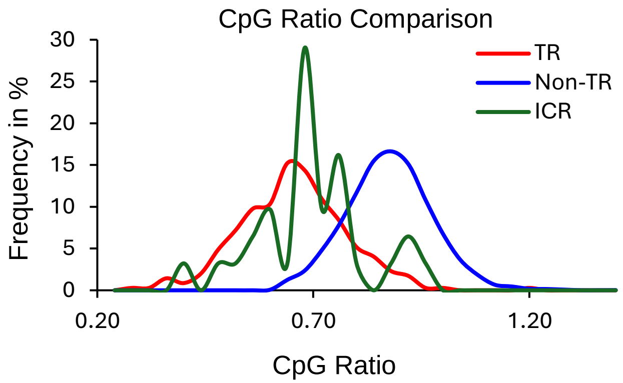

Suppl. Fig. 4

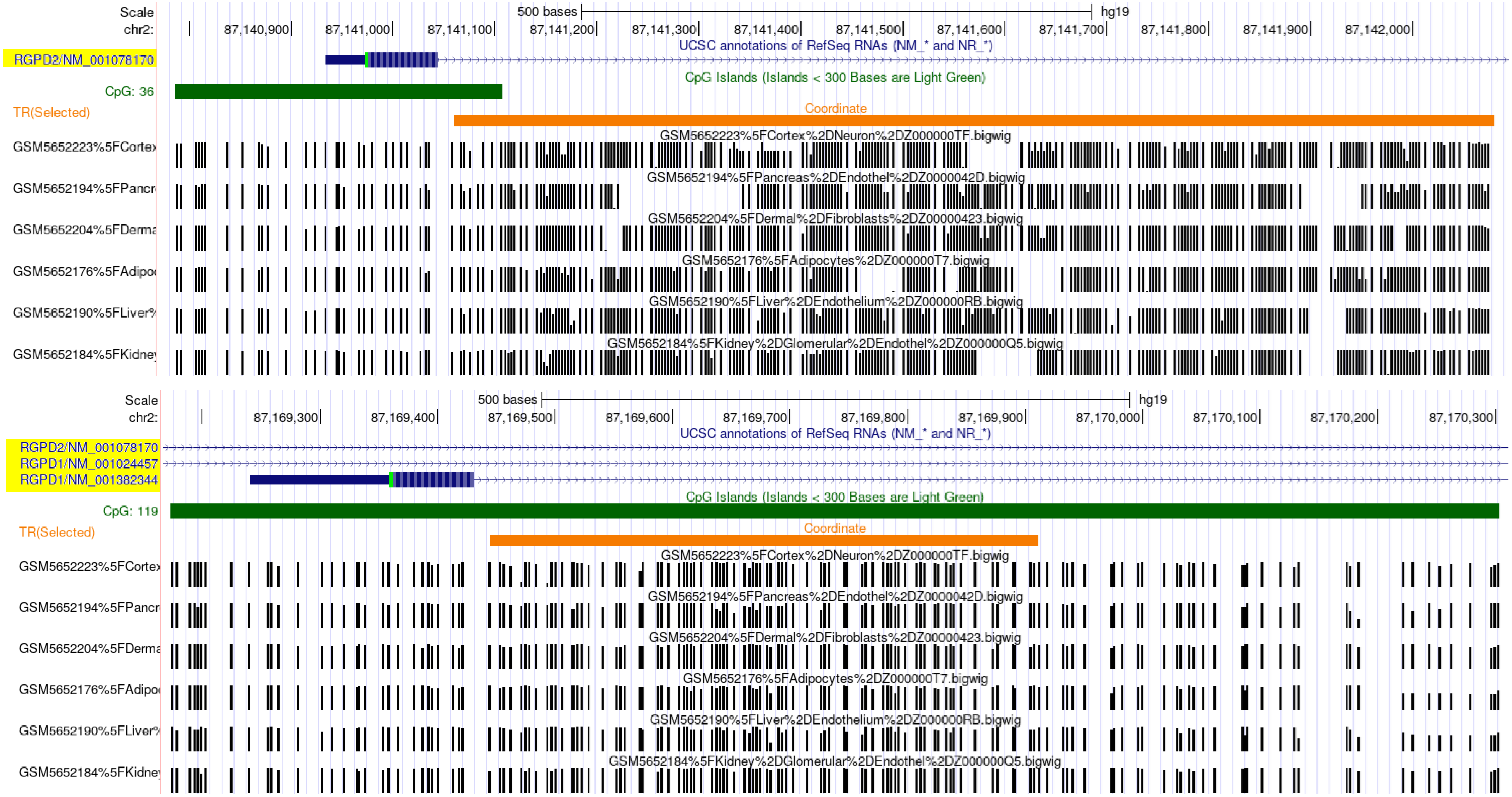
Scale
chr2:
500 bases
hg19
87,140,900
87,141,000
87,141,100
87,141,200
87,141,300
87,141,400
87,141,500
87,141,600
87,141,700
87,141,800
87,141,900
87,142,000
UCSC annotations of RefSeq RNAs (NM_* and NR_*)
RGPD2/NM_001078170
CpG Islands (Islands < 300 Bases are Light Green)
CpG: 36
TR(Selected)
Coordinate
GSM5652223%5FCortex%2DNeuron%2DZ000000TF.bigwig
GSM5652194%5FPancreas%2DEndothel%2DZ0000042D.bigwig
GSM5652204%5FDermal%2DFibroblasts%2DZ00000423.bigwig
GSM5652176%5FAdipocytes%2DZ000000T7.bigwig
GSM5652190%5FLiver%2DEndothelium%2DZ000000RB.bigwig
GSM5652184%5FKidney%2DGlomerular%2DEndothel%2DZ000000Q5.bigwig
Scale
chr2:
500 bases
hg19
87,169,300
87,169,400
87,169,500
87,169,600
87,169,700
87,169,800
87,169,900
87,170,000
87,170,100
87,170,200
87,170,300
UCSC annotations of RefSeq RNAs (NM_* and NR_*)
RGPD2/NM_001078170
RGPD1/NM_001024457
RGPD1/NM_001382344
CpG Islands (Islands < 300 Bases are Light Green)
CpG: 119
TR(Selected)
Coordinate
GSM5652223%5FCortex%2DNeuron%2DZ000000TF.bigwig
GSM5652194%5FPancreas%2DEndothel%2DZ0000042D.bigwig
GSM5652204%5FDermal%2DFibroblasts%2DZ00000423.bigwig
GSM5652176%5FAdipocytes%2DZ000000T7.bigwig
GSM5652190%5FLiver%2DEndothelium%2DZ000000RB.bigwig
GSM5652184%5FKidney%2DGlomerular%2DEndothel%2DZ000000Q5.bigwig

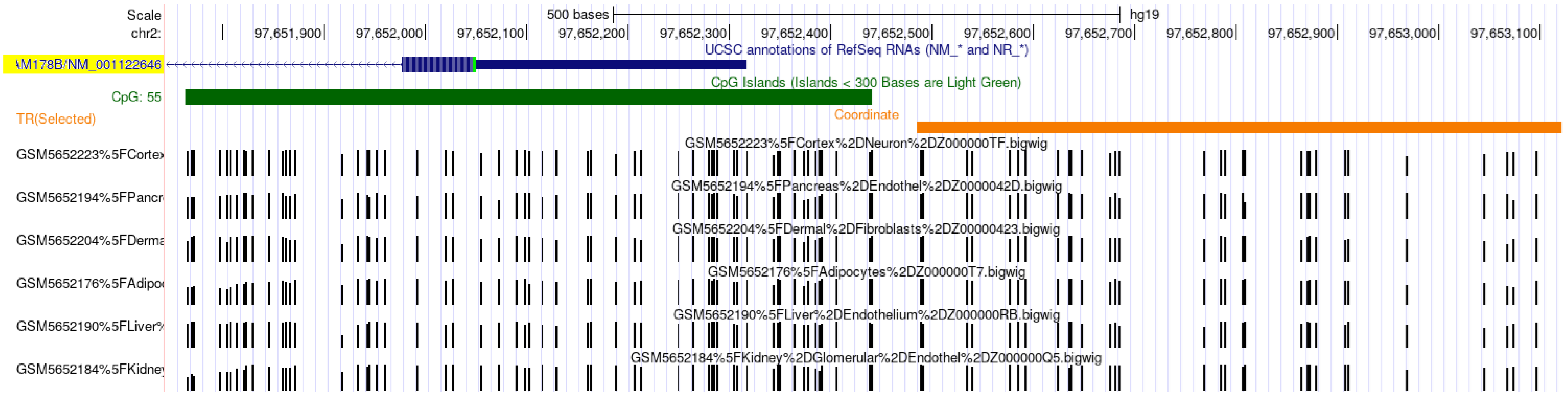
Scale
chr2:
500 bases
hg19
97,651,900
97,652,000
97,652,100
97,652,200
97,652,300
97,652,400
97,652,500
97,652,600
97,652,700
97,652,800
97,652,900
97,653,000
97,653,100
UCSC annotations of RefSeq RNAs (NM_* and NR_*)
\M178B/NM_001122646
CpG Islands (Islands < 300 Bases are Light Green)
CpG: 55
TR(Selected)
Coordinate
GSM5652223%5FCortex%2DNeuron%2DZ000000TF.bigwig
GSM5652223%5FCortex
GSM5652194%5FPancreas%2DEndothel%2DZ0000042D.bigwig
GSM5652194%5FPancr
GSM5652204%5FDermal%2DFibroblasts%2DZ00000423.bigwig
GSM5652204%5FDerma
GSM5652176%5FAdipocytes%2DZ000000T7.bigwig
GSM5652176%5FAdipoc
GSM5652190%5FLiver%2DEndothelium%2DZ000000RB.bigwig
GSM5652190%5FLiver%
GSM5652184%5FKidney%2DGlomerular%2DEndothel%2DZ000000Q5.bigwig
GSM5652184%5FKidney

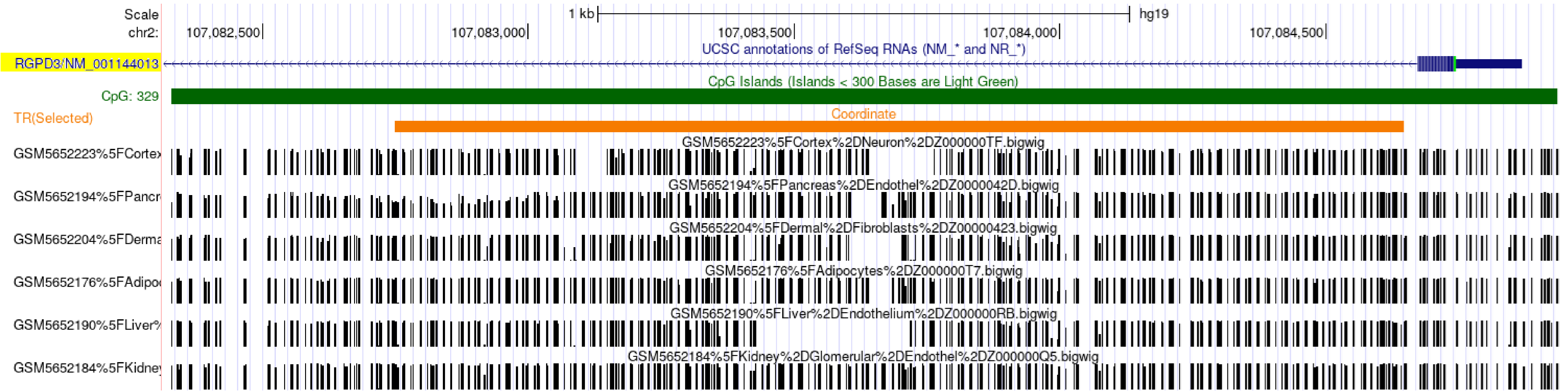
Scale
chr2:
1 kb
hg19
107,082,500
107,083,000
107,083,500
107,084,000
107,084,500
UCSC annotations of RefSeq RNAs (NM_* and NR_*)
RGPD3/NM_001144013
CpG Islands (Islands < 300 Bases are Light Green)
CpG: 329
TR(Selected)
Coordinate
GSM5652223%5FCortex%2DNeuron%2DZ000000TF.bigwig
GSM5652223%5FCortex
GSM5652194%5FPancreas%2DEndothel%2DZ0000042D.bigwig
GSM5652194%5FPancr
GSM5652204%5FDermal%2DFibroblasts%2DZ00000423.bigwig
GSM5652204%5FDerma
GSM5652176%5FAdipocytes%2DZ000000T7.bigwig
GSM5652176%5FAdipoc
GSM5652190%5FLiver%2DEndothelium%2DZ000000RB.bigwig
GSM5652190%5FLiver%
GSM5652184%5FKidney%2DGlomerular%2DEndothel%2DZ000000Q5.bigwig
GSM5652184%5FKidney

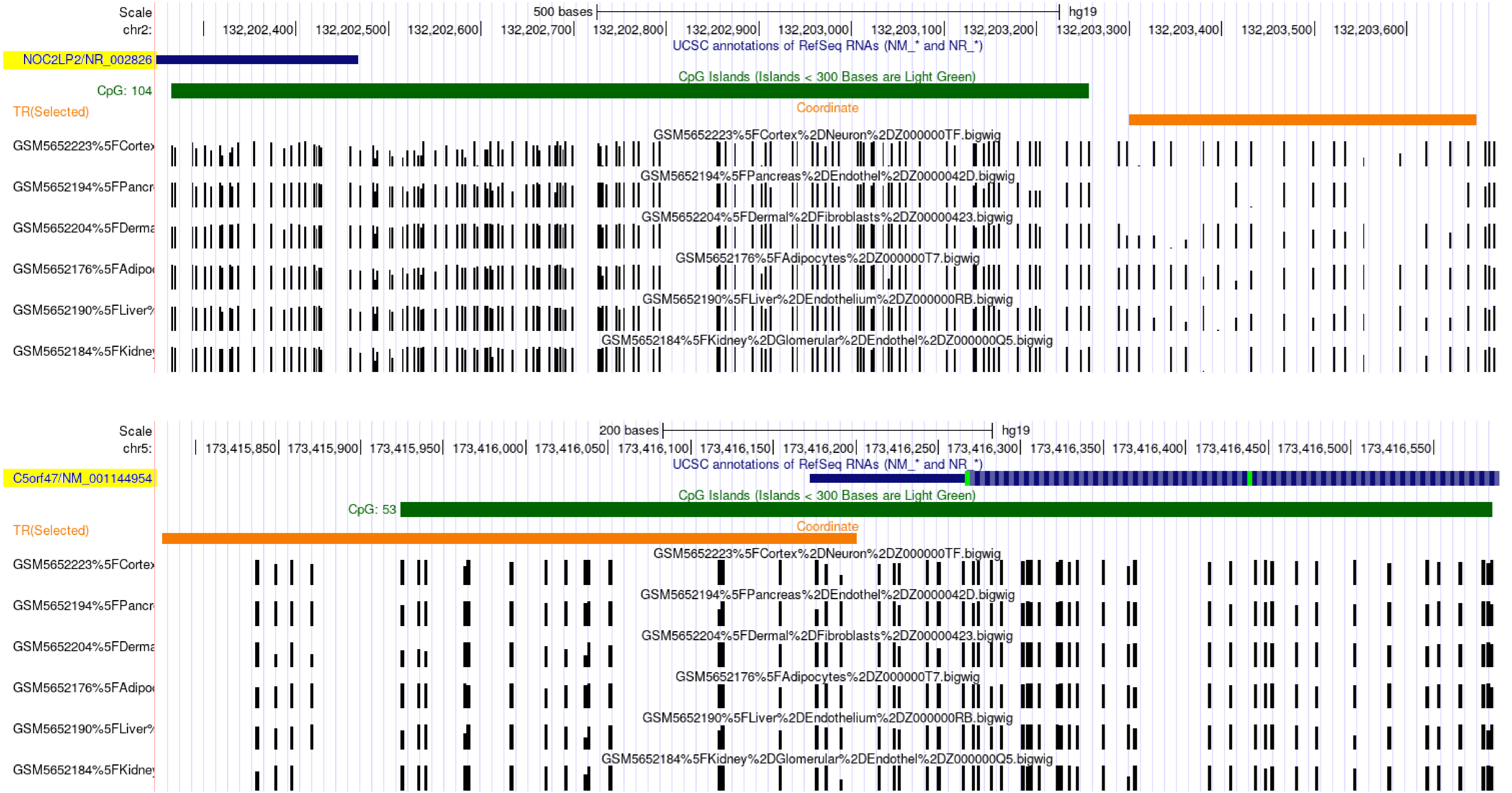
Scale
chr2:
500 bases
hg19
132,202,400
132,202,500
132,202,600
132,202,700
132,202,800
132,202,900
132,203,000
132,203,100
132,203,200
132,203,300
132,203,400
132,203,500
132,203,600
UCSC annotations of RefSeq RNAs (NM_* and NR_*)
NOC2LP2/NR_002826
CpG Islands (Islands < 300 Bases are Light Green)
CpG: 104
TR(Selected)
Coordinate
GSM5652223%5FCortex%2DNeuron%2DZ000000TF.bigwig
GSM5652194%5FPancreas%2DEndothel%2DZ0000042D.bigwig
GSM5652204%5FDermal%2DFibroblasts%2DZ00000423.bigwig
GSM5652176%5FAdipocytes%2DZ000000T7.bigwig
GSM5652190%5FLiver%2DEndothelium%2DZ000000RB.bigwig
GSM5652184%5FKidney%2DGlomerular%2DEndothel%2DZ000000Q5.bigwig
Scale
chr5:
200 bases
hg19
173,415,850
173,415,900
173,415,950
173,416,000
173,416,050
173,416,100
173,416,150
173,416,200
173,416,250
173,416,300
173,416,350
173,416,400
173,416,450
173,416,500
173,416,550
UCSC annotations of RefSeq RNAs (NM_* and NR_*)
C5orf47/NM_001144954
CpG Islands (Islands < 300 Bases are Light Green)
CpG: 53
TR(Selected)
Coordinate
GSM5652223%5FCortex%2DNeuron%2DZ000000TF.bigwig
GSM5652194%5FPancreas%2DEndothel%2DZ0000042D.bigwig
GSM5652204%5FDermal%2DFibroblasts%2DZ00000423.bigwig
GSM5652176%5FAdipocytes%2DZ000000T7.bigwig
GSM5652190%5FLiver%2DEndothelium%2DZ000000RB.bigwig
GSM5652184%5FKidney%2DGlomerular%2DEndothel%2DZ000000Q5.bigwig

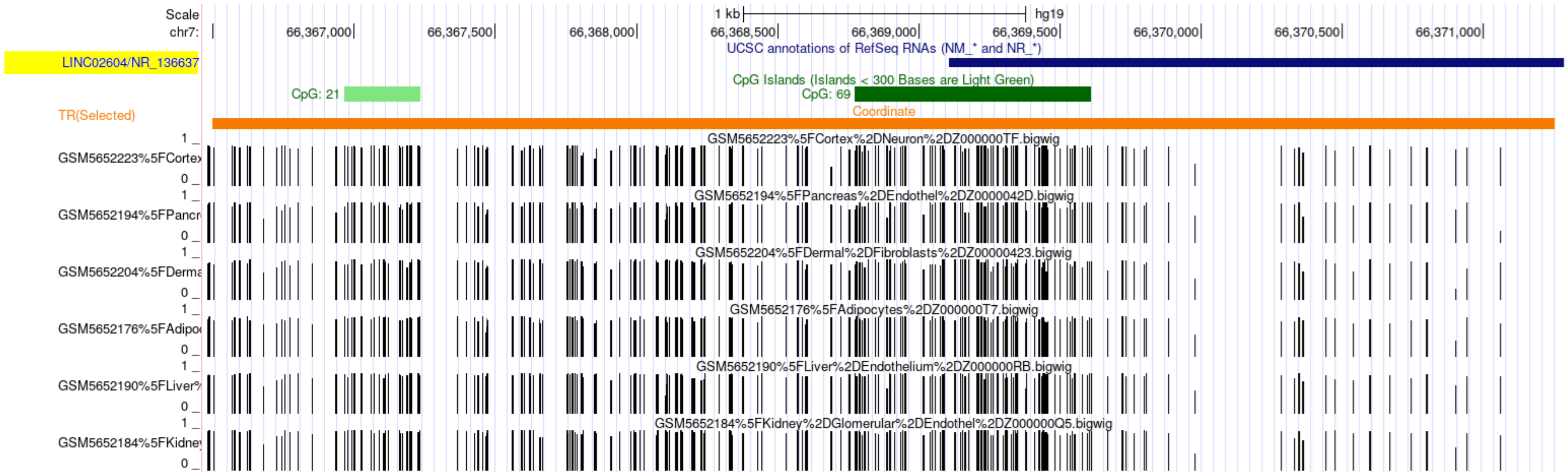

Scale
chr7:
1 kb
hg19
66,367,000
66,367,500
66,368,000
66,368,500
66,369,000
66,369,500
66,370,000
66,370,500
66,371,000
UCSC annotations of RefSeq RNAs (NM_* and NR_*)
LINC02604/NR_136637
CpG Islands (Islands < 300 Bases are Light Green)
CpG: 21
CpG: 69
Coordinate
TR(Selected)
GSM5652223%5FCortex%2DNeuron%2DZ000000TF.bigwig
GSM5652223%5FCortex
GSM5652194%5FPancreas%2DEndothel%2DZ0000042D.bigwig
GSM5652194%5FPancr
GSM5652204%5FDermal%2DFibroblasts%2DZ00000423.bigwig
GSM5652204%5FDerma
GSM5652176%5FAdipocytes%2DZ000000T7.bigwig
GSM5652176%5FAdipo
GSM5652190%5FLiver%2DEndothelium%2DZ000000RB.bigwig
GSM5652190%5FLiver
GSM5652184%5FKidney%2DGlomerular%2DEndothel%2DZ000000Q5.bigwig
GSM5652184%5FKidne


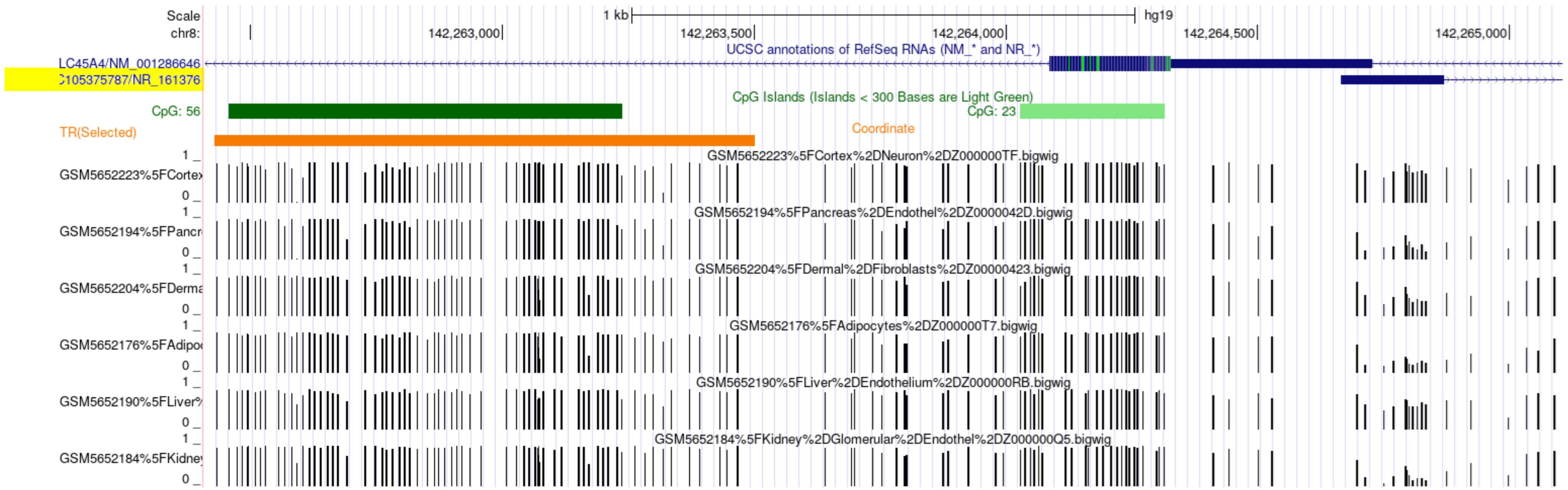

Scale
chr8:
1 kb
hg19
142,263,000
142,263,500
142,264,000
142,264,500
142,265,000
UCSC annotations of RefSeq RNAs (NM_* and NR_*)
LC45A4/NM_001286646
105375787/NR_161376
CpG Islands (Islands < 300 Bases are Light Green)
CpG: 56
CpG: 23
Coordinate
TR(Selected)
GSM5652223%5FCortex%2DNeuron%2DZ000000TF.bigwig
GSM5652223%5FCortex
GSM5652194%5FPancreas%2DEndothel%2DZ0000042D.bigwig
GSM5652194%5FPancr
GSM5652204%5FDermal%2DFibroblasts%2DZ00000423.bigwig
GSM5652204%5FDerma
GSM5652176%5FAdipocytes%2DZ000000T7.bigwig
GSM5652176%5FAdipo
GSM5652190%5FLiver%2DEndothelium%2DZ000000RB.bigwig
GSM5652190%5FLiver
GSM5652184%5FKidney%2DGlomerular%2DEndothel%2DZ000000Q5.bigwig
GSM5652184%5FKidne

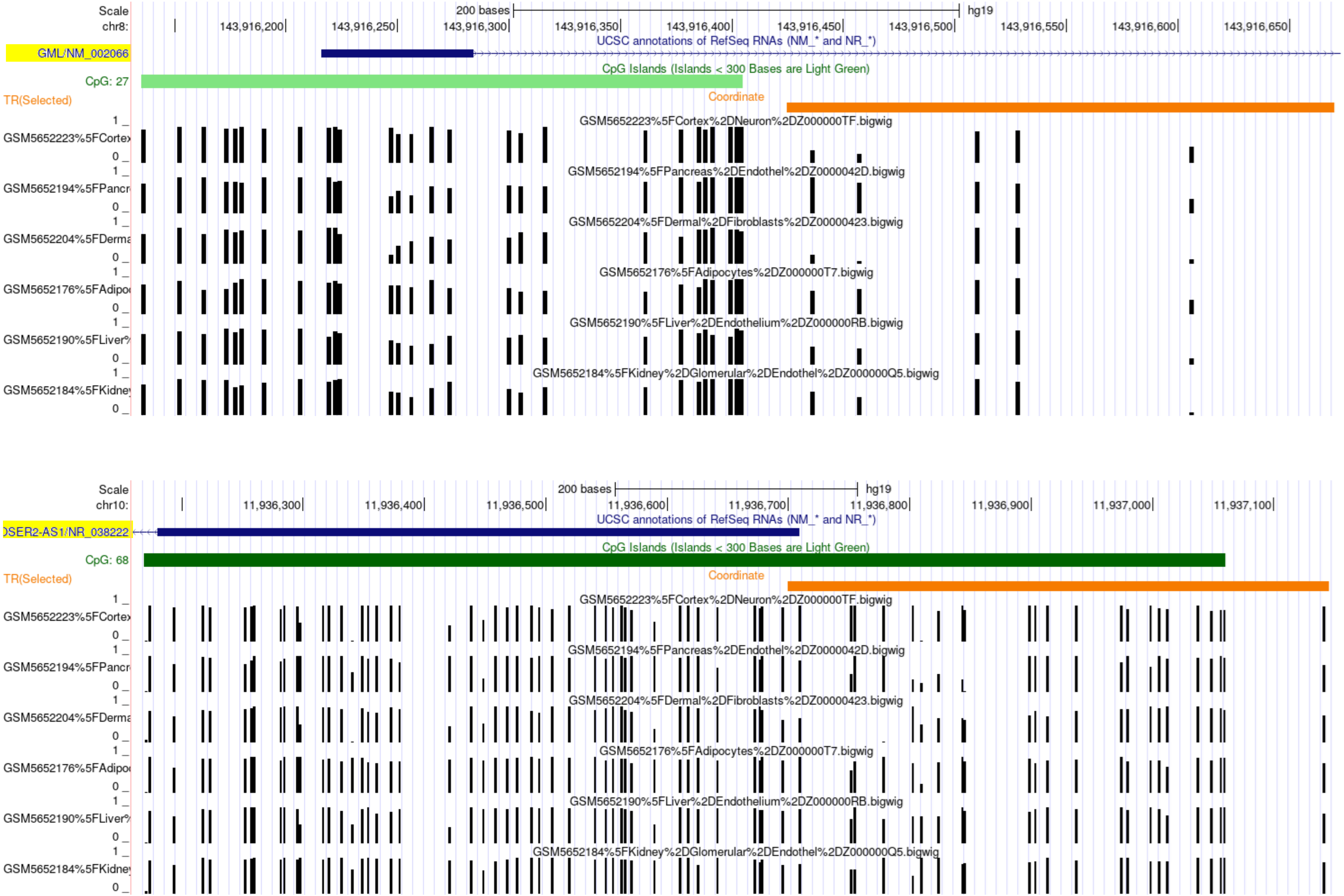
Scale
chr8:
200 bases
hg19
143,916,200
143,916,250
143,916,300
143,916,350
143,916,400
143,916,450
143,916,500
143,916,550
143,916,600
143,916,650
UCSC annotations of RefSeq RNAs (NM_* and NR_*)
GML/NM_002066
CpG Islands (Islands < 300 Bases are Light Green)
CpG: 27
TR(Selected)
Coordinate
GSM5652223%5FCortex%2DNeuron%2DZ000000TF.bigwig
GSM5652194%5FPancreas%2DEndothel%2DZ0000042D.bigwig
GSM5652204%5FDermal%2DFibroblasts%2DZ00000423.bigwig
GSM5652176%5FAdipocytes%2DZ000000T7.bigwig
GSM5652190%5FLiver%2DEndothelium%2DZ000000RB.bigwig
GSM5652184%5FKidney%2DGlomerular%2DEndothel%2DZ000000Q5.bigwig
Scale
chr10:
200 bases
hg19
11,936,300
11,936,400
11,936,500
11,936,600
11,936,700
11,936,800
11,936,900
11,937,000
11,937,100
UCSC annotations of RefSeq RNAs (NM_* and NR_*)
)SER2-AS1/NR_038222
CpG Islands (Islands < 300 Bases are Light Green)
CpG: 68
TR(Selected)
Coordinate
GSM5652223%5FCortex%2DNeuron%2DZ000000TF.bigwig
GSM5652194%5FPancreas%2DEndothel%2DZ0000042D.bigwig
GSM5652204%5FDermal%2DFibroblasts%2DZ00000423.bigwig
GSM5652176%5FAdipocytes%2DZ000000T7.bigwig
GSM5652190%5FLiver%2DEndothelium%2DZ000000RB.bigwig
GSM5652184%5FKidney%2DGlomerular%2DEndothel%2DZ000000Q5.bigwig

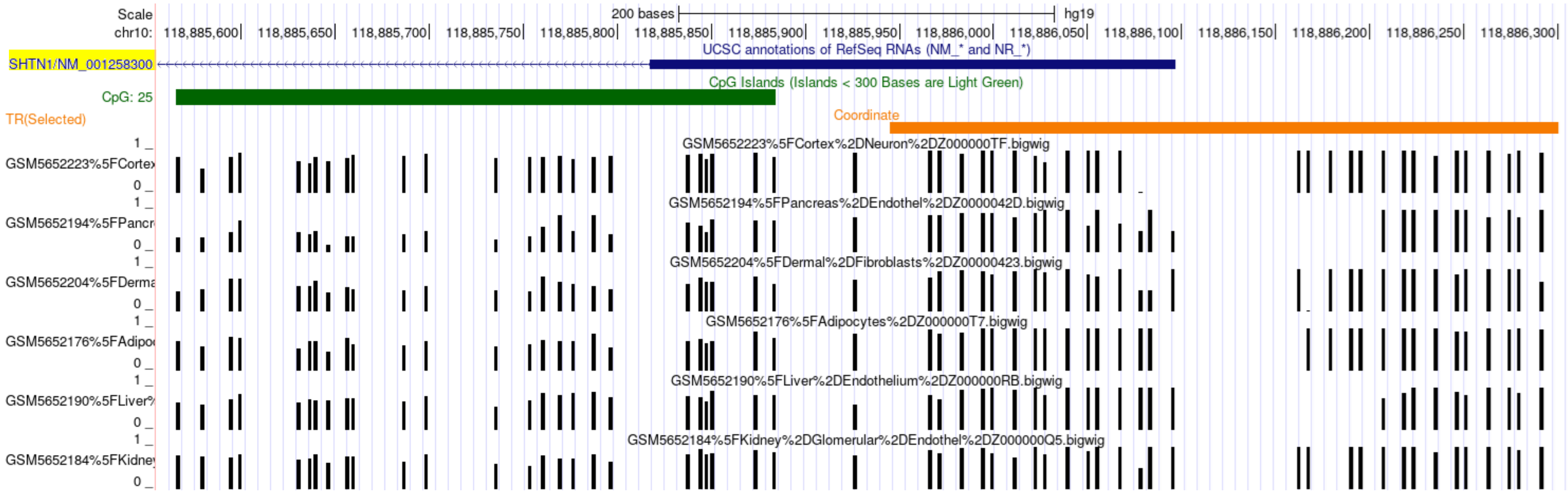
Scale
chr10:
200 bases
hg19
118,885,600
118,885,650
118,885,700
118,885,750
118,885,800
118,885,850
118,885,900
118,885,950
118,886,000
118,886,050
118,886,100
118,886,150
118,886,200
118,886,250
118,886,300
UCSC annotations of RefSeq RNAs (NM_* and NR_*)
SHTN1/NM_001258300
CpG Islands (Islands < 300 Bases are Light Green)
CpG: 25
TR(Selected)
Coordinate
GSM5652223%5FCortex%2DNeuron%2DZ000000TF.bigwig
GSM5652194%5FPancreas%2DEndothel%2DZ0000042D.bigwig
GSM5652204%5FDermal%2DFibroblasts%2DZ00000423.bigwig
GSM5652176%5FAdipocytes%2DZ000000T7.bigwig
GSM5652190%5FLiver%2DEndothelium%2DZ000000RB.bigwig
GSM5652184%5FKidney%2DGlomerular%2DEndothel%2DZ000000Q5.bigwig

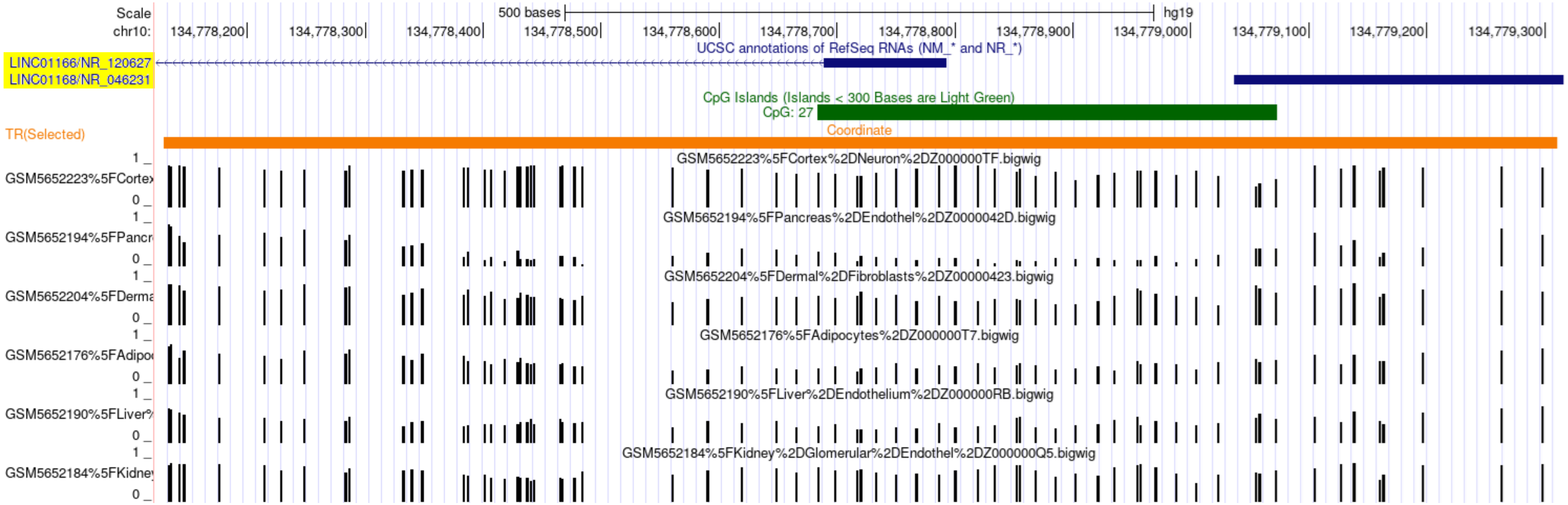
Scale
chr10:
500 bases
hg19
134,778,200
134,778,300
134,778,400
134,778,500
134,778,600
134,778,700
134,778,800
134,778,900
134,779,000
134,779,100
134,779,200
134,779,300
UCSC annotations of RefSeq RNAs (NM_* and NR_*)
LINC01166/NR_120627
LINC01168/NR_046231
CpG Islands (Islands < 300 Bases are Light Green)
CpG: 27
TR(Selected)
Coordinate
GSM5652223%5FCortex%2DNeuron%2DZ000000TF.bigwig
GSM5652194%5FPancreas%2DEndothel%2DZ0000042D.bigwig
GSM5652204%5FDermal%2DFibroblasts%2DZ00000423.bigwig
GSM5652176%5FAdipocytes%2DZ000000T7.bigwig
GSM5652190%5FLiver%2DEndothelium%2DZ000000RB.bigwig
GSM5652184%5FKidney%2DGlomerular%2DEndothel%2DZ000000Q5.bigwig

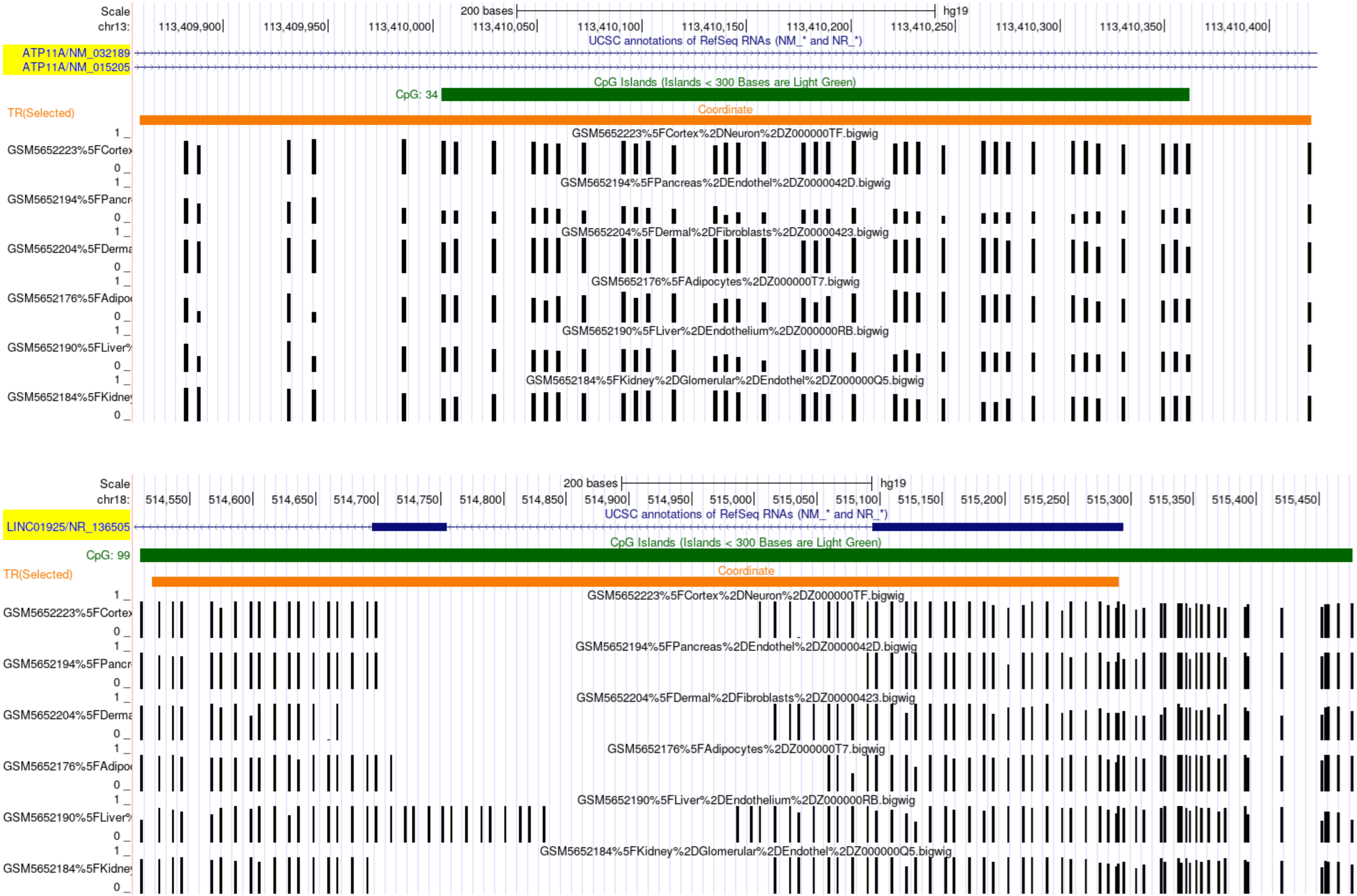

Scale
chr13:
200 bases
hg19
113,409,900
113,409,950
113,410,000
113,410,050
113,410,100
113,410,150
113,410,200
113,410,250
113,410,300
113,410,350
113,410,400
UCSC annotations of RefSeq RNAs (NM_* and NR_*)
ATP11A/NM_032189
ATP11A/NM_015205
CpG Islands (Islands < 300 Bases are Light Green)
CpG: 34
TR(Selected)
Coordinate
GSM5652223%5FCortex%2DNeuron%2DZ000000TF.bigwig
GSM5652194%5FPancreas%2DEndothel%2DZ0000042D.bigwig
GSM5652204%5FDermal%2DFibroblasts%2DZ00000423.bigwig
GSM5652176%5FAdipocytes%2DZ000000T7.bigwig
GSM5652190%5FLiver%2DEndothelium%2DZ000000RB.bigwig
GSM5652184%5FKidney%2DGlomerular%2DEndothel%2DZ000000Q5.bigwig
Scale
chr18:
200 bases
hg19
514,550
514,600
514,650
514,700
514,750
514,800
514,850
514,900
514,950
515,000
515,050
515,100
515,150
515,200
515,250
515,300
515,350
515,400
515,450
UCSC annotations of RefSeq RNAs (NM_* and NR_*)
LINC01925/NR_136505
CpG Islands (Islands < 300 Bases are Light Green)
CpG: 99
TR(Selected)
Coordinate
GSM5652223%5FCortex%2DNeuron%2DZ000000TF.bigwig
GSM5652194%5FPancreas%2DEndothel%2DZ0000042D.bigwig
GSM5652204%5FDermal%2DFibroblasts%2DZ00000423.bigwig
GSM5652176%5FAdipocytes%2DZ000000T7.bigwig
GSM5652190%5FLiver%2DEndothelium%2DZ000000RB.bigwig
GSM5652184%5FKidney%2DGlomerular%2DEndothel%2DZ000000Q5.bigwig

Suppl. Fig. 5

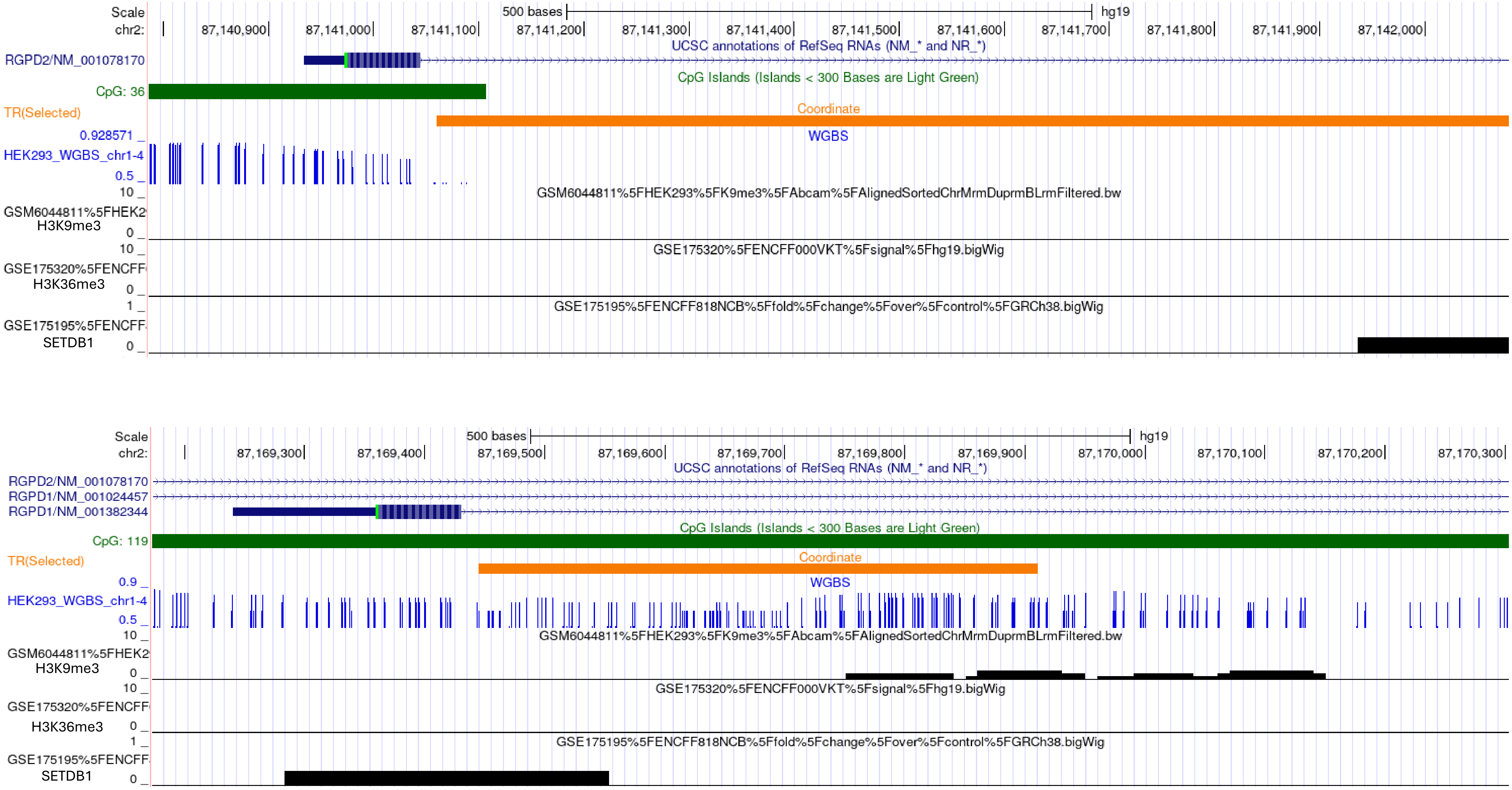
Scale
chr2:
500 bases
hg19
87,140,900
87,141,000
87,141,100
87,141,200
87,141,300
87,141,400
87,141,500
87,141,600
87,141,700
87,141,800
87,141,900
87,142,000
UCSC annotations of RefSeq RNAs (NM_* and NR_*)
RGPD2/NM_001078170
CpG Islands (Islands < 300 Bases are Light Green)
CpG: 36
TR(Selected)
Coordinate
0.928571
HEK293_WGBS_chr1-4
0.5
WGBS
10
GSM6044811%5FHEK293%5FK9me3%5FAbcam%5FAlignedSortedChrMrmDuprmBLrmFiltered.bw
H3K9me3
0
GSE175320%5FENCFF000VKT%5Fsignal%5Fhg19.bigWig
H3K36me3
1
GSE175195%5FENCFF818NCB%5Ffold%5Fchange%5Fover%5Fcontrol%5FGRCh38.bigWig
SETDB1
Scale
chr2:
500 bases
hg19
87,169,300
87,169,400
87,169,500
87,169,600
87,169,700
87,169,800
87,169,900
87,170,000
87,170,100
87,170,200
87,170,300
UCSC annotations of RefSeq RNAs (NM_* and NR_*)
RGPD2/NM_001078170
RGPD1/NM_001024457
RGPD1/NM_001382344
CpG Islands (Islands < 300 Bases are Light Green)
CpG: 119
TR(Selected)
Coordinate
0.9
HEK293_WGBS_chr1-4
0.5
WGBS
GSM6044811%5FHEK293%5FK9me3%5FAbcam%5FAlignedSortedChrMrmDuprmBLrmFiltered.bw
H3K9me3
GSE175320%5FENCFF000VKT%5Fsignal%5Fhg19.bigWig
H3K36me3
GSE175195%5FENCFF818NCB%5Ffold%5Fchange%5Fover%5Fcontrol%5FGRCh38.bigWig
SETDB1

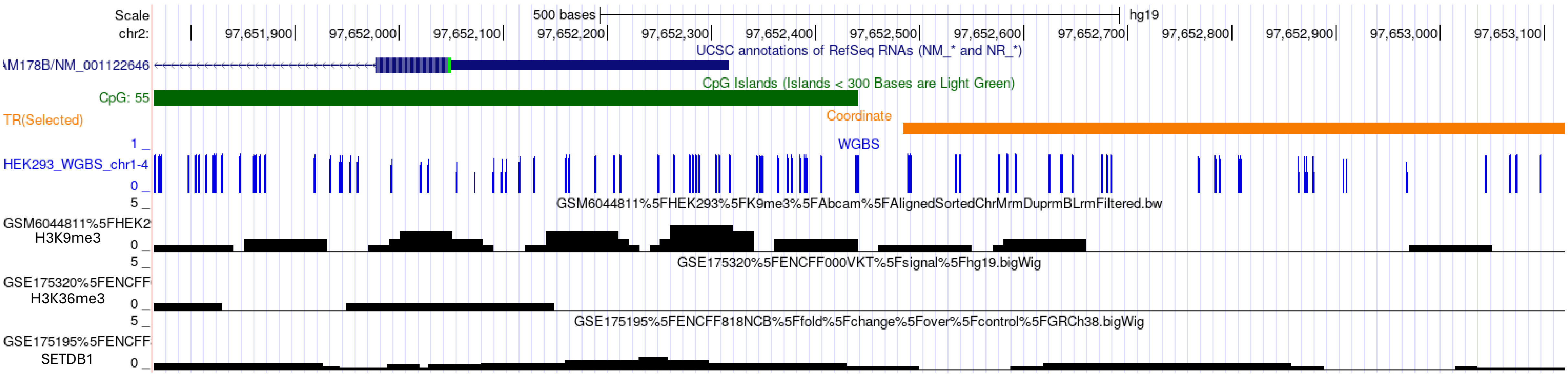
Scale
chr2:
500 bases
hg19
97,651,900
97,652,000
97,652,100
97,652,200
97,652,300
97,652,400
97,652,500
97,652,600
97,652,700
97,652,800
97,652,900
97,653,000
97,653,100
UCSC annotations of RefSeq RNAs (NM_* and NR_*)
AM178B/NM_001122646
CpG Islands (Islands < 300 Bases are Light Green)
CpG: 55
TR(Selected)
Coordinate
WGBS
1
HEK293_WGBS_chr1-4
0
5
GSM6044811%5FHEK293%5FK9me3%5FAbcam%5FAlignedSortedChrMrmDuprmBLrmFiltered.bw
GSM6044811%5FHEK2
H3K9me3
0
5
GSE175320%5FENCFF000VKT%5Fsignal%5Fhg19.bigWig
GSE175320%5FENCFF
H3K36me3
0
5
GSE175195%5FENCFF818NCB%5Ffold%5Fchange%5Fover%5Fcontrol%5FGRCh38.bigWig
GSE175195%5FENCFF
SETDB1
0

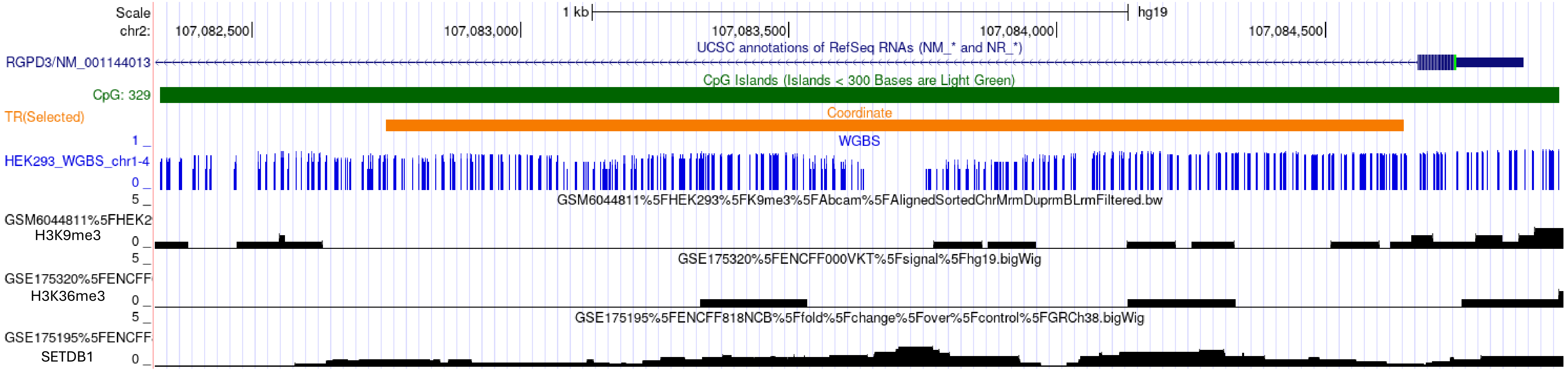
Scale
chr2:
1 kb
hg19
107,082,500
107,083,000
107,083,500
107,084,000
107,084,500
UCSC annotations of RefSeq RNAs (NM_* and NR_*)
RGPD3/NM_001144013
CpG Islands (Islands < 300 Bases are Light Green)
CpG: 329
TR(Selected)
Coordinate
WGBS
1
HEK293_WGBS_chr1-4
0
5
GSM6044811%5FHEK293%5FK9me3%5FAbcam%5FAlignedSortedChrMrmDuprmBLrmFiltered.bw
GSM6044811%5FHEK2
H3K9me3
0
5
GSE175320%5FENCFF000VKT%5Fsignal%5Fhg19.bigWig
GSE175320%5FENCFF
H3K36me3
0
5
GSE175195%5FENCFF818NCB%5Ffold%5Fchange%5Fover%5Fcontrol%5FGRCh38.bigWig
GSE175195%5FENCFF
SETDB1
0

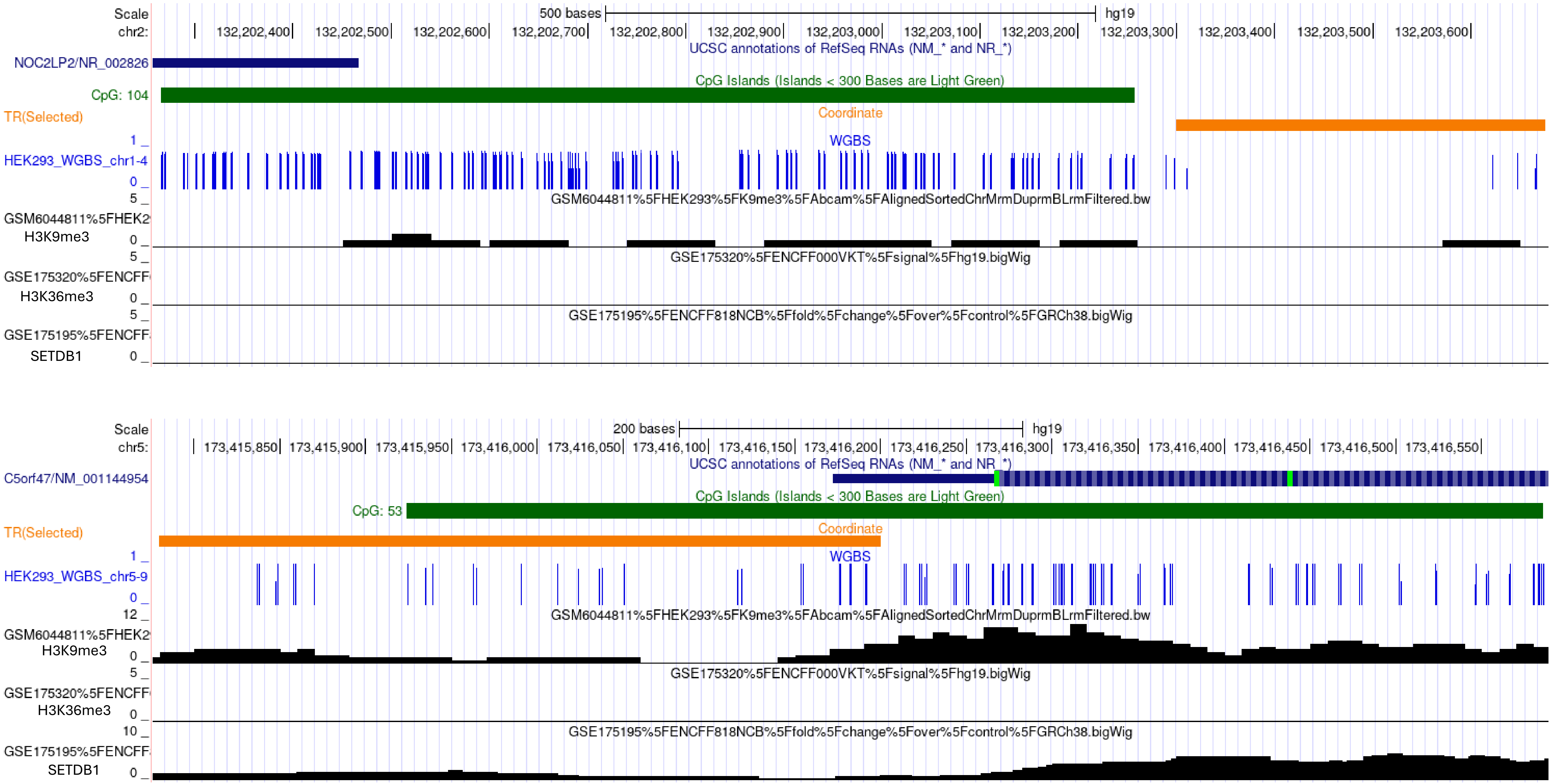

Scale
chr2:
500 bases
hg19
132,202,400
132,202,500
132,202,600
132,202,700
132,202,800
132,202,900
132,203,000
132,203,100
132,203,200
132,203,300
132,203,400
132,203,500
132,203,600
UCSC annotations of RefSeq RNAs (NM_* and NR_*)
NOC2LP2/NR_002826
CpG Islands (Islands < 300 Bases are Light Green)
CpG: 104
TR(Selected)
Coordinate
WGBS
HEK293_WGBS_chr1-4
GSM6044811%5FHEK293%5FK9me3%5FAbcam%5FAlignedSortedChrMrmDuprmBLrmFiltered.bw
GSM6044811%5FHEK2
H3K9me3
GSE175320%5FENCFF000VKT%5Fsignal%5Fhg19.bigWig
GSE175320%5FENCFF
H3K36me3
GSE175195%5FENCFF818NCB%5Ffold%5Fchange%5Fover%5Fcontrol%5FGRCh38.bigWig
GSE175195%5FENCFF
SETDB1
Scale
chr5:
200 bases
hg19
173,415,850
173,415,900
173,415,950
173,416,000
173,416,050
173,416,100
173,416,150
173,416,200
173,416,250
173,416,300
173,416,350
173,416,400
173,416,450
173,416,500
173,416,550
UCSC annotations of RefSeq RNAs (NM_* and NR_*)
C5orf47/NM_001144954
CpG Islands (Islands < 300 Bases are Light Green)
CpG: 53
TR(Selected)
Coordinate
WGBS
HEK293_WGBS_chr5-9
GSM6044811%5FHEK293%5FK9me3%5FAbcam%5FAlignedSortedChrMrmDuprmBLrmFiltered.bw
GSM6044811%5FHEK2
H3K9me3
GSE175320%5FENCFF000VKT%5Fsignal%5Fhg19.bigWig
GSE175320%5FENCFF
H3K36me3
GSE175195%5FENCFF818NCB%5Ffold%5Fchange%5Fover%5Fcontrol%5FGRCh38.bigWig
GSE175195%5FENCFF
SETDB1

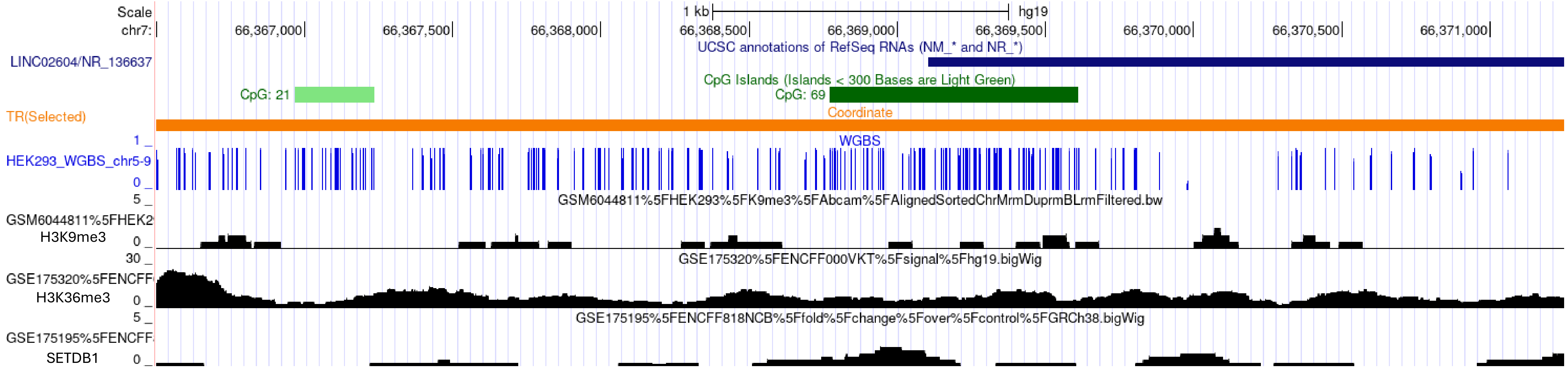
Scale
chr7:
66,367,000
66,367,500
66,368,000
66,368,500
66,369,000
66,369,500
66,370,000
66,370,500
66,371,000
1 kb
hg19
UCSC annotations of RefSeq RNAs (NM_* and NR_*)
LINC02604/NR_136637
CpG Islands (Islands < 300 Bases are Light Green)
CpG: 21
CpG: 69
TR(Selected)
Coordinate
WGBS
HEK293_WGBS_chr5-9
GSM6044811%5FHEK293%5FK9me3%5FAbcam%5FAlignedSortedChrMrmDuprmBLrmFiltered.bw
H3K9me3
GSE175320%5FENCFF000VKT%5Fsignal%5Fhg19.bigWig
H3K36me3
GSE175195%5FENCFF818NCB%5Ffold%5Fchange%5Fover%5Fcontrol%5FGRCh38.bigWig
SETDB1

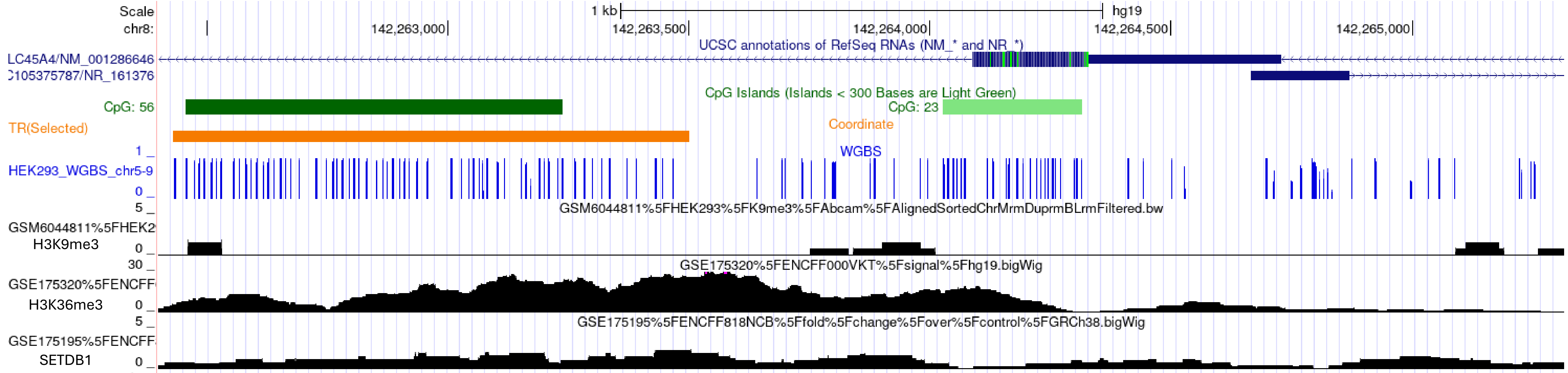
Scale
chr8:
142,263,000
142,263,500
142,264,000
142,264,500
142,265,000
1 kb
hg19
UCSC annotations of RefSeq RNAs (NM_* and NR_*)
LC45A4/NM_001286646
105375787/NR_161376
CpG Islands (Islands < 300 Bases are Light Green)
CpG: 56
CpG: 23
TR(Selected)
Coordinate
WGBS
HEK293_WGBS_chr5-9
GSM6044811%5FHEK293%5FK9me3%5FAbcam%5FAlignedSortedChrMrmDuprmBLrmFiltered.bw
H3K9me3
GSE175320%5FENCFF000VKT%5Fsignal%5Fhg19.bigWig
H3K36me3
GSE175195%5FENCFF818NCB%5Ffold%5Fchange%5Fover%5Fcontrol%5FGRCh38.bigWig
SETDB1

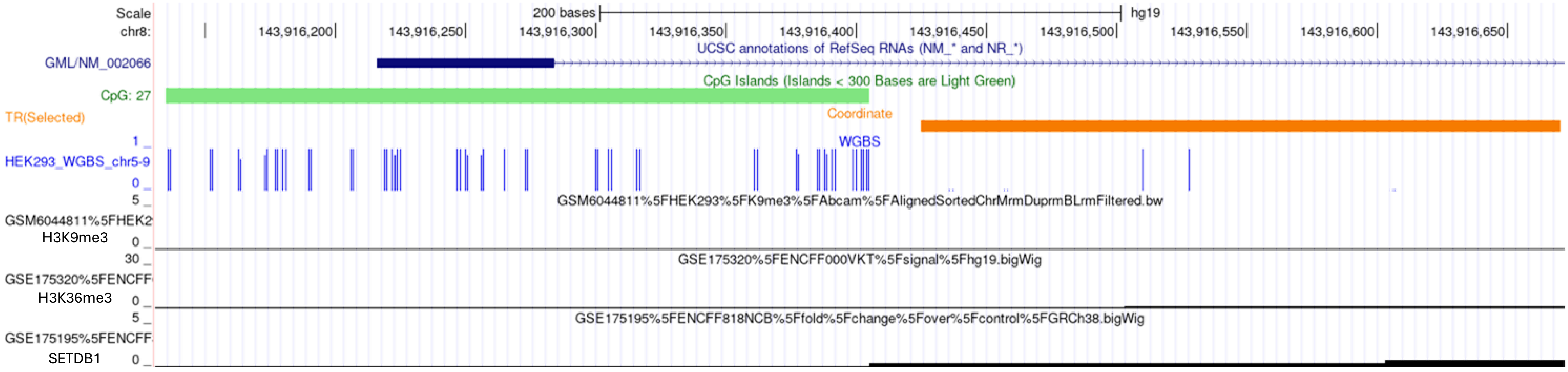
Scale
chr8:
200 bases
hg19
143,916,200
143,916,250
143,916,300
143,916,350
143,916,400
143,916,450
143,916,500
143,916,550
143,916,600
143,916,650
UCSC annotations of RefSeq RNAs (NM_* and NR_*)
GML/NM_002066
CpG Islands (Islands < 300 Bases are Light Green)
CpG: 27
TR(Selected)
Coordinate
WGBS
HEK293_WGBS_chr5-9
GSM6044811%5FHEK293%5FK9me3%5FAbcam%5FAlignedSortedChrMrmDuprmBLrmFiltered.bw
GSM6044811%5FHEK2
H3K9me3
GSE175320%5FENCFF000VKT%5Fsignal%5Fhg19.bigWig
GSE175320%5FENCFF
H3K36me3
GSE175195%5FENCFF818NCB%5Ffold%5Fchange%5Fover%5Fcontrol%5FGRCh38.bigWig
GSE175195%5FENCFF
SETDB1

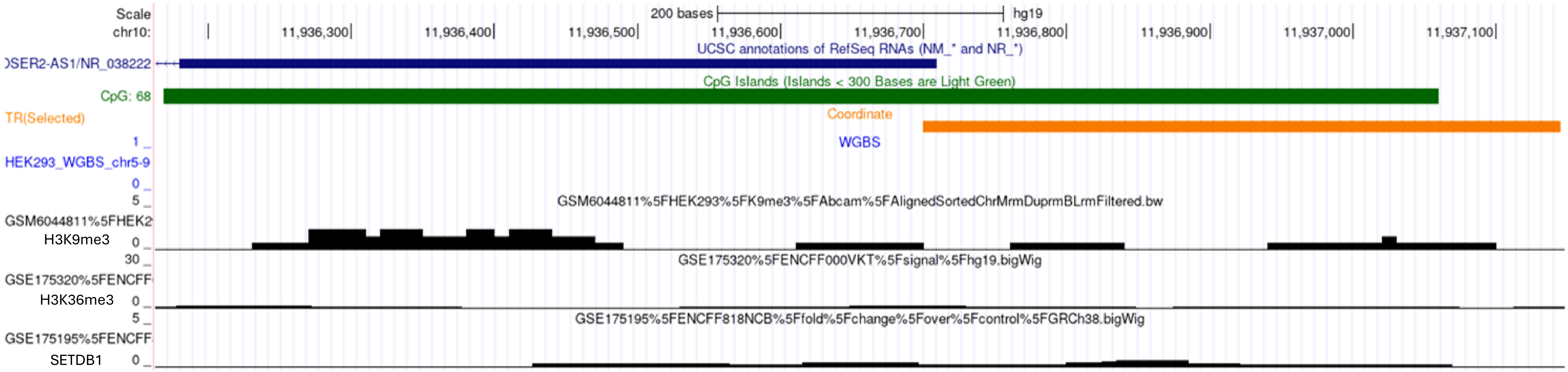
Scale
chr10:
200 bases
hg19
11,936,300
11,936,400
11,936,500
11,936,600
11,936,700
11,936,800
11,936,900
11,937,000
11,937,100
UCSC annotations of RefSeq RNAs (NM_* and NR_*)
)SER2-AS1/NR_038222
CpG Islands (Islands < 300 Bases are Light Green)
CpG: 68
TR(Selected)
Coordinate
WGBS
HEK293_WGBS_chr5-9
GSM6044811%5FHEK293%5FK9me3%5FAbcam%5FAlignedSortedChrMrmDuprmBLrmFiltered.bw
GSM6044811%5FHEK2
H3K9me3
GSE175320%5FENCFF000VKT%5Fsignal%5Fhg19.bigWig
GSE175320%5FENCFF
H3K36me3
GSE175195%5FENCFF818NCB%5Ffold%5Fchange%5Fover%5Fcontrol%5FGRCh38.bigWig
GSE175195%5FENCFF
SETDB1

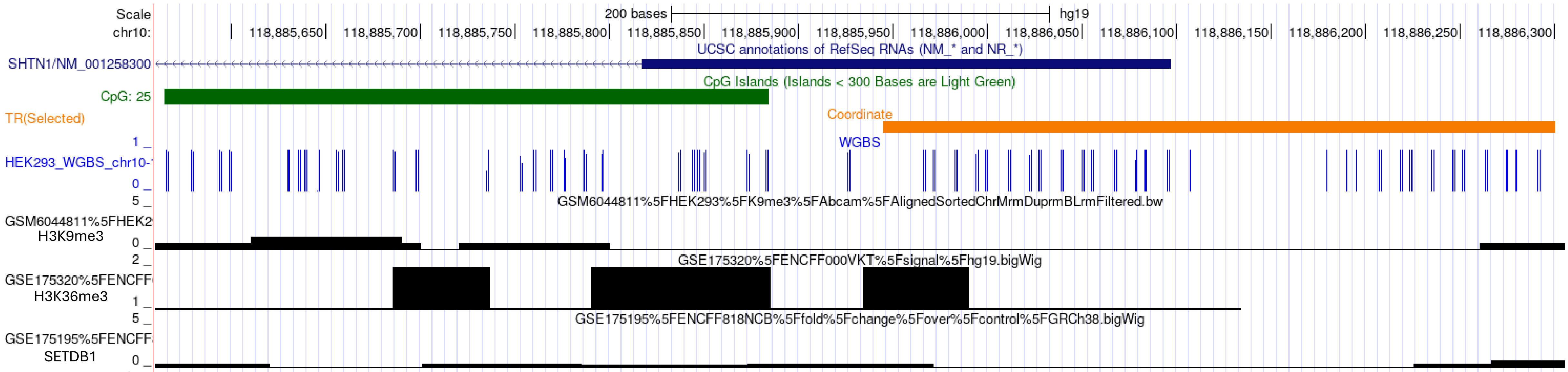
Scale
chr10:
200 bases
hg19
118,885,650
118,885,700
118,885,750
118,885,800
118,885,850
118,885,900
118,885,950
118,886,000
118,886,050
118,886,100
118,886,150
118,886,200
118,886,250
118,886,300
UCSC annotations of RefSeq RNAs (NM_* and NR_*)
SHTN1/NM_001258300
CpG Islands (Islands < 300 Bases are Light Green)
CpG: 25
TR(Selected)
Coordinate
WGBS
HEK293_WGBS_chr10-
GSM6044811%5FHEK293%5FK9me3%5FAbcam%5FAlignedSortedChrMrmDuprmBLrmFiltered.bw
GSM6044811%5FHEK2
H3K9me3
GSE175320%5FENCFF000VKT%5Fsignal%5Fhg19.bigWig
GSE175320%5FENCFF
H3K36me3
GSE175195%5FENCFF818NCB%5Ffold%5Fchange%5Fover%5Fcontrol%5FGRCh38.bigWig
GSE175195%5FENCFF
SETDB1

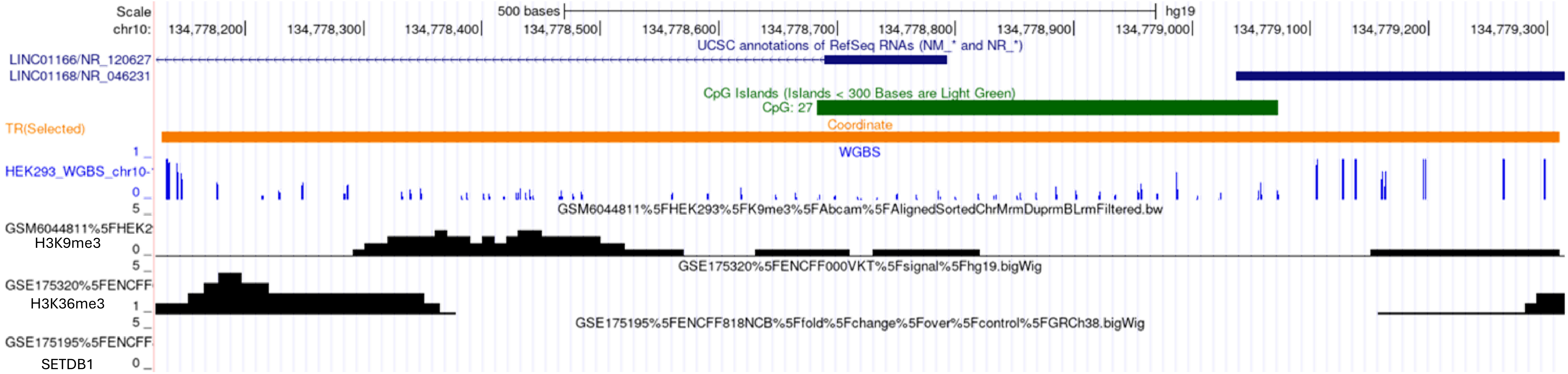
Scale
chr10:
500 bases
hg19
134,778,200
134,778,300
134,778,400
134,778,500
134,778,600
134,778,700
134,778,800
134,778,900
134,779,000
134,779,100
134,779,200
134,779,300
UCSC annotations of RefSeq RNAs (NM_* and NR_*)
LINC01166/NR_120627
LINC01168/NR_046231
CpG Islands (Islands < 300 Bases are Light Green)
CpG: 27
TR(Selected)
Coordinate
WGBS
HEK293_WGBS_chr10-
GSM6044811%5FHEK293%5FK9me3%5FAbcam%5FAlignedSortedChrMrmDuprmBLrmFiltered.bw
GSM6044811%5FHEK2
H3K9me3
GSE175320%5FENCFF000VKT%5Fsignal%5Fhg19.bigWig
GSE175320%5FENCFF
H3K36me3
GSE175195%5FENCFF818NCB%5Ffold%5Fchange%5Fover%5Fcontrol%5FGRCh38.bigWig
GSE175195%5FENCFF
SETDB1

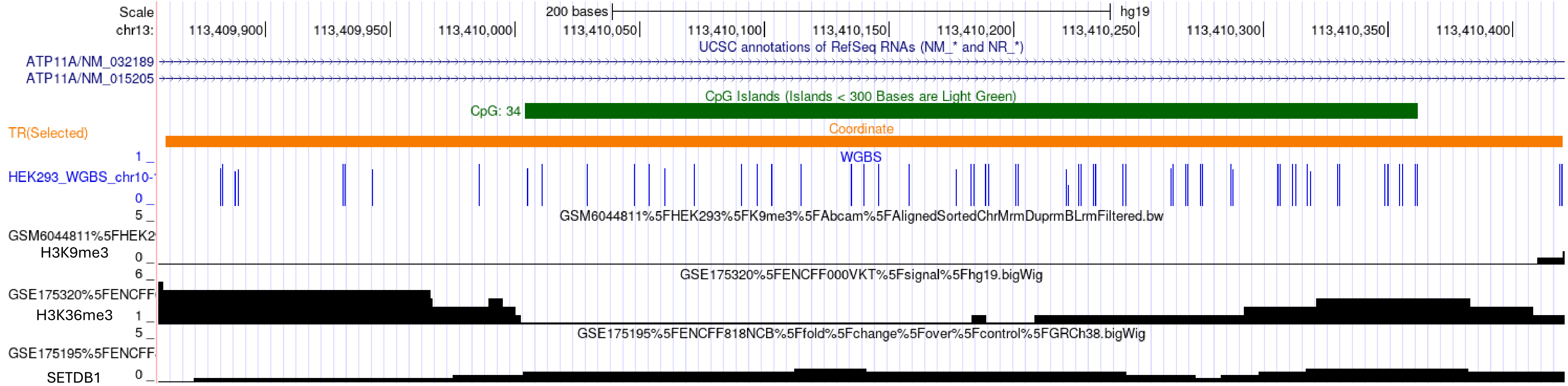
Scale
chr13:
200 bases
hg19
113,409,900
113,409,950
113,410,000
113,410,050
113,410,100
113,410,150
113,410,200
113,410,250
113,410,300
113,410,350
113,410,400
UCSC annotations of RefSeq RNAs (NM_* and NR_*)
ATP11A/NM_032189
ATP11A/NM_015205
CpG Islands (Islands < 300 Bases are Light Green)
CpG: 34
TR(Selected)
Coordinate
WGBS
HEK293_WGBS_chr10-
GSM6044811%5FHEK293%5FK9me3%5FAbcam%5FAlignedSortedChrMrmDuprmBLrmFiltered.bw
GSM6044811%5FHEK2
H3K9me3
GSE175320%5FENCFF000VKT%5Fsignal%5Fhg19.bigWig
GSE175320%5FENCFF
H3K36me3
GSE175195%5FENCFF818NCB%5Ffold%5Fchange%5Fover%5Fcontrol%5FGRCh38.bigWig
GSE175195%5FENCFF
SETDB1

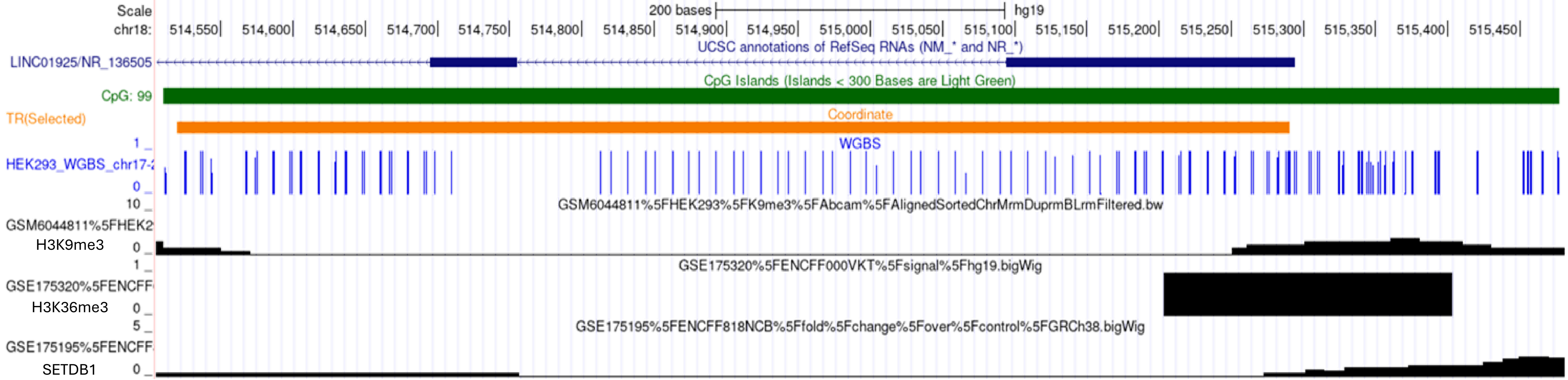
Scale
chr18:
200 bases
hg19
514,550
514,600
514,650
514,700
514,750
514,800
514,850
514,900
514,950
515,000
515,050
515,100
515,150
515,200
515,250
515,300
515,350
515,400
515,450
UCSC annotations of RefSeq RNAs (NM_* and NR_*)
LINC01925/NR_136505
CpG Islands (Islands < 300 Bases are Light Green)
CpG: 99
TR(Selected)
Coordinate
WGBS
HEK293_WGBS_chr17-
GSM6044811%5FHEK293%5FK9me3%5FAbcam%5FAlignedSortedChrMrmDuprmBLrmFiltered.bw
GSM6044811%5FHEK2
H3K9me3
GSE175320%5FENCFF000VKT%5Fsignal%5Fhg19.bigWig
GSE175320%5FENCFF
H3K36me3
GSE175195%5FENCFF818NCB%5Ffold%5Fchange%5Fover%5Fcontrol%5FGRCh38.bigWig
GSE175195%5FENCFF
SETDB1

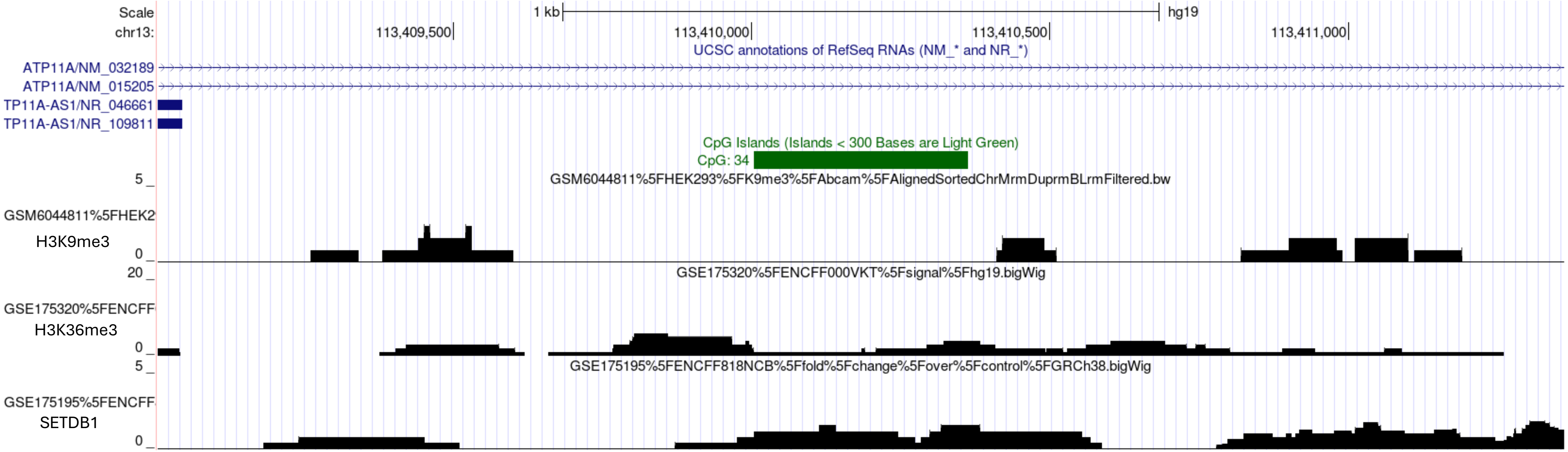

Scale
chr13:
1 kb
hg19
113,409,500
113,410,000
113,410,500
113,411,000
UCSC annotations of RefSeq RNAs (NM_* and NR_*)
ATP11A/NM_032189
ATP11A/NM_015205
TP11A-AS1/NR_046661
TP11A-AS1/NR_109811
CpG Islands (Islands < 300 Bases are Light Green)
CpG: 34
GSM6044811%5FHEK293%5FK9me3%5FAbcam%5FAlignedSortedChrMrmDuprmBLrmFiltered.bw
GSM6044811%5FHEK2
H3K9me3
GSE175320%5FENCFF000VKT%5Fsignal%5Fhg19.bigWig
GSE175320%5FENCFF
H3K36me3
GSE175195%5FENCFF818NCB%5Ffold%5Fchange%5Fover%5Fcontrol%5FGRCh38.bigWig
GSE175195%5FENCFF
SETDB1

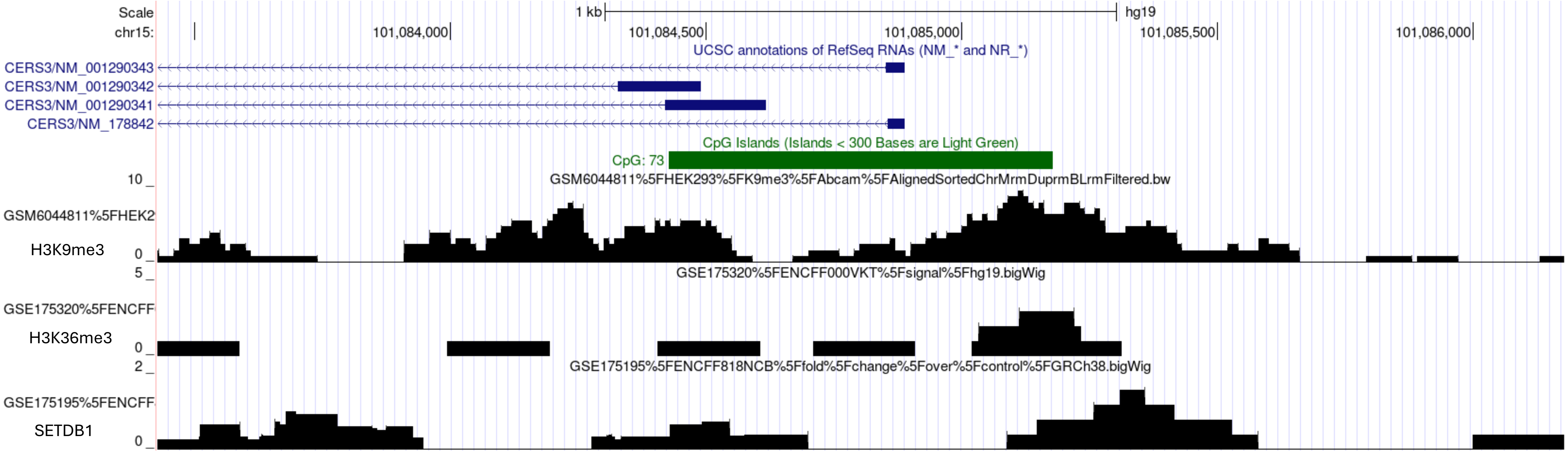

Scale
chr15:
1 kb
hg19
101,084,000
101,084,500
101,085,000
101,085,500
101,086,000
UCSC annotations of RefSeq RNAs (NM_* and NR_*)
CERS3/NM_001290343
CERS3/NM_001290342
CERS3/NM_001290341
CERS3/NM_178842
CpG Islands (Islands < 300 Bases are Light Green)
CpG: 73
GSM6044811%5FHEK293%5FK9me3%5FAbcam%5FAlignedSortedChrMrmDuprmBLrmFiltered.bw
GSM6044811%5FHEK2
H3K9me3
GSE175320%5FENCFF000VKT%5Fsignal%5Fhg19.bigWig
GSE175320%5FENCFF
H3K36me3
GSE175195%5FENCFF818NCB%5Ffold%5Fchange%5Fover%5Fcontrol%5FGRCh38.bigWig
GSE175195%5FENCFF
SETDB1

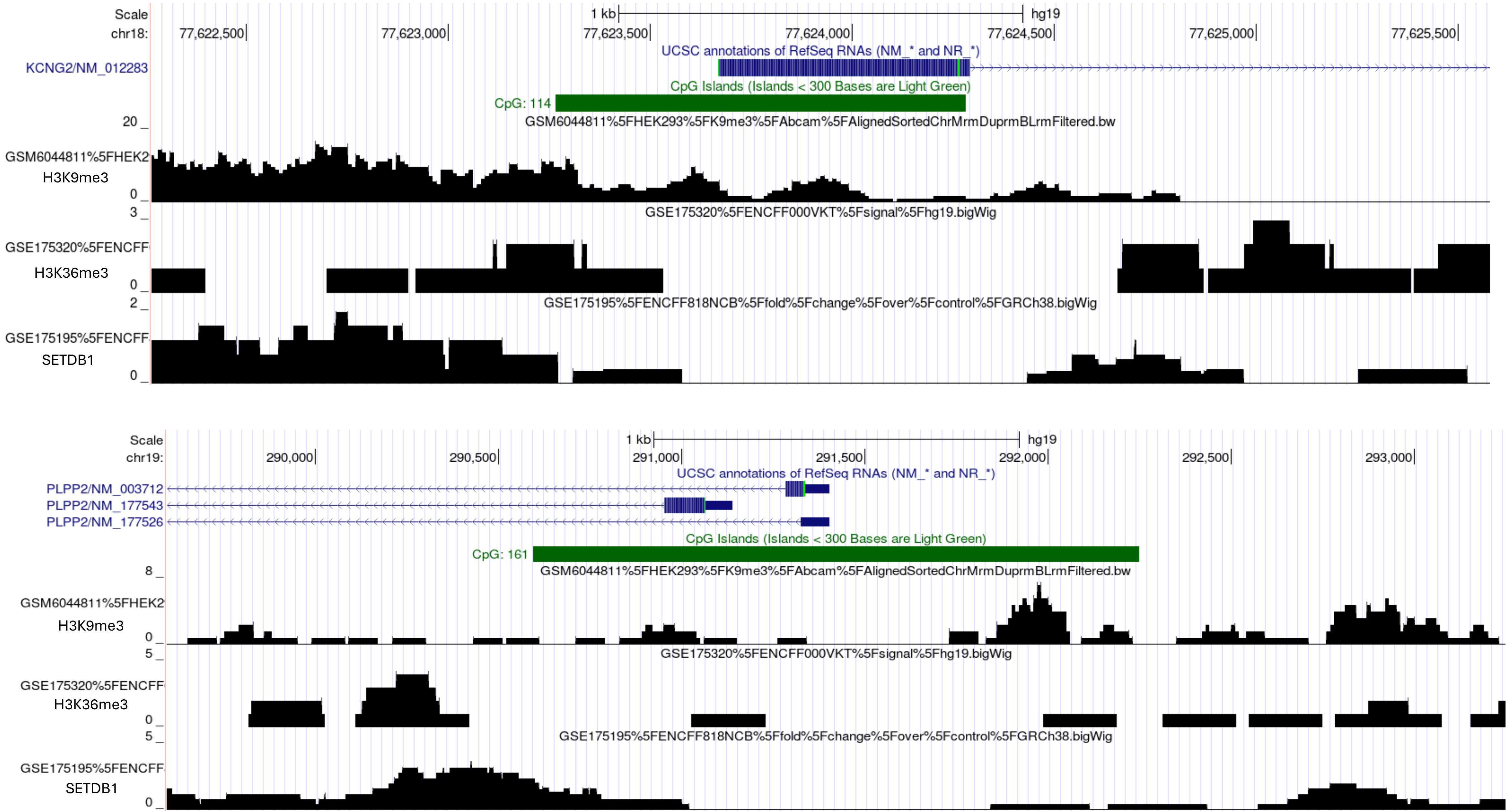
Scale
chr18:
1 kb
hg19
77,622,500
77,623,000
77,623,500
77,624,000
77,624,500
77,625,000
77,625,500
UCSC annotations of RefSeq RNAs (NM_* and NR_*)
KCNG2/NM_012283
CpG Islands (Islands < 300 Bases are Light Green)
CpG: 114
GSM6044811%5FHEK293%5FK9me3%5FAbcam%5FAlignedSortedChrMrmDuprmBLrmFiltered.bw
20
GSM6044811%5FHEK2
H3K9me3
0
GSE175320%5FENCFF000VKT%5Fsignal%5Fhg19.bigWig
3
GSE175320%5FENCFF
H3K36me3
0
GSE175195%5FENCFF818NCB%5Ffold%5Fchange%5Fover%5Fcontrol%5FGRCh38.bigWig
2
GSE175195%5FENCFF
SETDB1
0
Scale
chr19:
1 kb
hg19
290,000
290,500
291,000
291,500
292,000
292,500
293,000
UCSC annotations of RefSeq RNAs (NM_* and NR_*)
PLPP2/NM_003712
PLPP2/NM_177543
PLPP2/NM_177526
CpG Islands (Islands < 300 Bases are Light Green)
CpG: 161
GSM6044811%5FHEK293%5FK9me3%5FAbcam%5FAlignedSortedChrMrmDuprmBLrmFiltered.bw
8
GSM6044811%5FHEK2
H3K9me3
0
GSE175320%5FENCFF000VKT%5Fsignal%5Fhg19.bigWig
5
GSE175320%5FENCFF
H3K36me3
0
GSE175195%5FENCFF818NCB%5Ffold%5Fchange%5Fover%5Fcontrol%5FGRCh38.bigWig
5
GSE175195%5FENCFF
SETDB1
0

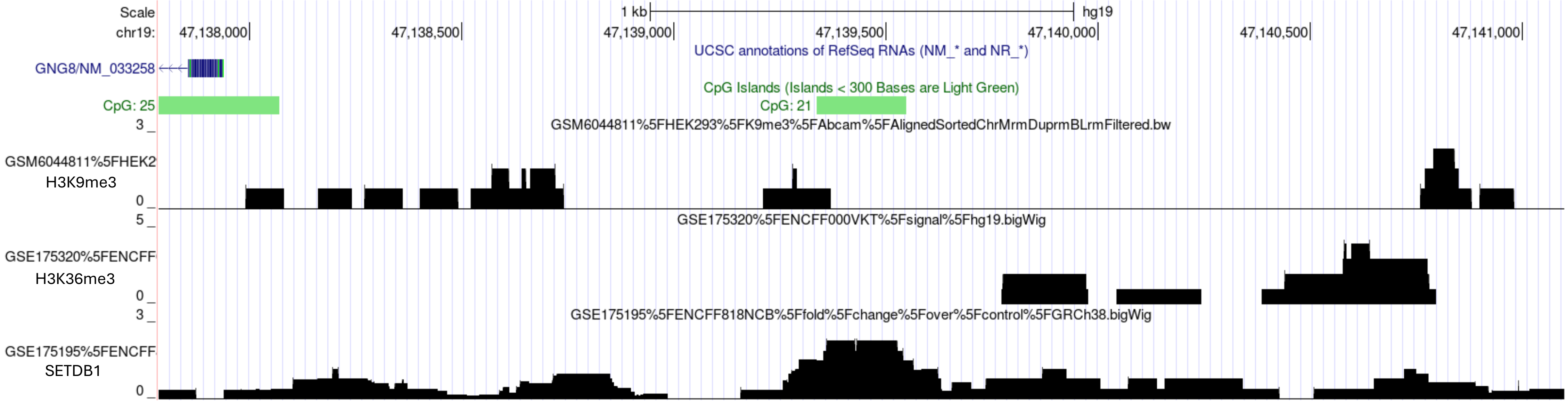

Scale
chr19:
1 kb
hg19
47,138,000
47,138,500
47,139,000
47,139,500
47,140,000
47,140,500
47,141,000
UCSC annotations of RefSeq RNAs (NM_* and NR_*)
GNG8/NM_033258
CpG Islands (Islands < 300 Bases are Light Green)
CpG: 25
CpG: 21
GSM6044811%5FHEK293%5FK9me3%5FAbcam%5FAlignedSortedChrMrmDuprmBLrmFiltered.bw
GSM6044811%5FHEK2
H3K9me3
GSE175320%5FENCFF000VKT%5Fsignal%5Fhg19.bigWig
GSE175320%5FENCFF
H3K36me3
GSE175195%5FENCFF818NCB%5Ffold%5Fchange%5Fover%5Fcontrol%5FGRCh38.bigWig
GSE175195%5FENCFF
SETDB1


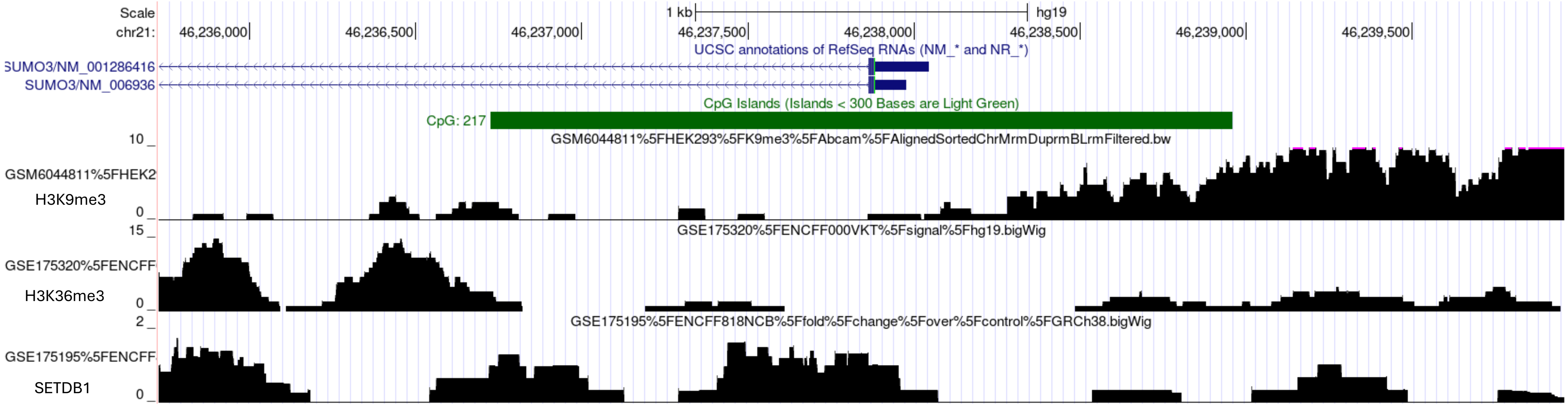

Scale
chr21:
1 kb
hg19
46,236,000
46,236,500
46,237,000
46,237,500
46,238,000
46,238,500
46,239,000
46,239,500
UCSC annotations of RefSeq RNAs (NM_* and NR_*)
SUMO3/NM_001286416
SUMO3/NM_006936
CpG Islands (Islands < 300 Bases are Light Green)
CpG: 217
GSM6044811%5FHEK293%5FK9me3%5FAbcam%5FAlignedSortedChrMrmDuprmBLrmFiltered.bw
GSM6044811%5FHEK2
H3K9me3
GSE175320%5FENCFF000VKT%5Fsignal%5Fhg19.bigWig
GSE175320%5FENCFF
H3K36me3
GSE175195%5FENCFF818NCB%5Ffold%5Fchange%5Fover%5Fcontrol%5FGRCh38.bigWig
GSE175195%5FENCFF
SETDB1

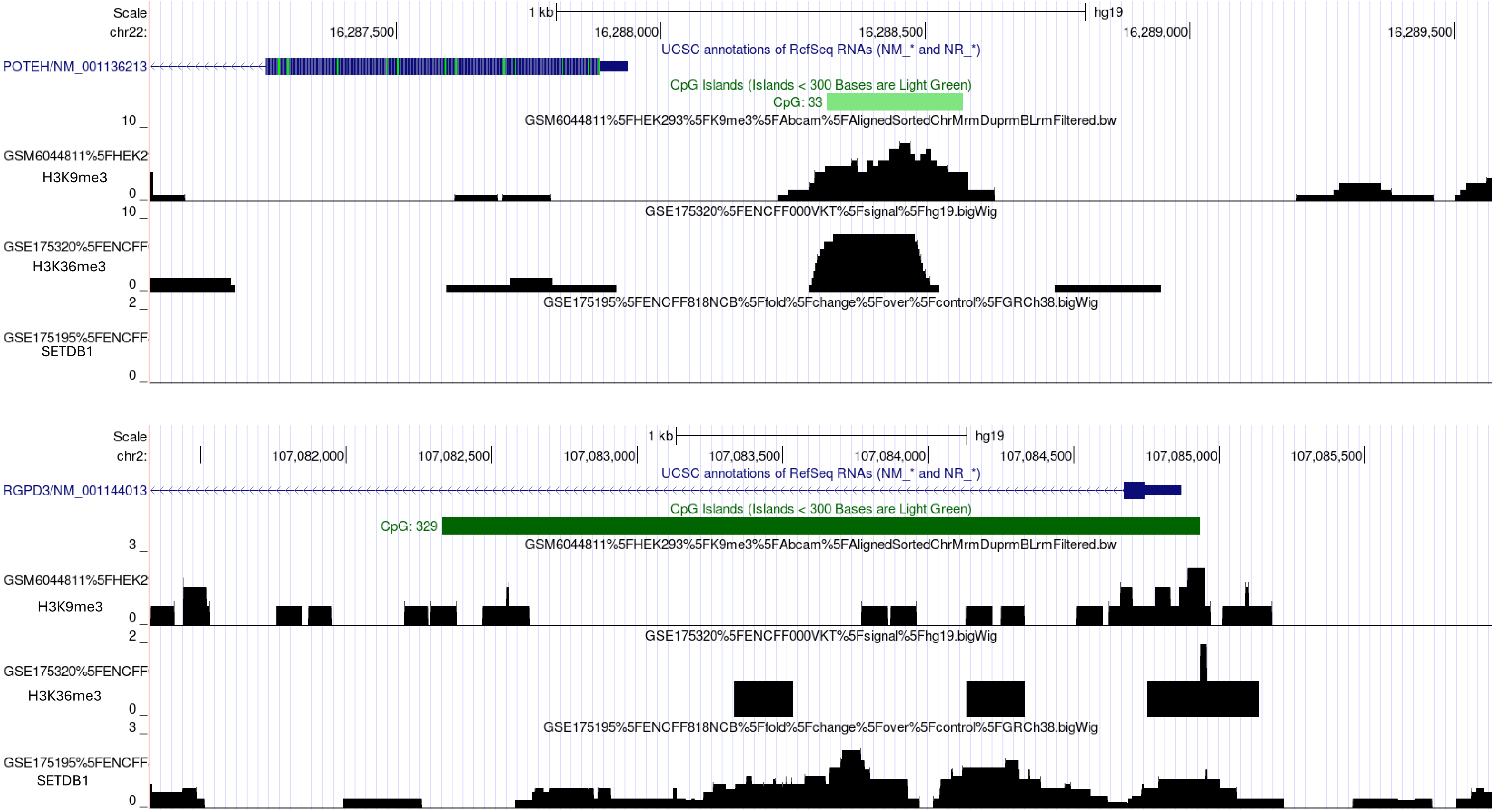

Scale
chr22:
1 kb
hg19
16,287,500
16,288,000
16,288,500
16,289,000
16,289,500
UCSC annotations of RefSeq RNAs (NM_* and NR_*)
POTEH/NM_001136213
CpG Islands (Islands < 300 Bases are Light Green)
CpG: 33
10 _
GSM6044811%5FHEK293%5FK9me3%5FAbcam%5FAlignedSortedChrMrmDuprmBLrmFiltered.bw
GSM6044811%5FHEK2
H3K9me3
0 _
10 _
GSE175320%5FENCFF000VKT%5Fsignal%5Fhg19.bigWig
GSE175320%5FENCFF
H3K36me3
0 _
2 _
GSE175195%5FENCFF818NCB%5Ffold%5Fchange%5Fover%5Fcontrol%5FGRCh38.bigWig
GSE175195%5FENCFF
SETDB1
0 _
Scale
chr2:
1 kb
hg19
107,082,000
107,082,500
107,083,000
107,083,500
107,084,000
107,084,500
107,085,000
107,085,500
UCSC annotations of RefSeq RNAs (NM_* and NR_*)
RGPD3/NM_001144013
CpG Islands (Islands < 300 Bases are Light Green)
CpG: 329
3 _
GSM6044811%5FHEK293%5FK9me3%5FAbcam%5FAlignedSortedChrMrmDuprmBLrmFiltered.bw
GSM6044811%5FHEK2
H3K9me3
0 _
2 _
GSE175320%5FENCFF000VKT%5Fsignal%5Fhg19.bigWig
GSE175320%5FENCFF
H3K36me3
0 _
3 _
GSE175195%5FENCFF818NCB%5Ffold%5Fchange%5Fover%5Fcontrol%5FGRCh38.bigWig
GSE175195%5FENCFF
SETDB1
0 _

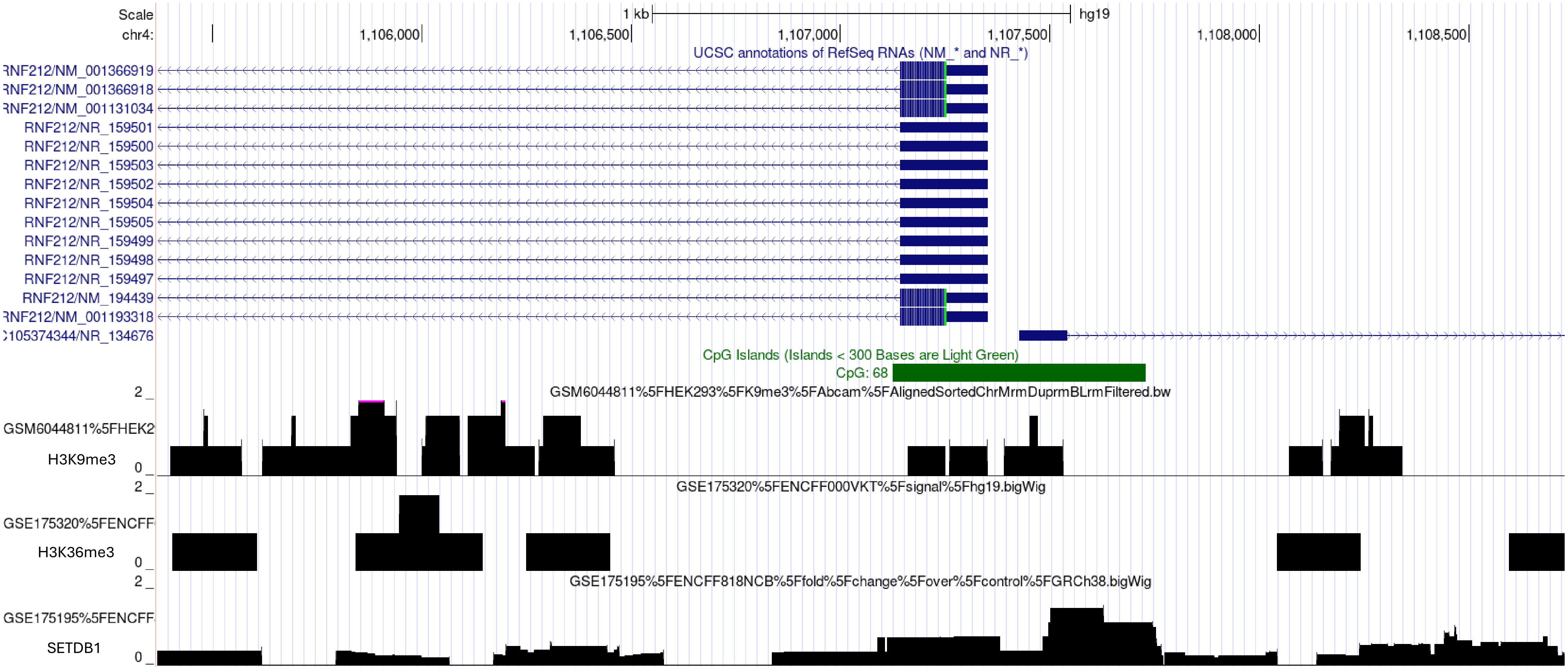
Scale
chr4:
1 kb
hg19
1,106,000
1,106,500
1,107,000
1,107,500
1,108,000
1,108,500
UCSC annotations of RefSeq RNAs (NM_* and NR_*)
RNF212/NM_001366919
RNF212/NM_001366918
RNF212/NM_001131034
RNF212/NR_159501
RNF212/NR_159500
RNF212/NR_159503
RNF212/NR_159502
RNF212/NR_159504
RNF212/NR_159505
RNF212/NR_159499
RNF212/NR_159498
RNF212/NR_159497
RNF212/NM_194439
RNF212/NM_001193318
105374344/NR_134676
CpG Islands (Islands < 300 Bases are Light Green)
CpG: 68
GSM6044811%5FHEK293%5FK9me3%5FAbcam%5FAlignedSortedChrMrmDuprmBLrmFiltered.bw
GSM6044811%5FHEK2
H3K9me3
GSE175320%5FENCFF000VKT%5Fsignal%5Fhg19.bigWig
GSE175320%5FENCFF
H3K36me3
GSE175195%5FENCFF818NCB%5Ffold%5Fchange%5Fover%5Fcontrol%5FGRCh38.bigWig
GSE175195%5FENCFF
SETDB1
2
0

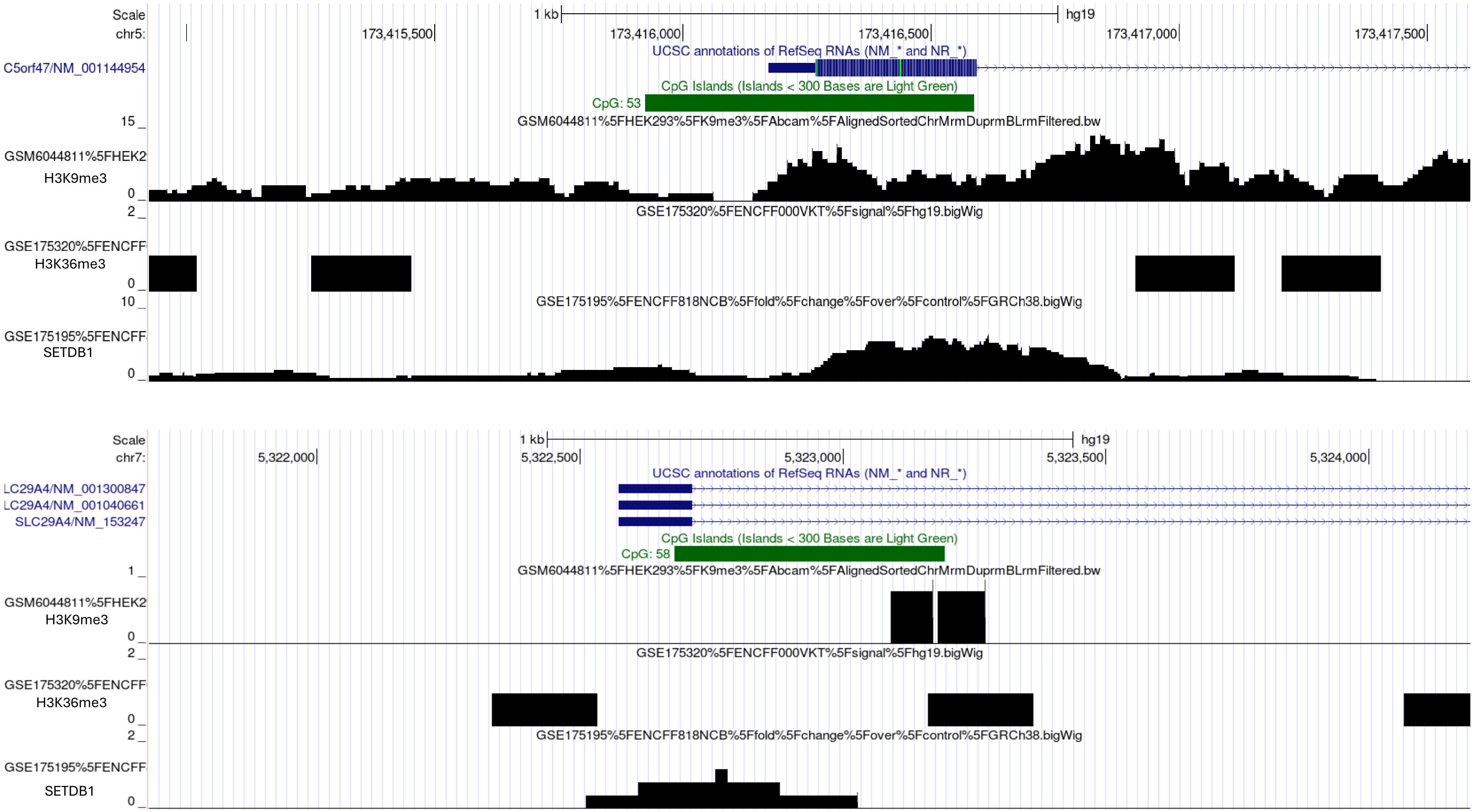

Scale
chr5:
1 kb
hg19
173,415,500
173,416,000
173,416,500
173,417,000
173,417,500
UCSC annotations of RefSeq RNAs (NM_* and NR_*)
C5orf47/NM_001144954
CpG Islands (Islands < 300 Bases are Light Green)
CpG: 53
15
GSM6044811%5FHEK293%5FK9me3%5FAbcam%5FAlignedSortedChrMrmDuprmBLrmFiltered.bw
GSM6044811%5FHEK2
H3K9me3
0
2
GSE175320%5FENCFF000VKT%5Fsignal%5Fhg19.bigWig
GSE175320%5FENCFF
H3K36me3
0
10
GSE175195%5FENCFF818NCB%5Ffold%5Fchange%5Fover%5Fcontrol%5FGRCh38.bigWig
GSE175195%5FENCFF
SETDB1
0
Scale
chr7:
1 kb
hg19
5,322,000
5,322,500
5,323,000
5,323,500
5,324,000
UCSC annotations of RefSeq RNAs (NM_* and NR_*)
LC29A4/NM_001300847
LC29A4/NM_001040661
SLC29A4/NM_153247
CpG Islands (Islands < 300 Bases are Light Green)
CpG: 58
1
GSM6044811%5FHEK293%5FK9me3%5FAbcam%5FAlignedSortedChrMrmDuprmBLrmFiltered.bw
GSM6044811%5FHEK2
H3K9me3
0
2
GSE175320%5FENCFF000VKT%5Fsignal%5Fhg19.bigWig
GSE175320%5FENCFF
H3K36me3
0
2
GSE175195%5FENCFF818NCB%5Ffold%5Fchange%5Fover%5Fcontrol%5FGRCh38.bigWig
GSE175195%5FENCFF
SETDB1
0

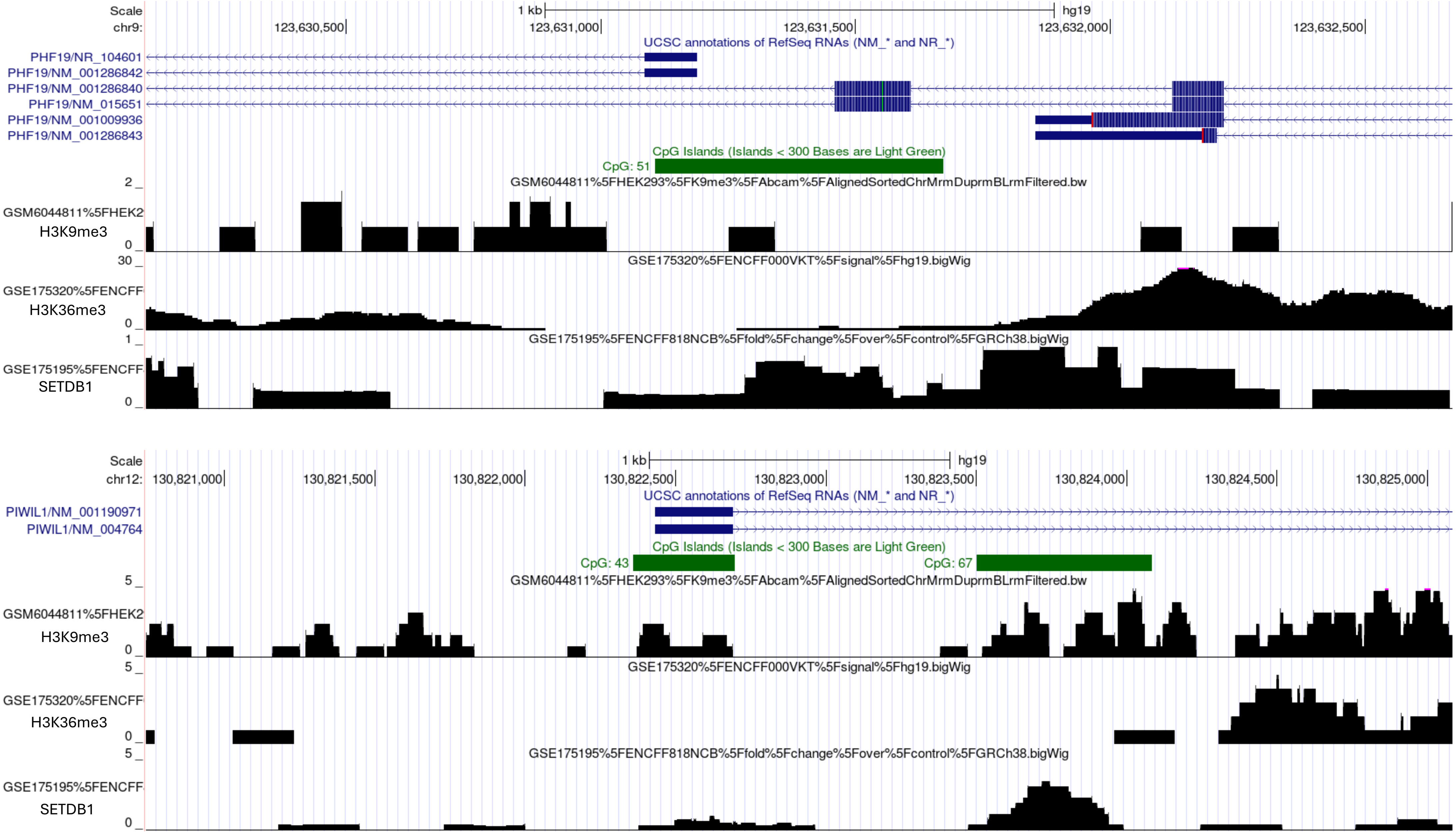
Scale
1 kb
hg19
chr9:
123,630,500
123,631,000
123,631,500
123,632,000
123,632,500
UCSC annotations of RefSeq RNAs (NM_* and NR_*)
PHF19/NR_104601
PHF19/NM_001286842
PHF19/NM_001286840
PHF19/NM_015651
PHF19/NM_001009936
PHF19/NM_001286843
CpG Islands (Islands < 300 Bases are Light Green)
CpG: 51
2
0
GSM6044811%5FHEK293%5FK9me3%5FAbcam%5FAlignedSortedChrMrmDuprmBLrmFiltered.bw
GSM6044811%5FHEK2
H3K9me3
30
0
GSE175320%5FENCFF000VKT%5Fsignal%5Fhg19.bigWig
GSE175320%5FENCFF
H3K36me3
1
0
GSE175195%5FENCFF818NCB%5Ffold%5Fchange%5Fover%5Fcontrol%5FGRCh38.bigWig
GSE175195%5FENCFF
SETDB1
Scale
1 kb
hg19
chr12:
130,821,000
130,821,500
130,822,000
130,822,500
130,823,000
130,823,500
130,824,000
130,824,500
130,825,000
UCSC annotations of RefSeq RNAs (NM_* and NR_*)
PIWIL1/NM_001190971
PIWIL1/NM_004764
CpG Islands (Islands < 300 Bases are Light Green)
CpG: 43
CpG: 67
5
0
GSM6044811%5FHEK293%5FK9me3%5FAbcam%5FAlignedSortedChrMrmDuprmBLrmFiltered.bw
GSM6044811%5FHEK2
H3K9me3
5
0
GSE175320%5FENCFF000VKT%5Fsignal%5Fhg19.bigWig
GSE175320%5FENCFF
H3K36me3
5
0
GSE175195%5FENCFF818NCB%5Ffold%5Fchange%5Fover%5Fcontrol%5FGRCh38.bigWig
GSE175195%5FENCFF
SETDB1

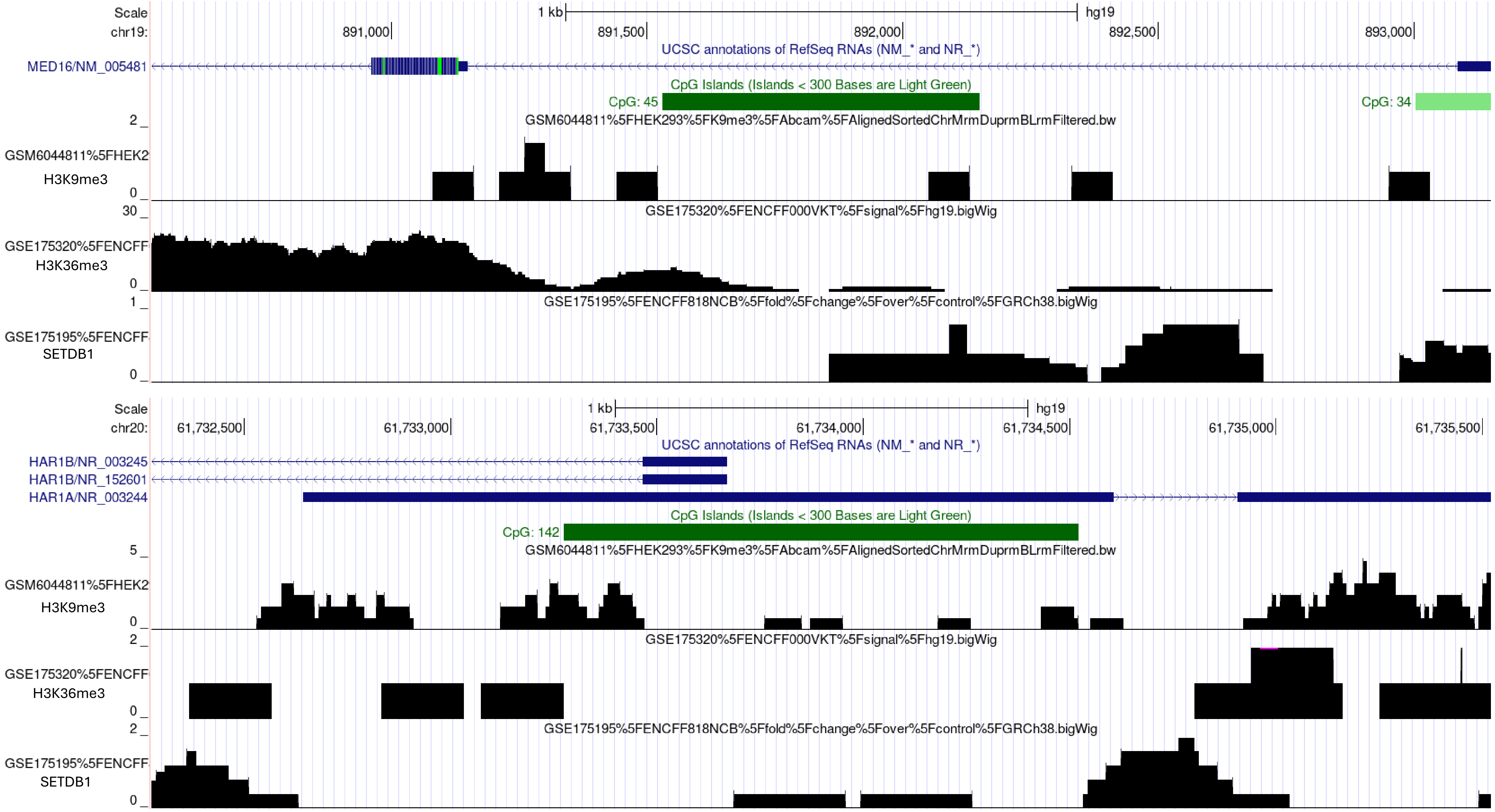
Scale
1 kb
hg19
chr19:
891,000
891,500
892,000
892,500
893,000
UCSC annotations of RefSeq RNAs (NM_* and NR_*)
MED16/NM_005481
CpG Islands (Islands < 300 Bases are Light Green)
CpG: 45
CpG: 34
GSM6044811%5FHEK293%5FK9me3%5FAbcam%5FAlignedSortedChrMrmDuprmBLrmFiltered.bw
GSM6044811%5FHEK2
H3K9me3
GSE175320%5FENCFF000VKT%5Fsignal%5Fhg19.bigWig
GSE175320%5FENCFF
H3K36me3
GSE175195%5FENCFF818NCB%5Ffold%5Fchange%5Fover%5Fcontrol%5FGRCh38.bigWig
GSE175195%5FENCFF
SETDB1
Scale
1 kb
hg19
chr20:
61,732,500
61,733,000
61,733,500
61,734,000
61,734,500
61,735,000
61,735,500
UCSC annotations of RefSeq RNAs (NM_* and NR_*)
HAR1B/NR_003245
HAR1B/NR_152601
HAR1A/NR_003244
CpG Islands (Islands < 300 Bases are Light Green)
CpG: 142
GSM6044811%5FHEK293%5FK9me3%5FAbcam%5FAlignedSortedChrMrmDuprmBLrmFiltered.bw
GSM6044811%5FHEK2
H3K9me3
GSE175320%5FENCFF000VKT%5Fsignal%5Fhg19.bigWig
GSE175320%5FENCFF
H3K36me3
GSE175195%5FENCFF818NCB%5Ffold%5Fchange%5Fover%5Fcontrol%5FGRCh38.bigWig
GSE175195%5FENCFF
SETDB1

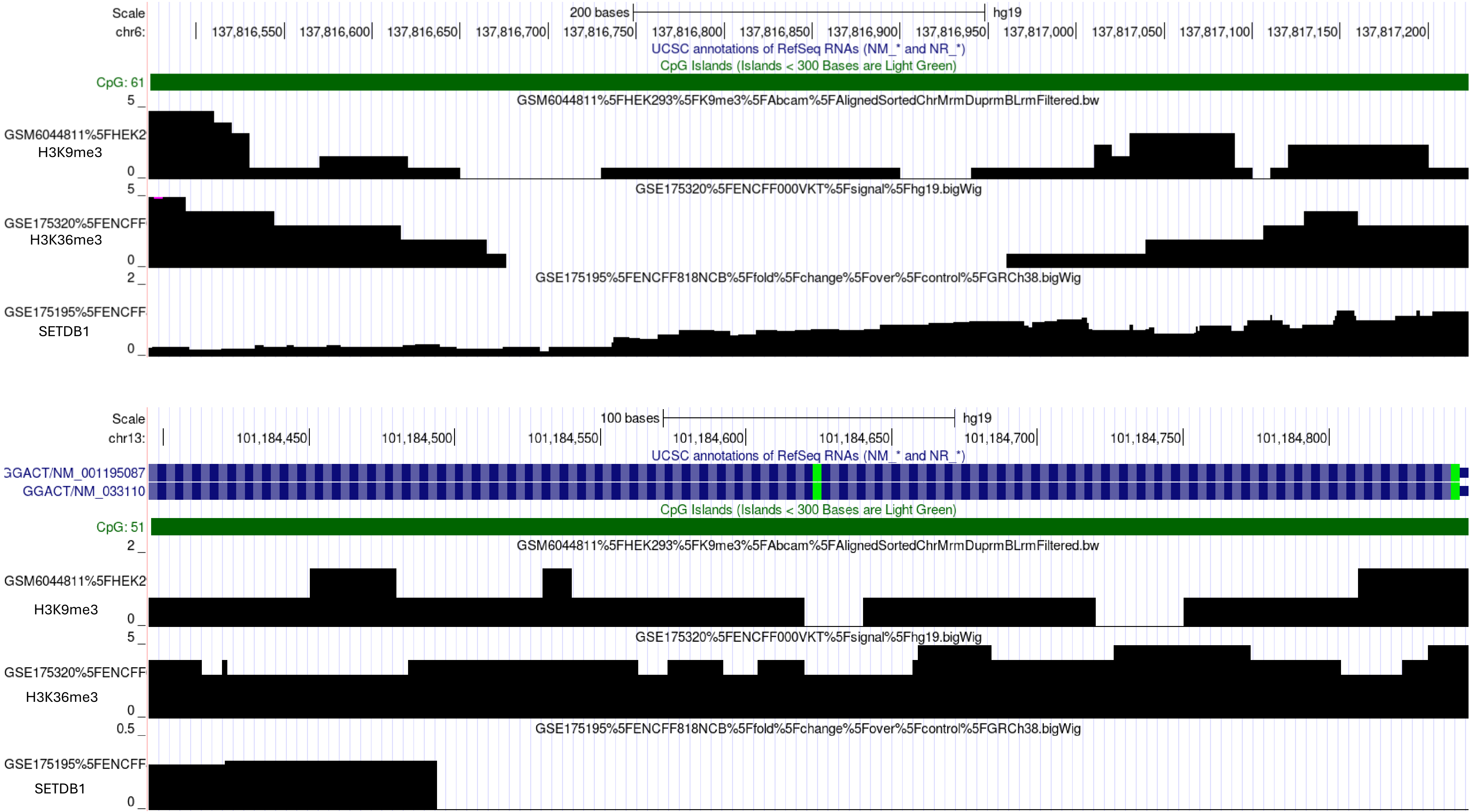
Scale
chr6:
200 bases
hg19
137,816,550
137,816,600
137,816,650
137,816,700
137,816,750
137,816,800
137,816,850
137,816,900
137,816,950
137,817,000
137,817,050
137,817,100
137,817,150
137,817,200
UCSC annotations of RefSeq RNAs (NM_* and NR_*)
CpG Islands (Islands < 300 Bases are Light Green)
CpG: 61
GSM6044811%5FHEK293%5FK9me3%5FAbcam%5FAlignedSortedChrMrmDuprmBLrmFiltered.bw
GSM6044811%5FHEK2
H3K9me3
GSE175320%5FENCFF000VKT%5Fsignal%5Fhg19.bigWig
GSE175320%5FENCFF
H3K36me3
GSE175195%5FENCFF818NCB%5Ffold%5Fchange%5Fover%5Fcontrol%5FGRCh38.bigWig
GSE175195%5FENCFF
SETDB1
Scale
chr13:
100 bases
hg19
101,184,450
101,184,500
101,184,550
101,184,600
101,184,650
101,184,700
101,184,750
101,184,800
UCSC annotations of RefSeq RNAs (NM_* and NR_*)
GGACT/NM_001195087
GGACT/NM_033110
CpG Islands (Islands < 300 Bases are Light Green)
CpG: 51
GSM6044811%5FHEK293%5FK9me3%5FAbcam%5FAlignedSortedChrMrmDuprmBLrmFiltered.bw
GSM6044811%5FHEK2
H3K9me3
GSE175320%5FENCFF000VKT%5Fsignal%5Fhg19.bigWig
GSE175320%5FENCFF
H3K36me3
GSE175195%5FENCFF818NCB%5Ffold%5Fchange%5Fover%5Fcontrol%5FGRCh38.bigWig
GSE175195%5FENCFF
SETDB1

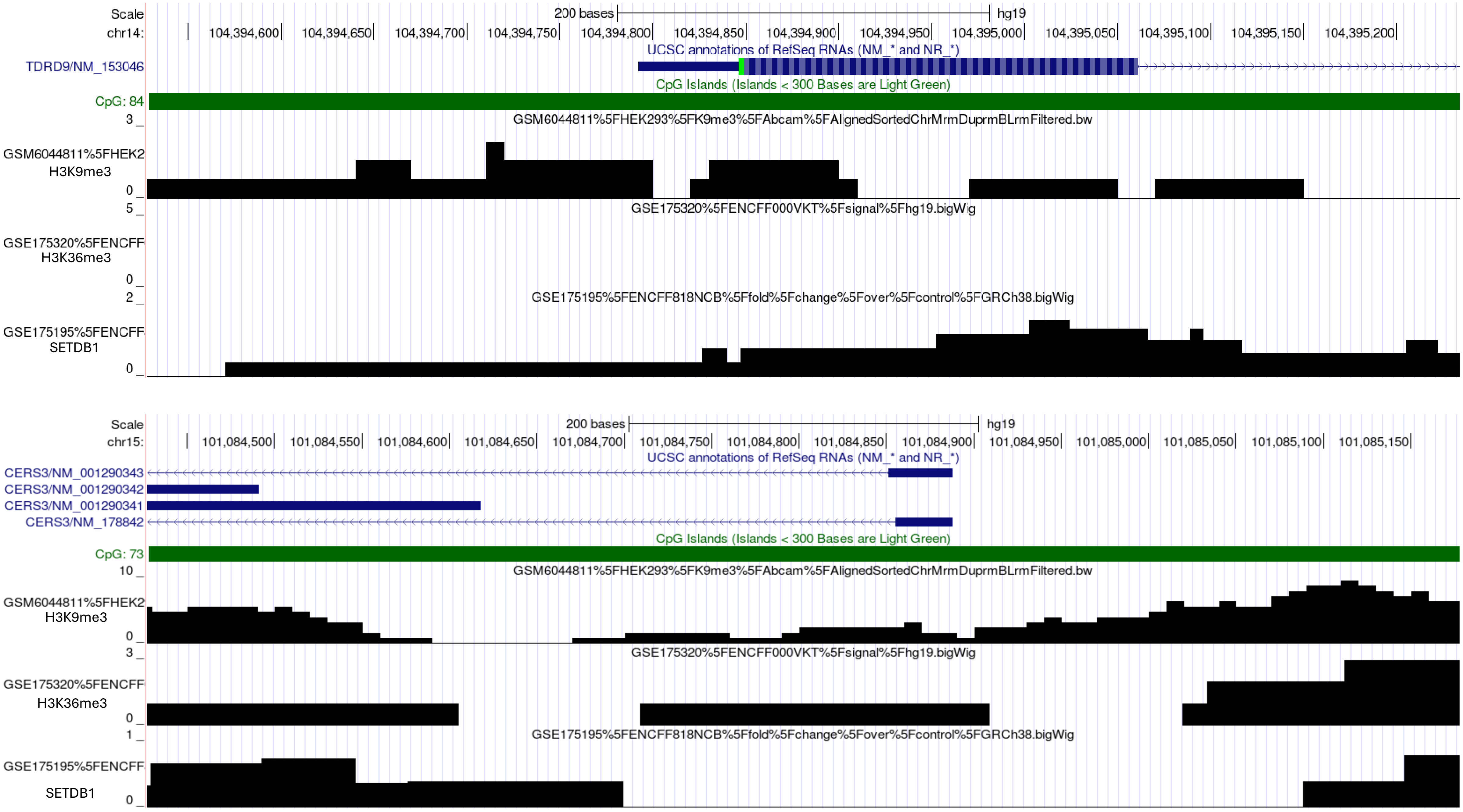
Scale
chr14:
200 bases
hg19
104,394,600
104,394,650
104,394,700
104,394,750
104,394,800
104,394,850
104,394,900
104,394,950
104,395,000
104,395,050
104,395,100
104,395,150
104,395,200
UCSC annotations of RefSeq RNAs (NM_* and NR_*)
TDRD9/NM_153046
CpG Islands (Islands < 300 Bases are Light Green)
CpG: 84
GSM6044811%5FHEK293%5FK9me3%5FAbcam%5FAlignedSortedChrMrmDuprmBLrmFiltered.bw
GSM6044811%5FHEK2
H3K9me3
GSE175320%5FENCFF000VKT%5Fsignal%5Fhg19.bigWig
GSE175320%5FENCFF
H3K36me3
GSE175195%5FENCFF818NCB%5Ffold%5Fchange%5Fover%5Fcontrol%5FGRCh38.bigWig
GSE175195%5FENCFF
SETDB1
Scale
chr15:
200 bases
hg19
101,084,500
101,084,550
101,084,600
101,084,650
101,084,700
101,084,750
101,084,800
101,084,850
101,084,900
101,084,950
101,085,000
101,085,050
101,085,100
101,085,150
UCSC annotations of RefSeq RNAs (NM_* and NR_*)
CERS3/NM_001290343
CERS3/NM_001290342
CERS3/NM_001290341
CERS3/NM_178842
CpG Islands (Islands < 300 Bases are Light Green)
CpG: 73
GSM6044811%5FHEK293%5FK9me3%5FAbcam%5FAlignedSortedChrMrmDuprmBLrmFiltered.bw
GSM6044811%5FHEK2
H3K9me3
GSE175320%5FENCFF000VKT%5Fsignal%5Fhg19.bigWig
GSE175320%5FENCFF
H3K36me3
GSE175195%5FENCFF818NCB%5Ffold%5Fchange%5Fover%5Fcontrol%5FGRCh38.bigWig
GSE175195%5FENCFF
SETDB1

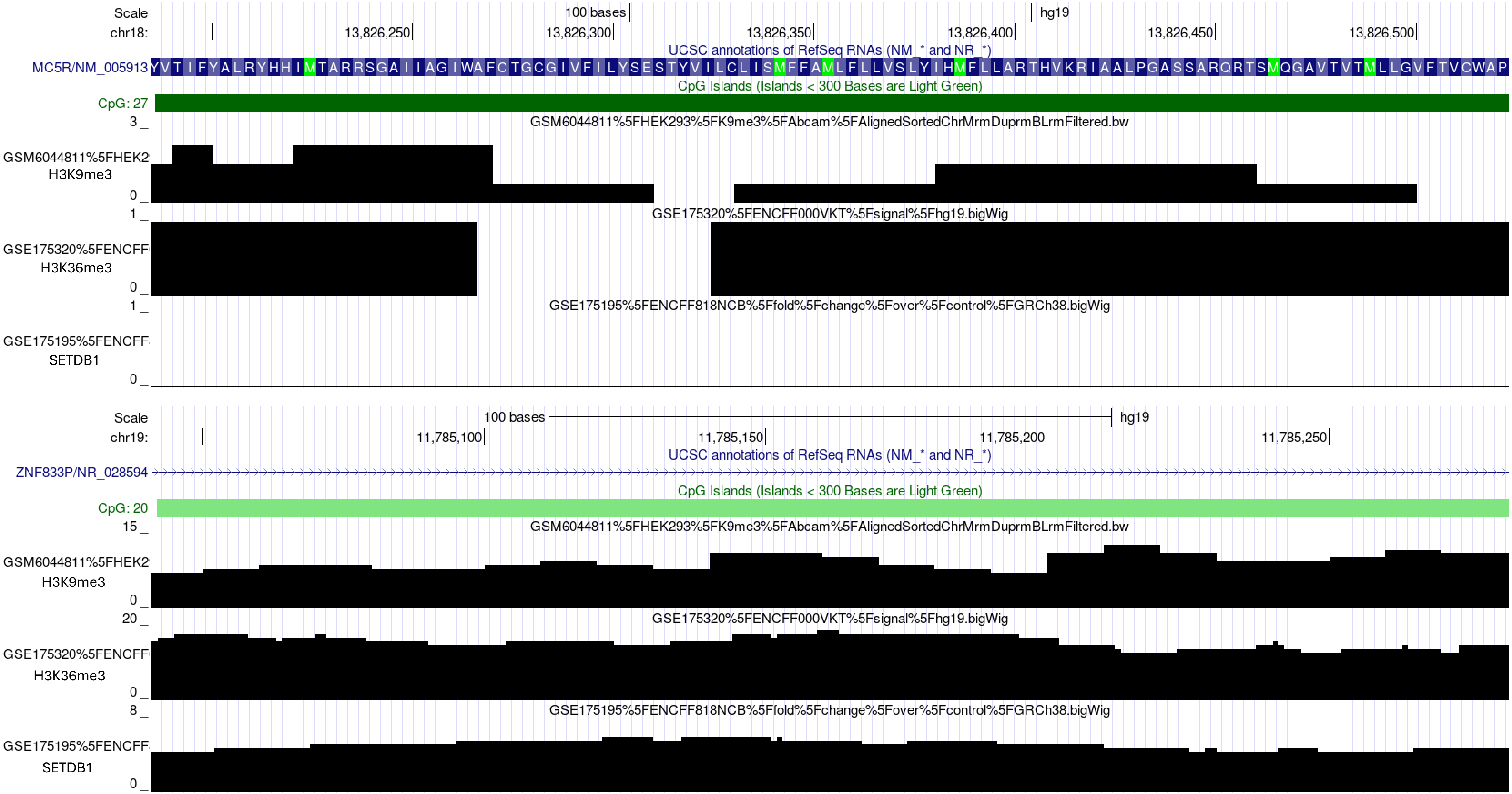
Scale
chr18:
100 bases
hg19
13,826,250
13,826,300
13,826,350
13,826,400
13,826,450
13,826,500
UCSC annotations of RefSeq RNAs (NM_* and NR_*)
MC5R/NM_005913
YVTIFYALRYHHIMTARRSGAIIAGIWAFCTGCGIVFILYSESTYVILCLISMFFAMLFLLVSLYIHMFLLARTHVKRIAALPGASSARQRTSMQGAVTVTMLLGVFTVCWAP
CpG Islands (Islands < 300 Bases are Light Green)
CpG: 27
GSM6044811%5FHEK293%5FK9me3%5FAbcam%5FAlignedSortedChrMrmDuprmBLrmFiltered.bw
GSM6044811%5FHEK2
H3K9me3
3
0
GSE175320%5FENCFF000VKT%5Fsignal%5Fhg19.bigWig
GSE175320%5FENCFF
H3K36me3
1
0
GSE175195%5FENCFF818NCB%5Ffold%5Fchange%5Fover%5Fcontrol%5FGRCh38.bigWig
GSE175195%5FENCFF
SETDB1
1
0
Scale
chr19:
100 bases
hg19
11,785,100
11,785,150
11,785,200
11,785,250
UCSC annotations of RefSeq RNAs (NM_* and NR_*)
ZNF833P/NR_028594
CpG Islands (Islands < 300 Bases are Light Green)
CpG: 20
GSM6044811%5FHEK293%5FK9me3%5FAbcam%5FAlignedSortedChrMrmDuprmBLrmFiltered.bw
GSM6044811%5FHEK2
H3K9me3
15
0
GSE175320%5FENCFF000VKT%5Fsignal%5Fhg19.bigWig
GSE175320%5FENCFF
H3K36me3
20
0
GSE175195%5FENCFF818NCB%5Ffold%5Fchange%5Fover%5Fcontrol%5FGRCh38.bigWig
GSE175195%5FENCFF
SETDB1
8
0

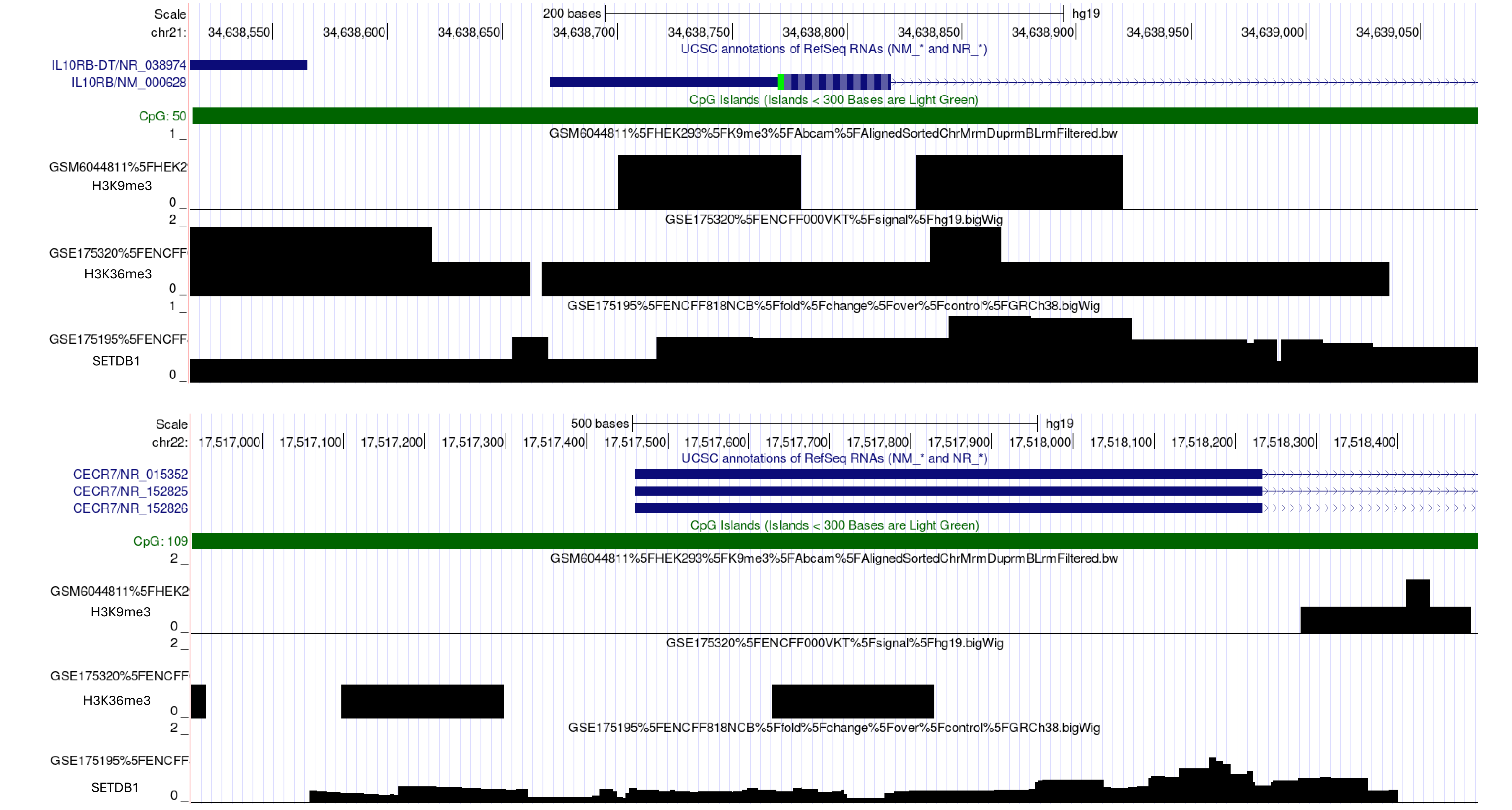

Scale
200 bases
hg19
chr21:
34,638,550
34,638,600
34,638,650
34,638,700
34,638,750
34,638,800
34,638,850
34,638,900
34,638,950
34,639,000
34,639,050
UCSC annotations of RefSeq RNAs (NM_* and NR_*)
IL10RB-DT/NR_038974
IL10RB/NM_000628
CpG Islands (Islands < 300 Bases are Light Green)
CpG: 50
GSM6044811%5FHEK293%5FK9me3%5FAbcam%5FAlignedSortedChrMrmDuprmBLrmFiltered.bw
GSM6044811%5FHEK2
H3K9me3
GSE175320%5FENCFF000VKT%5Fsignal%5Fhg19.bigWig
GSE175320%5FENCFF
H3K36me3
GSE175195%5FENCFF818NCB%5Ffold%5Fchange%5Fover%5Fcontrol%5FGRCh38.bigWig
GSE175195%5FENCFF
SETDB1
Scale
500 bases
hg19
chr22:
17,517,000
17,517,100
17,517,200
17,517,300
17,517,400
17,517,500
17,517,600
17,517,700
17,517,800
17,517,900
17,518,000
17,518,100
17,518,200
17,518,300
17,518,400
UCSC annotations of RefSeq RNAs (NM_* and NR_*)
CECR7/NR_015352
CECR7/NR_152825
CECR7/NR_152826
CpG Islands (Islands < 300 Bases are Light Green)
CpG: 109
GSM6044811%5FHEK293%5FK9me3%5FAbcam%5FAlignedSortedChrMrmDuprmBLrmFiltered.bw
GSM6044811%5FHEK2
H3K9me3
GSE175320%5FENCFF000VKT%5Fsignal%5Fhg19.bigWig
GSE175320%5FENCFF
H3K36me3
GSE175195%5FENCFF818NCB%5Ffold%5Fchange%5Fover%5Fcontrol%5FGRCh38.bigWig
GSE175195%5FENCFF
SETDB1

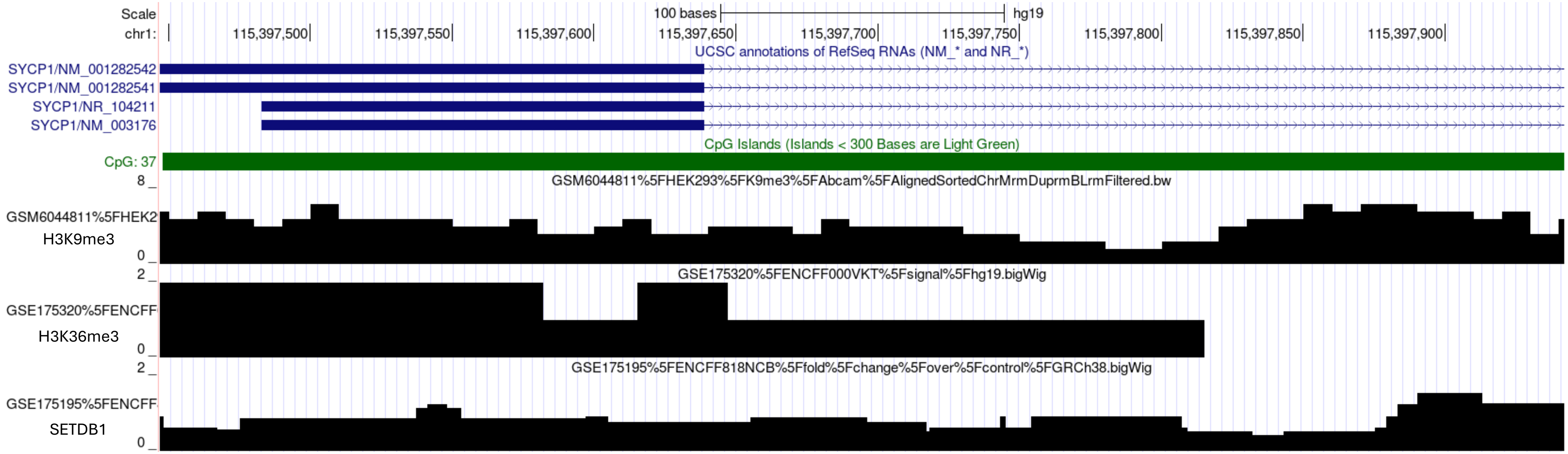
Scale
100 bases
hg19
chr1:
115,397,500
115,397,550
115,397,600
115,397,650
115,397,700
115,397,750
115,397,800
115,397,850
115,397,900
UCSC annotations of RefSeq RNAs (NM_* and NR_*)
SYCP1/NM_001282542
SYCP1/NM_001282541
SYCP1/NR_104211
SYCP1/NM_003176
CpG Islands (Islands < 300 Bases are Light Green)
CpG: 37
8
GSM6044811%5FHEK293%5FK9me3%5FAbcam%5FAlignedSortedChrMrmDuprmBLrmFiltered.bw
GSM6044811%5FHEK2
H3K9me3
0
2
GSE175320%5FENCFF000VKT%5Fsignal%5Fhg19.bigWig
GSE175320%5FENCFF
H3K36me3
0
2
GSE175195%5FENCFF818NCB%5Ffold%5Fchange%5Fover%5Fcontrol%5FGRCh38.bigWig
GSE175195%5FENCFF
SETDB1
0

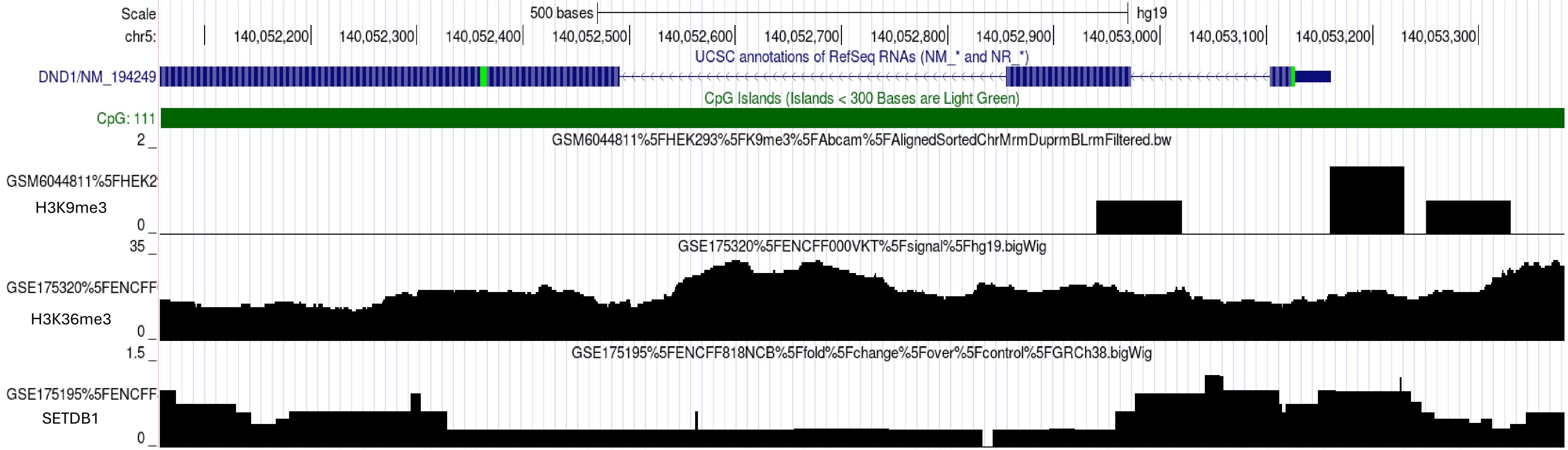
Scale
500 bases
hg19
chr5:
140,052,200
140,052,300
140,052,400
140,052,500
140,052,600
140,052,700
140,052,800
140,052,900
140,053,000
140,053,100
140,053,200
140,053,300
UCSC annotations of RefSeq RNAs (NM_* and NR_*)
DND1/NM_194249
CpG Islands (Islands < 300 Bases are Light Green)
CpG: 111
2
GSM6044811%5FHEK293%5FK9me3%5FAbcam%5FAlignedSortedChrMrmDuprmBLrmFiltered.bw
GSM6044811%5FHEK2
H3K9me3
0
35
GSE175320%5FENCFF000VKT%5Fsignal%5Fhg19.bigWig
GSE175320%5FENCFF
H3K36me3
0
1.5
GSE175195%5FENCFF818NCB%5Ffold%5Fchange%5Fover%5Fcontrol%5FGRCh38.bigWig
GSE175195%5FENCFF
SETDB1
0

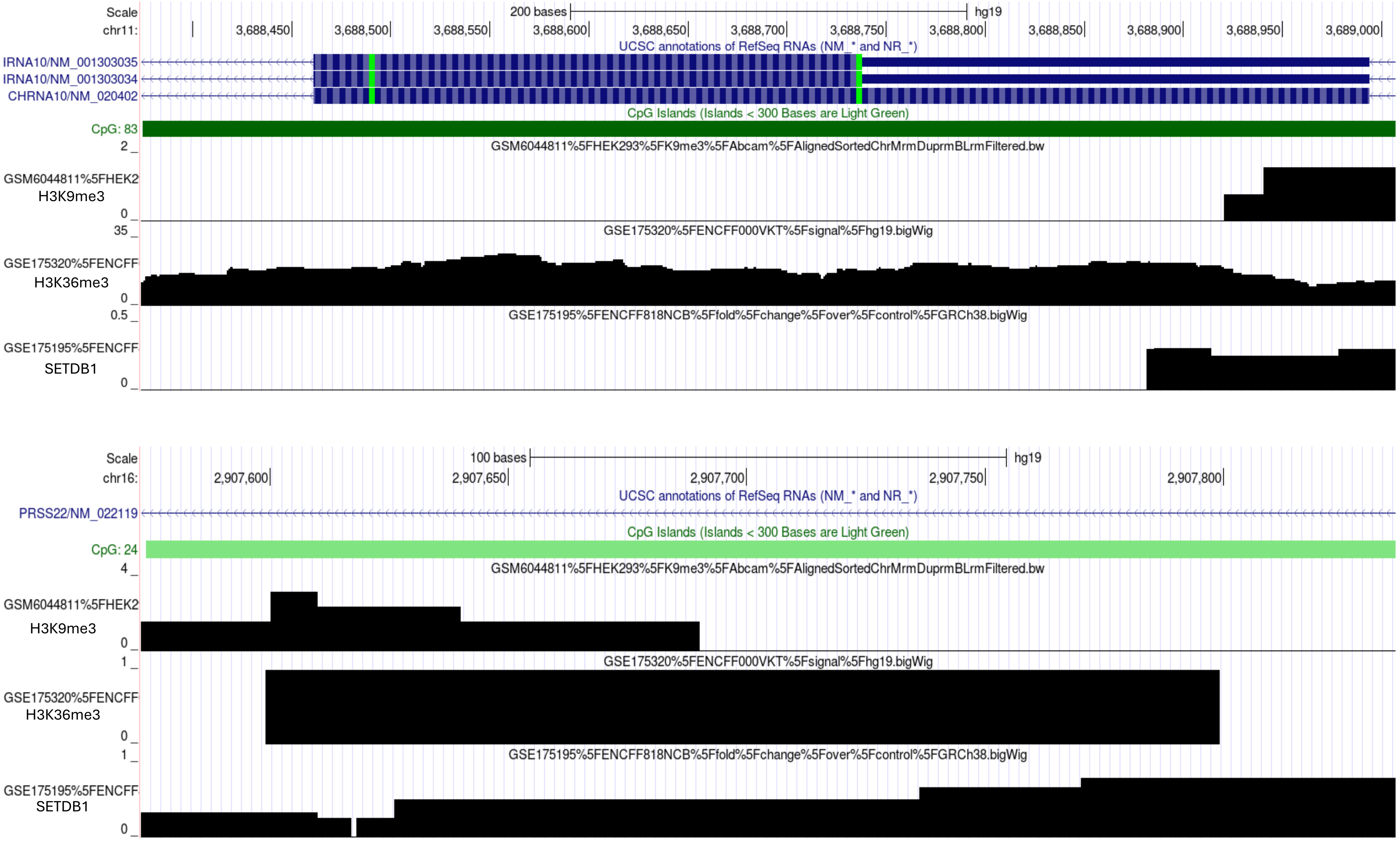

Scale
200 bases
hg19
chr11:
3,688,450
3,688,500
3,688,550
3,688,600
3,688,650
3,688,700
3,688,750
3,688,800
3,688,850
3,688,900
3,688,950
3,689,000
UCSC annotations of RefSeq RNAs (NM_* and NR_*)
IRNA10/NM_001303035
IRNA10/NM_001303034
CHRNA10/NM_020402
CpG Islands (Islands < 300 Bases are Light Green)
CpG: 83
GSM6044811%5FHEK293%5FK9me3%5FAbcam%5FAlignedSortedChrMrmDuprmBLrmFiltered.bw
GSM6044811%5FHEK2
H3K9me3
GSE175320%5FENCFF000VKT%5Fsignal%5Fhg19.bigWig
GSE175320%5FENCFF
H3K36me3
GSE175195%5FENCFF818NCB%5Ffold%5Fchange%5Fover%5Fcontrol%5FGRCh38.bigWig
GSE175195%5FENCFF
SETDB1
Scale
100 bases
hg19
chr16:
2,907,600
2,907,650
2,907,700
2,907,750
2,907,800
UCSC annotations of RefSeq RNAs (NM_* and NR_*)
PRSS22/NM_022119
CpG Islands (Islands < 300 Bases are Light Green)
CpG: 24
GSM6044811%5FHEK293%5FK9me3%5FAbcam%5FAlignedSortedChrMrmDuprmBLrmFiltered.bw
GSM6044811%5FHEK2
H3K9me3
GSE175320%5FENCFF000VKT%5Fsignal%5Fhg19.bigWig
GSE175320%5FENCFF
H3K36me3
GSE175195%5FENCFF818NCB%5Ffold%5Fchange%5Fover%5Fcontrol%5FGRCh38.bigWig
GSE175195%5FENCFF
SETDB1

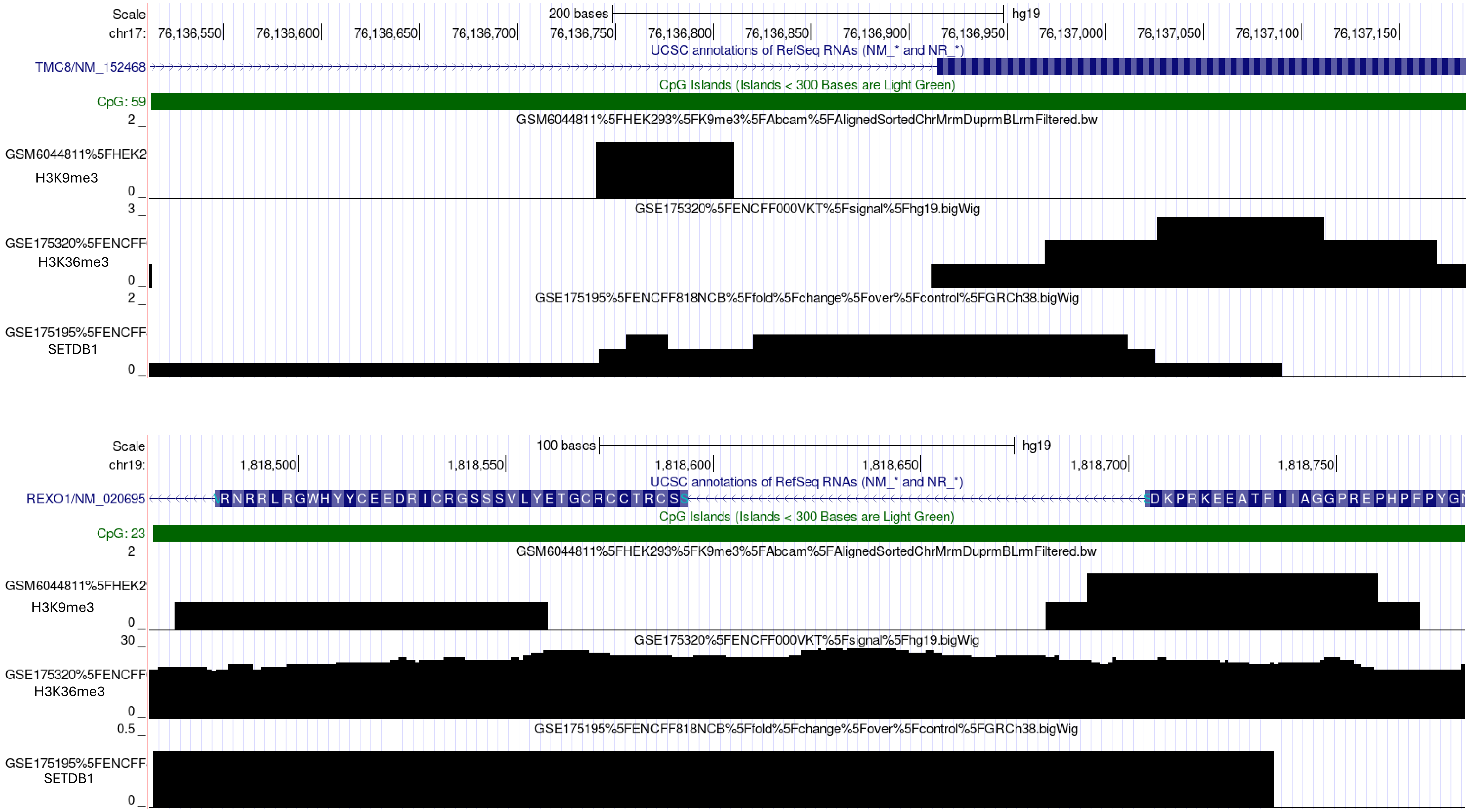
Scale
200 bases
hg19
chr17: 76,136,550 76,136,600 76,136,650 76,136,700 76,136,750 76,136,800 76,136,850 76,136,900 76,136,950 76,137,000 76,137,050 76,137,100 76,137,150
UCSC annotations of RefSeq RNAs (NM_* and NR_*)
TMC8/NM_152468
CpG Islands (Islands < 300 Bases are Light Green)
CpG: 59
GSM6044811%5FHEK293%5FK9me3%5FAbcam%5FAlignedSortedChrMrmDuprmBLrmFiltered.bw
GSM6044811%5FHEK2
H3K9me3
GSE175320%5FENCFF000VKT%5Fsignal%5Fhg19.bigWig
GSE175320%5FENCFF
H3K36me3
GSE175195%5FENCFF818NCB%5Ffold%5Fchange%5Fover%5Fcontrol%5FGRCh38.bigWig
GSE175195%5FENCFF
SETDB1
Scale
100 bases
hg19
chr19: 1,818,500 1,818,550 1,818,600 1,818,650 1,818,700 1,818,750
UCSC annotations of RefSeq RNAs (NM_* and NR_*)
REXO1/NM_020695
RNRRLRGWHYYCEEDRICRGSSSVLYETGCRCCTRCSS
DKPRKEEATFIIAGGPREPHPFPYG
CpG Islands (Islands < 300 Bases are Light Green)
CpG: 23
GSM6044811%5FHEK293%5FK9me3%5FAbcam%5FAlignedSortedChrMrmDuprmBLrmFiltered.bw
GSM6044811%5FHEK2
H3K9me3
GSE175320%5FENCFF000VKT%5Fsignal%5Fhg19.bigWig
GSE175320%5FENCFF
H3K36me3
GSE175195%5FENCFF818NCB%5Ffold%5Fchange%5Fover%5Fcontrol%5FGRCh38.bigWig
GSE175195%5FENCFF
SETDB1

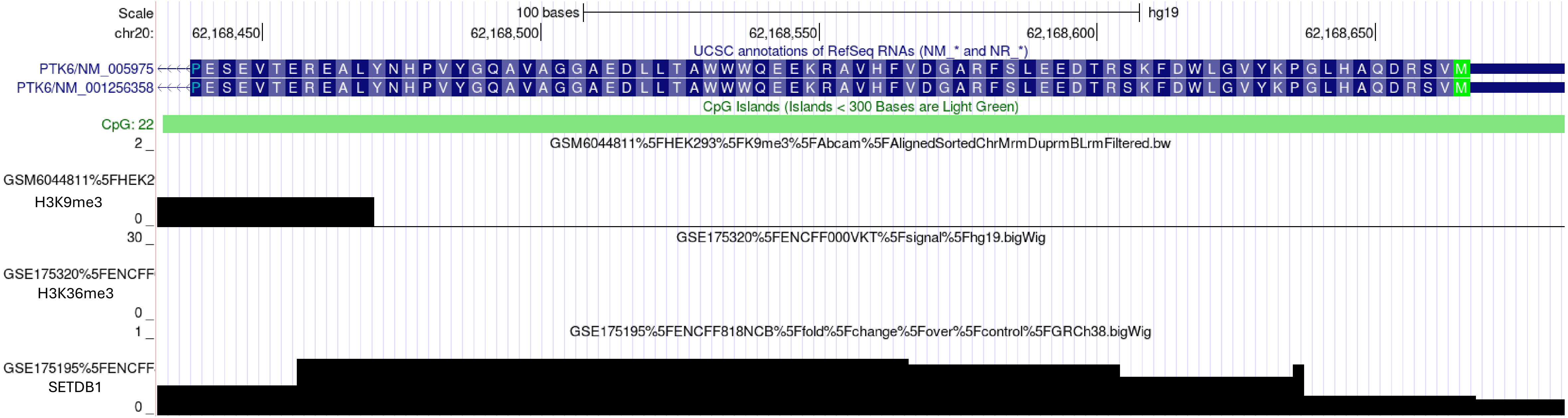

Scale
100 bases
hg19
chr20:
62,168,450
62,168,500
62,168,550
62,168,600
62,168,650
UCSC annotations of RefSeq RNAs (NM_* and NR_*)
PTK6/NM_005975
PTK6/NM_001256358
CpG Islands (Islands < 300 Bases are Light Green)
CpG: 22
GSM6044811%5FHEK293%5FK9me3%5FAbcam%5FAlignedSortedChrMrmDuprmBLrmFiltered.bw
GSM6044811%5FHEK2
H3K9me3
GSE175320%5FENCFF000VKT%5Fsignal%5Fhg19.bigWig
GSE175320%5FENCFF
H3K36me3
GSE175195%5FENCFF818NCB%5Ffold%5Fchange%5Fover%5Fcontrol%5FGRCh38.bigWig
GSE175195%5FENCFF
SETDB1

Suppl. Fig. 6

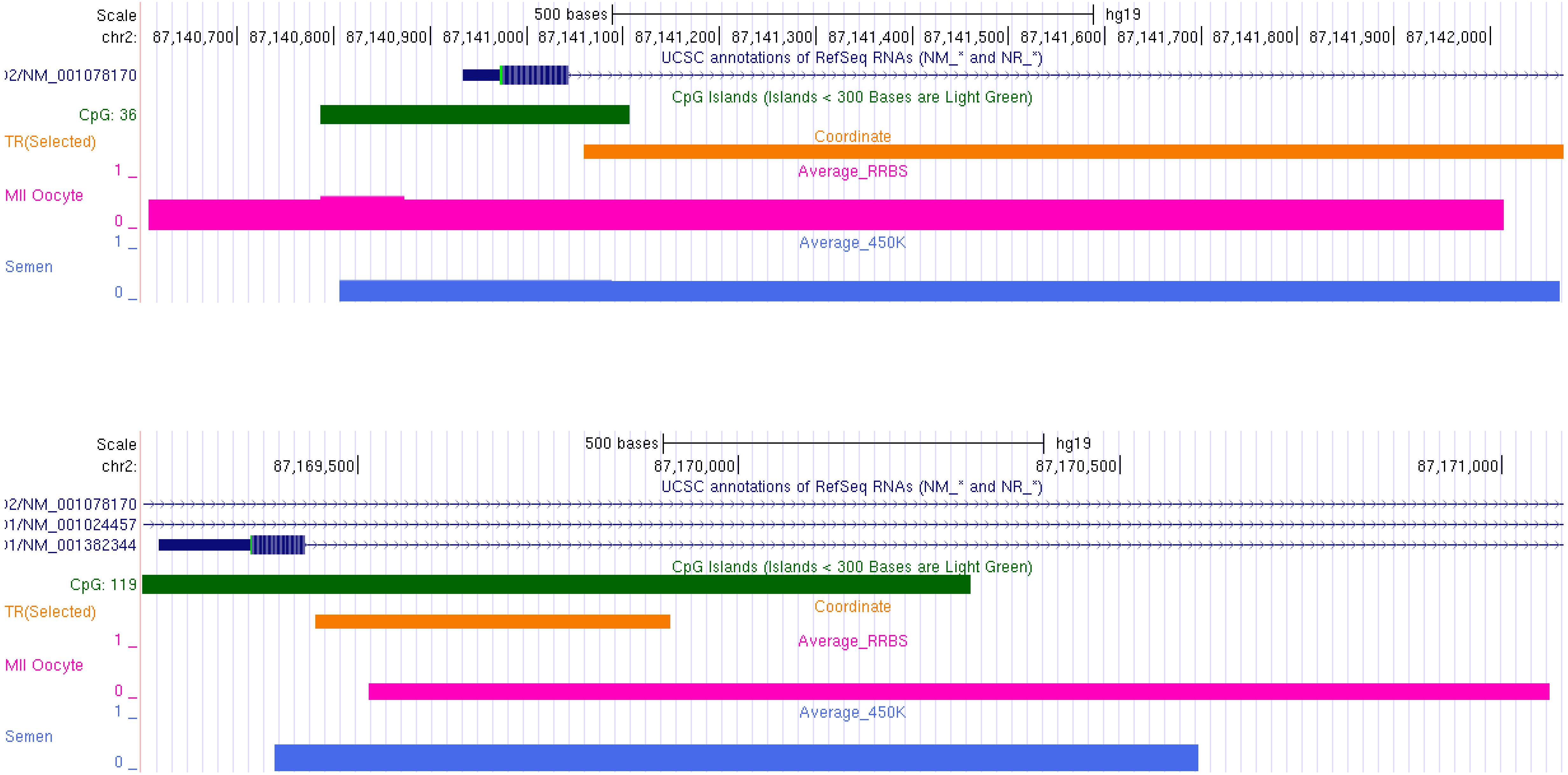
Scale
chr2:
500 bases
hg19
87,140,700
87,140,800
87,140,900
87,141,000
87,141,100
87,141,200
87,141,300
87,141,400
87,141,500
87,141,600
87,141,700
87,141,800
87,141,900
87,142,000
UCSC annotations of RefSeq RNAs (NM_* and NR_*)
)2/NM_001078170
CpG Islands (Islands < 300 Bases are Light Green)
CpG: 36
TR(Selected)
Coordinate
Average_RRBS
1 _
MII Oocyte
0 _
1 _
Average_450K
Semen
0 _
Scale
chr2:
500 bases
hg19
87,169,500
87,170,000
87,170,500
87,171,000
UCSC annotations of RefSeq RNAs (NM_* and NR_*)
)2/NM_001078170
)1/NM_001024457
)1/NM_001382344
CpG Islands (Islands < 300 Bases are Light Green)
CpG: 119
TR(Selected)
Coordinate
Average_RRBS
1 _
MII Oocyte
0 _
1 _
Average_450K
Semen
0 _

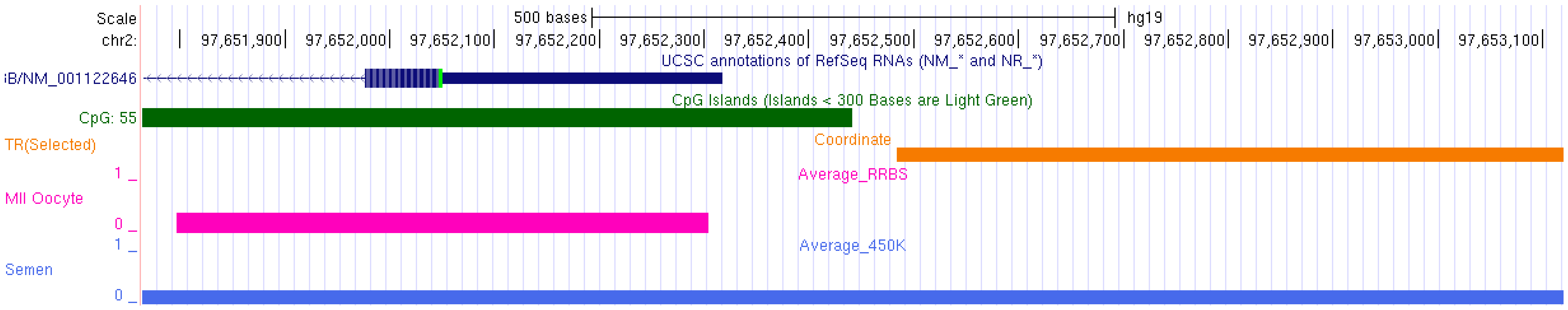
Scale
500 bases
hg19
chr2:
97,651,900
97,652,000
97,652,100
97,652,200
97,652,300
97,652,400
97,652,500
97,652,600
97,652,700
97,652,800
97,652,900
97,653,000
97,653,100
UCSC annotations of RefSeq RNAs (NM_* and NR_*)
B/NM_001122646
CpG Islands (Islands < 300 Bases are Light Green)
CpG: 55
TR(Selected)
Coordinate
1 _
Average_RRBS
MII Oocyte
0 _
1 _
Average_450K
Semen
0 _

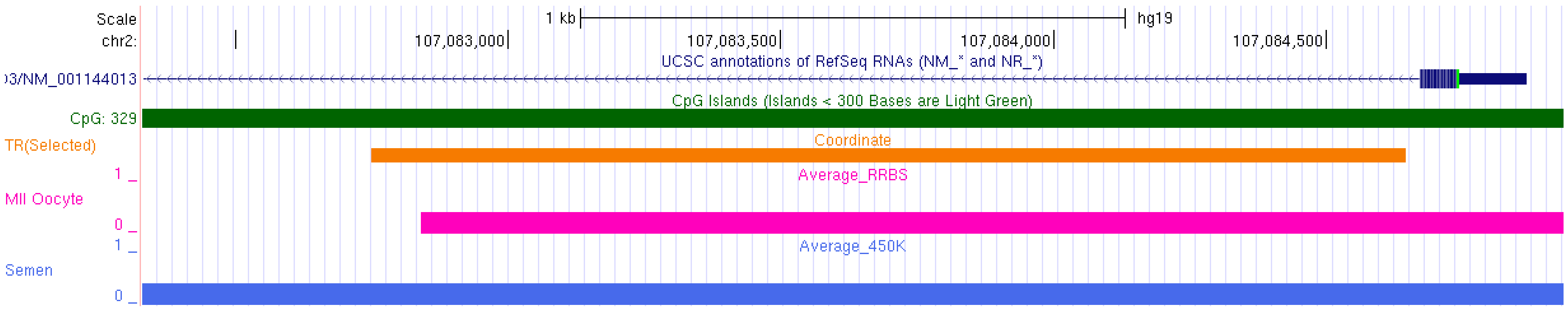
Scale
1 kb
hg19
chr2:
107,083,000
107,083,500
107,084,000
107,084,500
UCSC annotations of RefSeq RNAs (NM_* and NR_*)
3/NM_001144013
CpG Islands (Islands < 300 Bases are Light Green)
CpG: 329
TR(Selected)
Coordinate
1 _
Average_RRBS
MII Oocyte
0 _
1 _
Average_450K
Semen
0 _

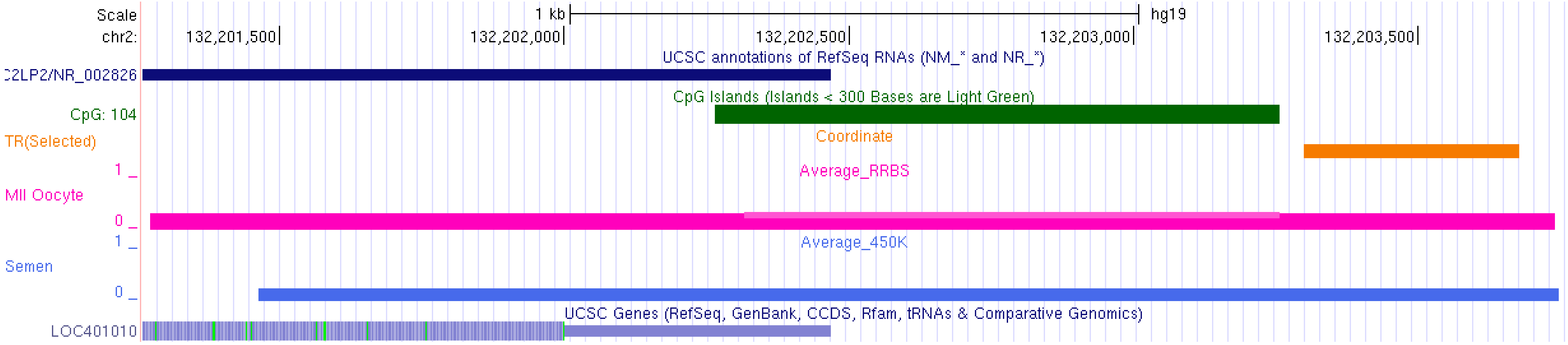
Scale
chr2:
1 kb
hg19
132,201,500
132,202,000
132,202,500
132,203,000
132,203,500
UCSC annotations of RefSeq RNAs (NM_* and NR_*)
C2LP2/NR_002826
CpG Islands (Islands < 300 Bases are Light Green)
CpG: 104
TR(Selected)
Coordinate
Average_RRBS
MII Oocyte
Average_450K
Semen
UCSC Genes (RefSeq, GenBank, CCDS, Rfam, tRNAs & Comparative Genomics)
LOC401010

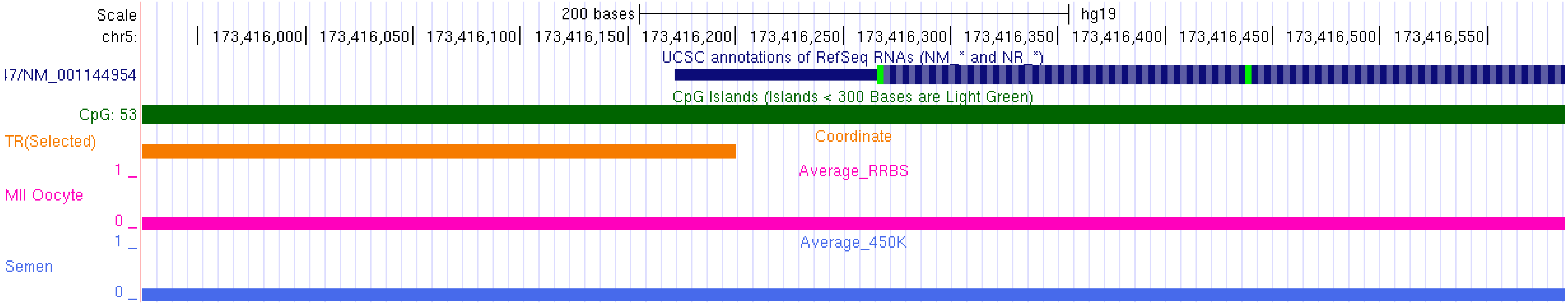
Scale
chr5:
200 bases
hg19
173,416,000
173,416,050
173,416,100
173,416,150
173,416,200
173,416,250
173,416,300
173,416,350
173,416,400
173,416,450
173,416,500
173,416,550
UCSC annotations of RefSeq RNAs (NM_* and NR_*)
I7/NM_001144954
CpG Islands (Islands < 300 Bases are Light Green)
CpG: 53
TR(Selected)
Coordinate
Average_RRBS
MII Oocyte
Average_450K
Semen

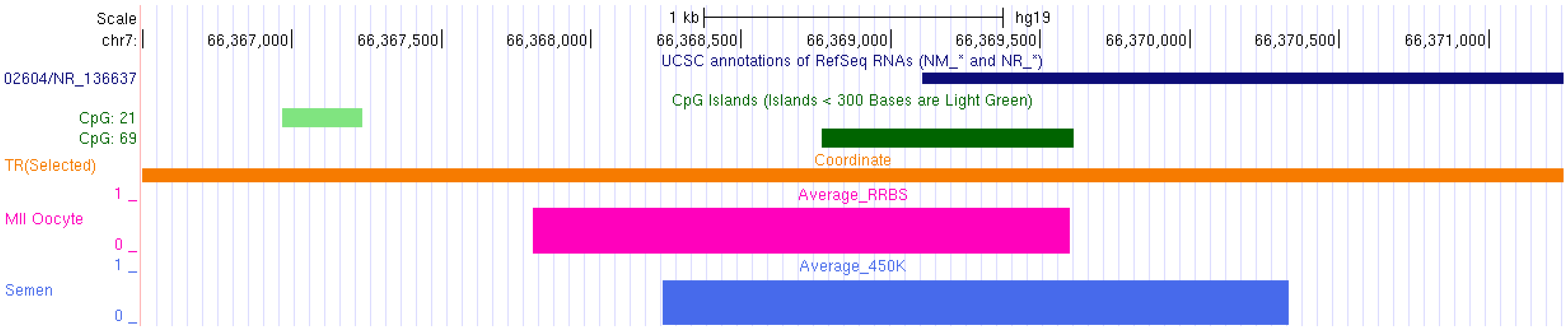
Scale
chr7:
1 kb
hg19
66,367,000
66,367,500
66,368,000
66,368,500
66,369,000
66,369,500
66,370,000
66,370,500
66,371,000
UCSC annotations of RefSeq RNAs (NM_* and NR_*)
02604/NR_136637
CpG Islands (Islands < 300 Bases are Light Green)
CpG: 21
CpG: 69
TR(Selected)
Coordinate
Average_RRBS
MII Oocyte
Average_450K
Semen

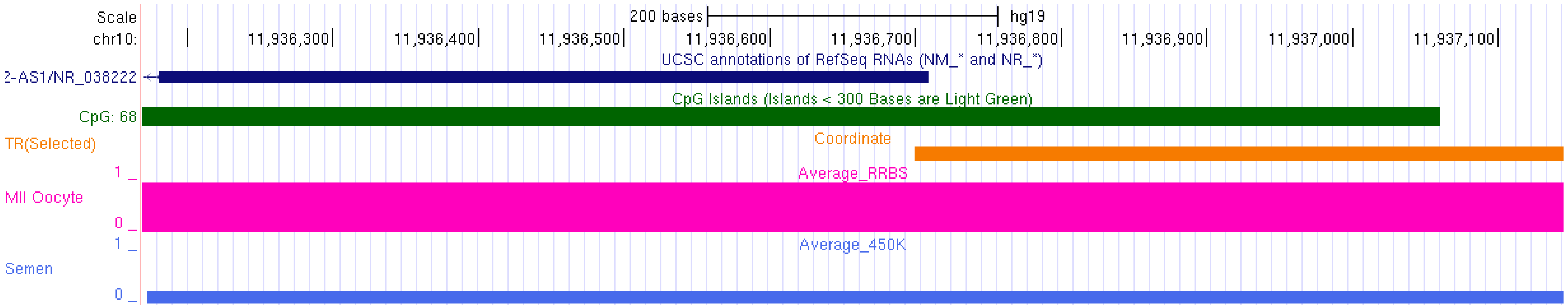
Scale
chr10:
200 bases
hg19
11,936,300
11,936,400
11,936,500
11,936,600
11,936,700
11,936,800
11,936,900
11,937,000
11,937,100
UCSC annotations of RefSeq RNAs (NM_* and NR_*)
2-AS1/NR_038222
CpG Islands (Islands < 300 Bases are Light Green)
CpG: 68
TR(Selected)
Coordinate
Average_RRBS
MII Oocyte
Average_450K
Semen

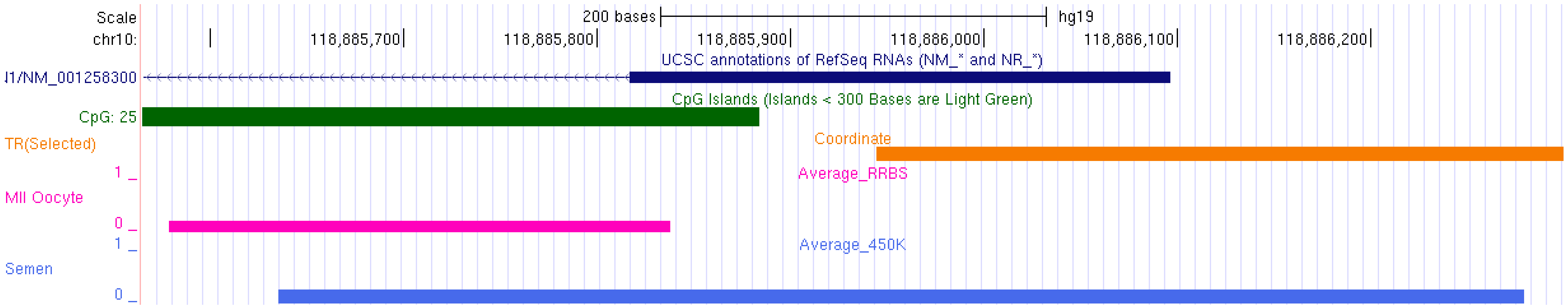
Scale
chr10:
200 bases
hg19
118,885,700
118,885,800
118,885,900
118,886,000
118,886,100
118,886,200
UCSC annotations of RefSeq RNAs (NM_* and NR_*)
l1/NM_001258300
CpG Islands (Islands < 300 Bases are Light Green)
CpG: 25
TR(Selected)
Coordinate
Average_RRBS
MII Oocyte
Average_450K
Semen
1
0
1
0

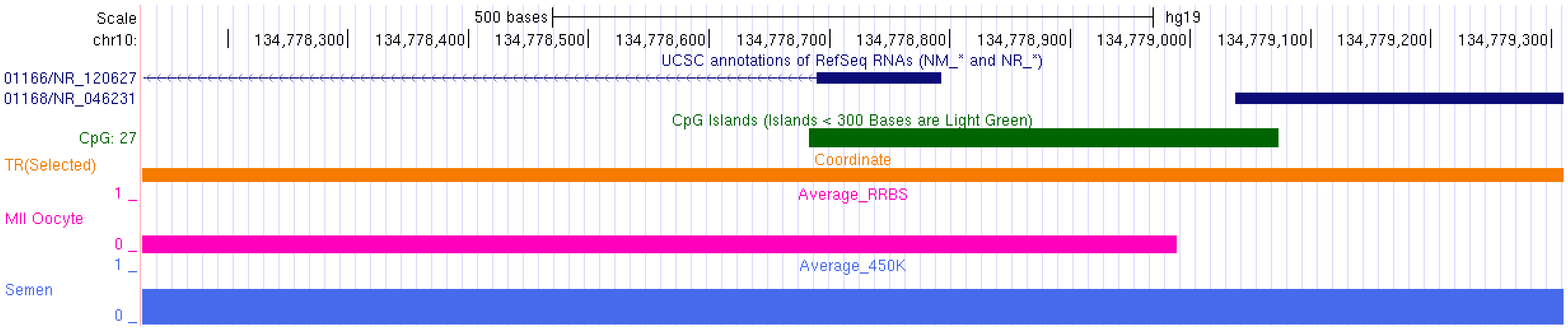
Scale
chr10:
500 bases
hg19
134,778,300
134,778,400
134,778,500
134,778,600
134,778,700
134,778,800
134,778,900
134,779,000
134,779,100
134,779,200
134,779,300
UCSC annotations of RefSeq RNAs (NM_* and NR_*)
01166/NR_120627
01168/NR_046231
CpG Islands (Islands < 300 Bases are Light Green)
CpG: 27
TR(Selected)
Coordinate
Average_RRBS
MII Oocyte
Average_450K
Semen
1
0
1
0

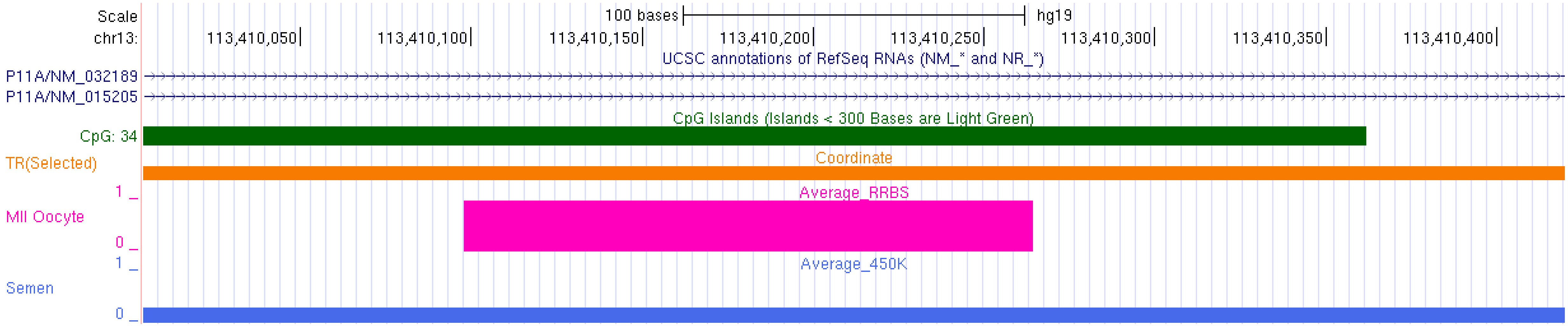
Scale
chr13:
100 bases
hg19
113,410,050
113,410,100
113,410,150
113,410,200
113,410,250
113,410,300
113,410,350
113,410,400
UCSC annotations of RefSeq RNAs (NM_* and NR_*)
P11A/NM_032189
P11A/NM_015205
CpG Islands (Islands < 300 Bases are Light Green)
CpG: 34
TR(Selected)
Coordinate
Average_RRBS
MII Oocyte
Average_450K
Semen

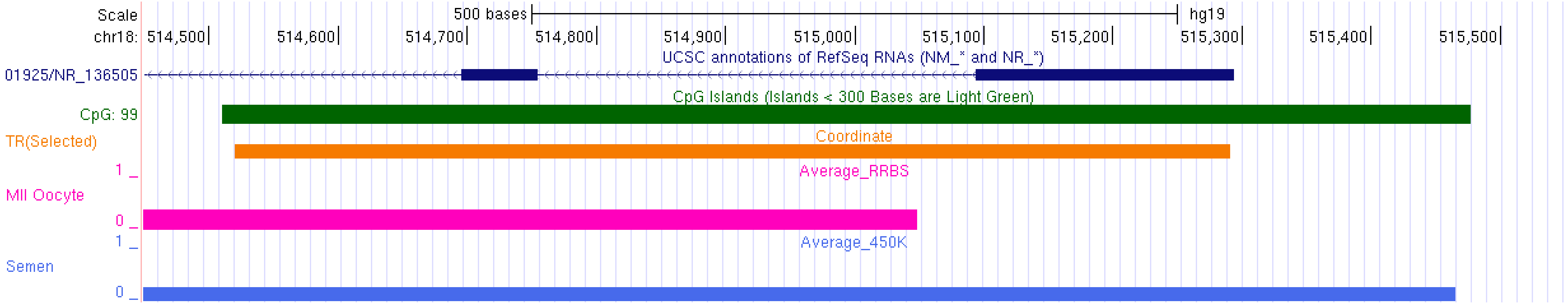
Scale
chr18:
500 bases
hg19
514,500
514,600
514,700
514,800
514,900
515,000
515,100
515,200
515,300
515,400
515,500
UCSC annotations of RefSeq RNAs (NM_* and NR_*)
01925/NR_136505
CpG Islands (Islands < 300 Bases are Light Green)
CpG: 99
TR(Selected)
Coordinate
Average_RRBS
MII Oocyte
Average_450K
Semen

Suppl. Fig. 7

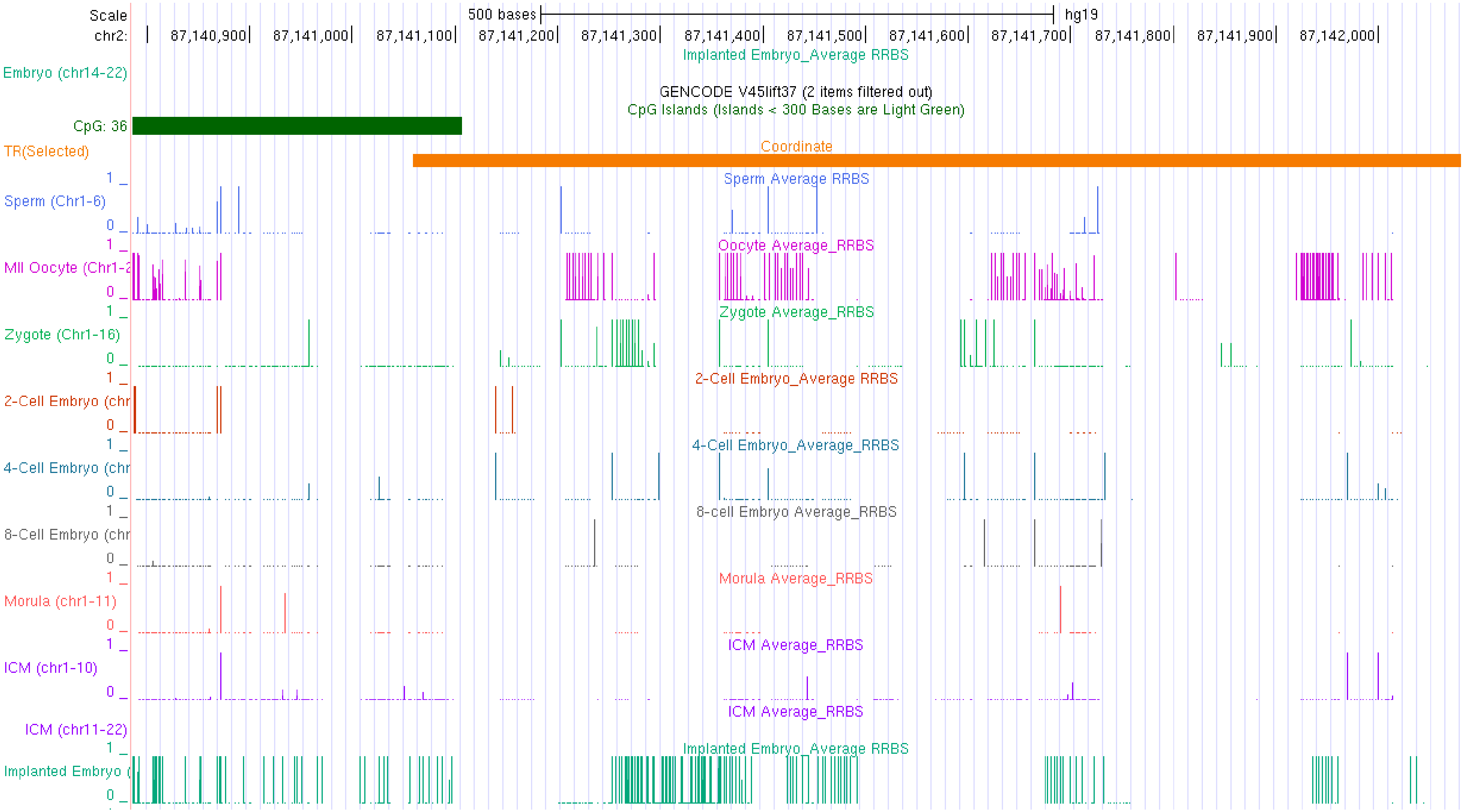
Scale
500 bases
hg19
chr2:
87,140,900
87,141,000
87,141,100
87,141,200
87,141,300
87,141,400
87,141,500
87,141,600
87,141,700
87,141,800
87,141,900
87,142,000
Implanted Embryo_Average RRBS
Embryo (chr14-22)
GENCODE V45lift37 (2 items filtered out)
CpG Islands (Islands < 300 Bases are Light Green)
CpG: 36
TR(Selected)
Coordinate
Sperm Average RRBS
Sperm (Chr1-6)
Oocyte Average_RRBS
MII Oocyte (Chr1-2
Zygote Average_RRBS
Zygote (Chr1-16)
2-Cell Embryo_Average RRBS
2-Cell Embryo (chr
4-Cell Embryo_Average_RRBS
4-Cell Embryo (chr
8-cell Embryo Average_RRBS
8-Cell Embryo (chr
Morula Average_RRBS
Morula (chr1-11)
ICM Average_RRBS
ICM (chr1-10)
ICM Average_RRBS
ICM (chr11-22)
Implanted Embryo_Average RRBS
Implanted Embryo (

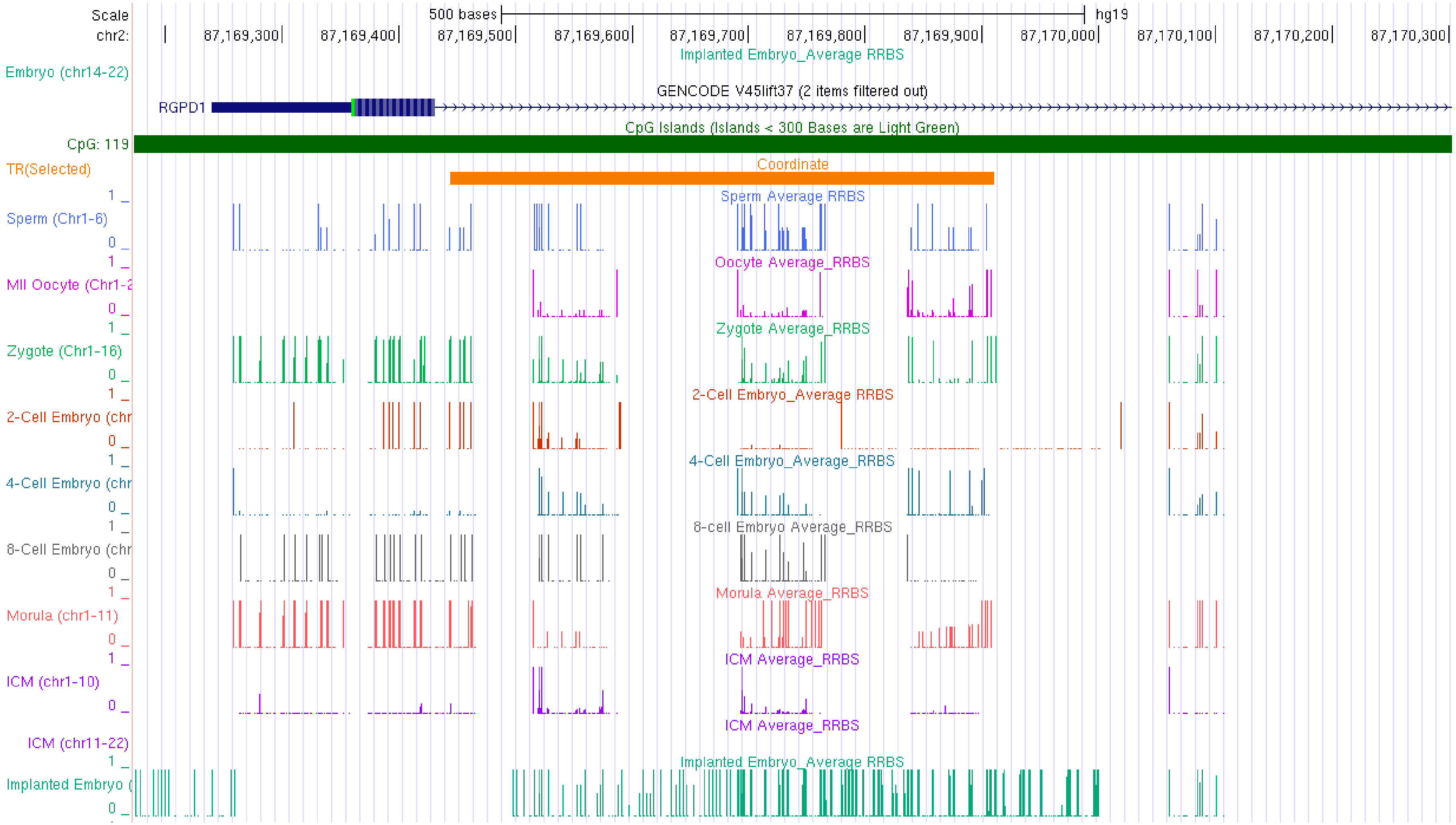

Scale
chr2:
500 bases
hg19
87,169,300
87,169,400
87,169,500
87,169,600
87,169,700
87,169,800
87,169,900
87,170,000
87,170,100
87,170,200
87,170,300
Implanted Embryo_Average RRBS
Embryo (chr14-22)
GENCODE V45lift37 (2 items filtered out)
RGPD1
CpG Islands (Islands < 300 Bases are Light Green)
CpG: 119
TR(Selected)
Coordinate
Sperm Average RRBS
Sperm (Chr1-6)
Oocyte Average_RRBS
MII Oocyte (Chr1-2
Zygote Average_RRBS
Zygote (Chr1-16)
2-Cell Embryo_Average RRBS
2-Cell Embryo (chr
4-Cell Embryo_Average_RRBS
4-Cell Embryo (chr
8-cell Embryo Average_RRBS
8-Cell Embryo (chr
Morula Average_RRBS
Morula (chr1-11)
ICM Average_RRBS
ICM (chr1-10)
ICM Average_RRBS
ICM (chr11-22)
Implanted Embryo_Average RRBS
Implanted Embryo (

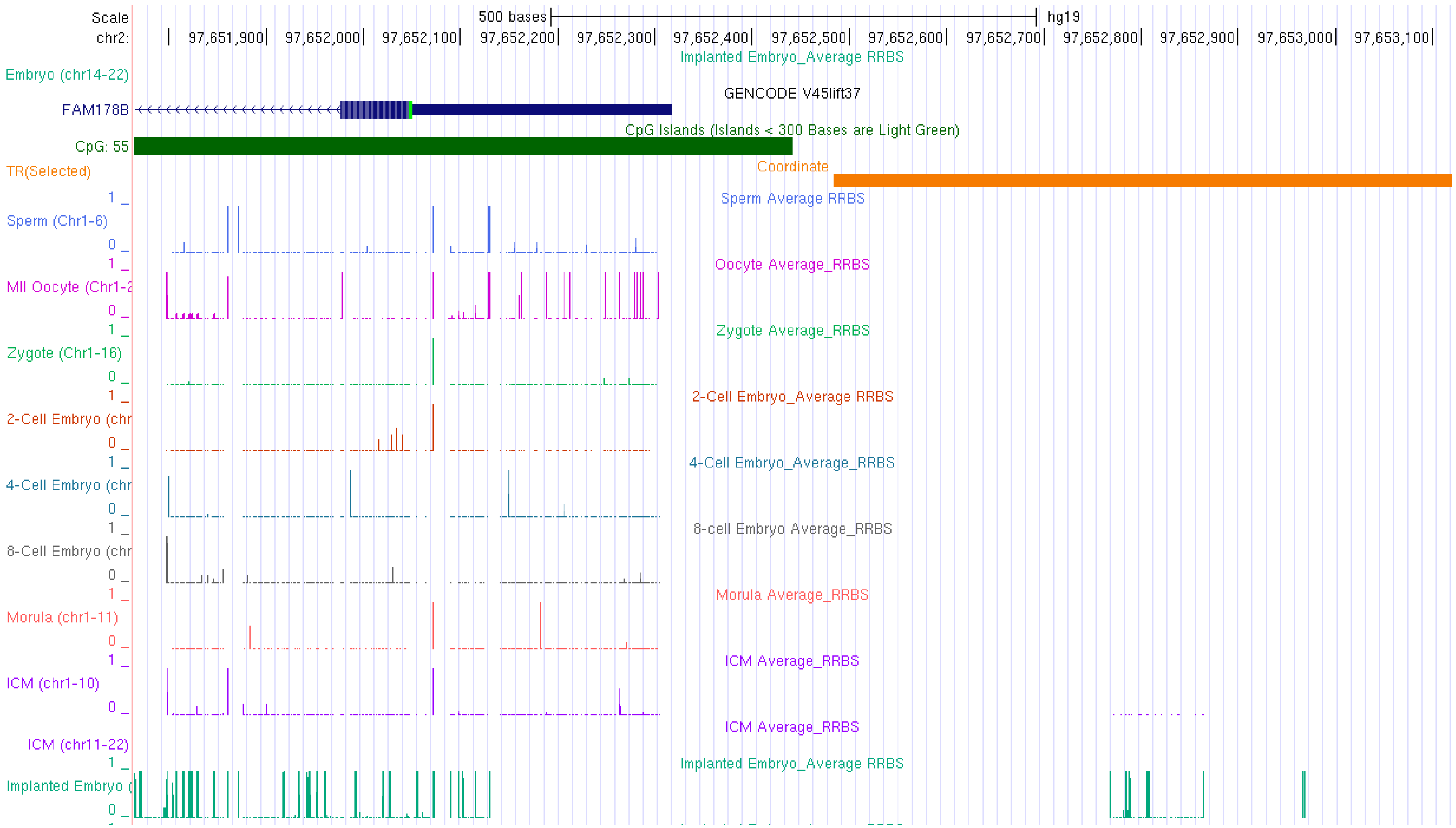
Scale
chr2:
500 bases
hg19
97,651,900
97,652,000
97,652,100
97,652,200
97,652,300
97,652,400
97,652,500
97,652,600
97,652,700
97,652,800
97,652,900
97,653,000
97,653,100
Implanted Embryo_Average RRBS
Embryo (chr14-22)
GENCODE V45lift37
FAM178B
CpG Islands (Islands < 300 Bases are Light Green)
CpG: 55
Coordinate
TR(Selected)
Sperm Average RRBS
Sperm (Chr1-6)
Oocyte Average_RRBS
MII Oocyte (Chr1-2
Zygote Average_RRBS
Zygote (Chr1-16)
2-Cell Embryo_Average RRBS
2-Cell Embryo (chr
4-Cell Embryo_Average_RRBS
4-Cell Embryo (chr
8-cell Embryo Average_RRBS
8-Cell Embryo (chr
Morula Average_RRBS
Morula (chr1-11)
ICM Average_RRBS
ICM (chr1-10)
ICM Average_RRBS
ICM (chr11-22)
Implanted Embryo_Average RRBS
Implanted Embryo (

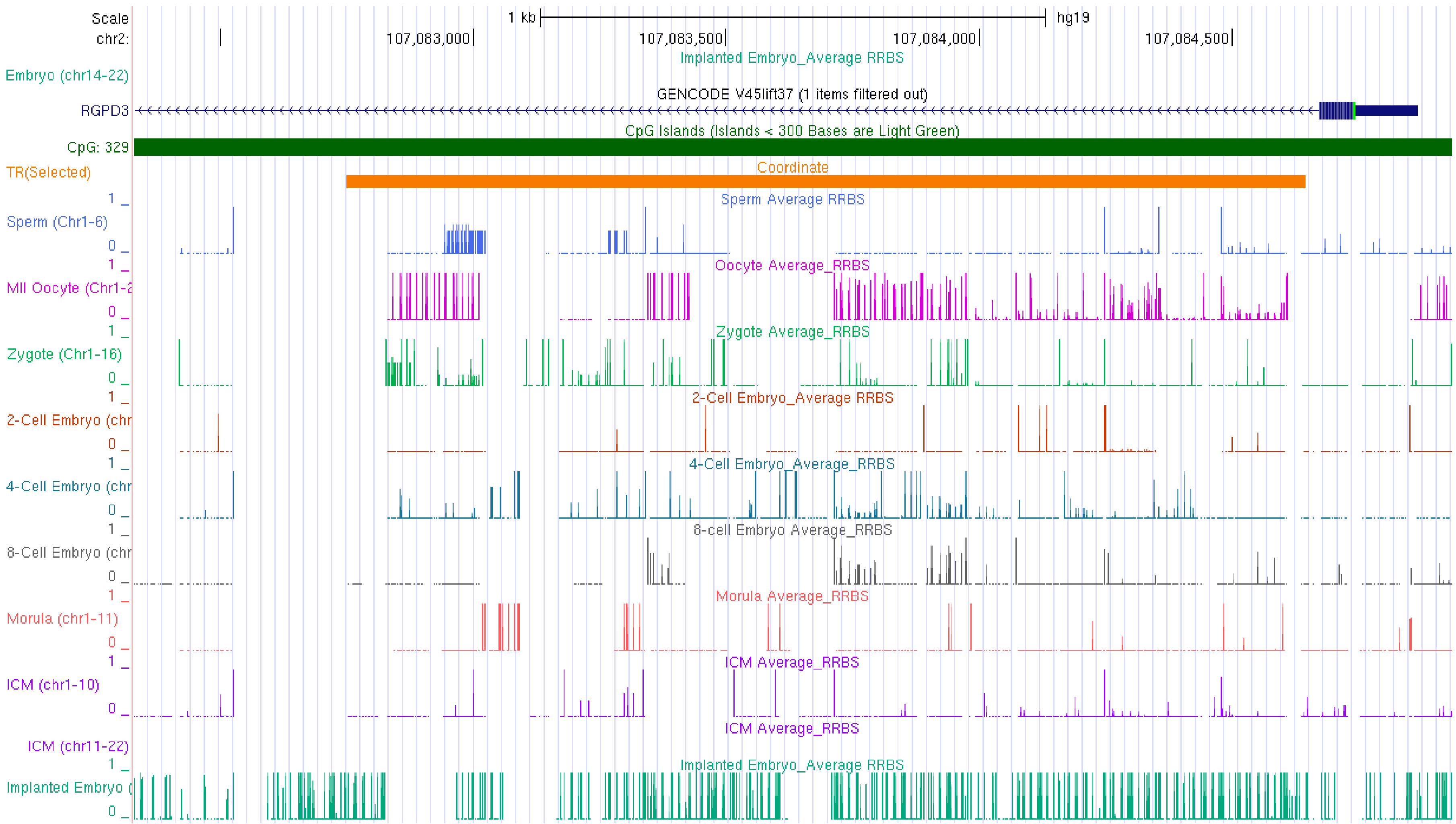

Scale
chr2:
1 kb
hg19
107,083,000
107,083,500
107,084,000
107,084,500
Implanted Embryo_Average RRBS
Embryo (chr14-22)
GENCODE V45lift37 (1 items filtered out)
RGPD3
CpG Islands (Islands < 300 Bases are Light Green)
CpG: 329
TR(Selected)
Coordinate
Sperm Average RRBS
Sperm (Chr1-6)
Oocyte Average_RRBS
MII Oocyte (Chr1-2
Zygote Average_RRBS
Zygote (Chr1-16)
2-Cell Embryo_Average RRBS
2-Cell Embryo (chr
4-Cell Embryo_Average_RRBS
4-Cell Embryo (chr
8-cell Embryo Average_RRBS
8-Cell Embryo (chr
Morula Average_RRBS
Morula (chr1-11)
ICM Average_RRBS
ICM (chr1-10)
ICM Average_RRBS
ICM (chr11-22)
Implanted Embryo_Average RRBS
Implanted Embryo (

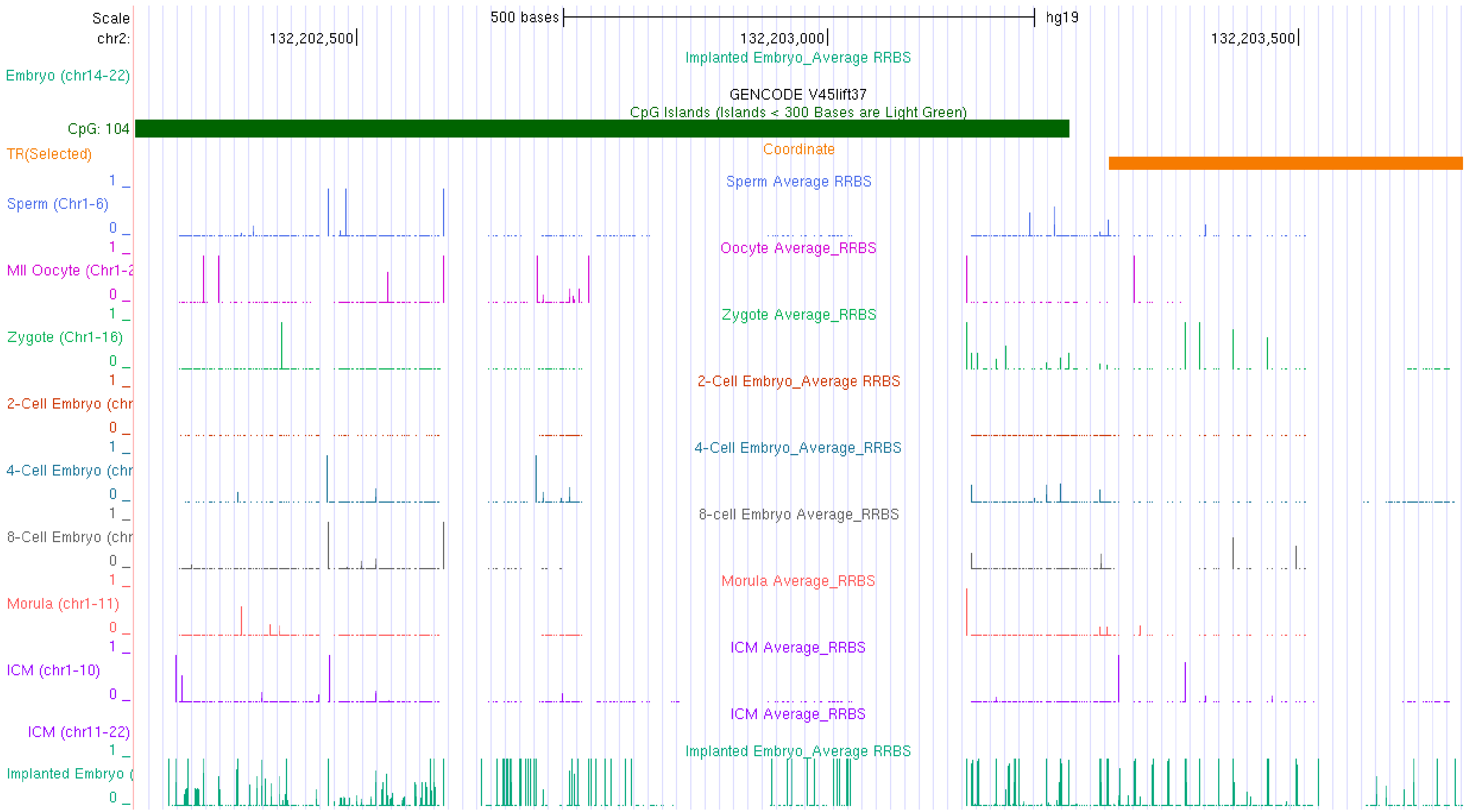
Scale
chr2:
500 bases
hg19
132,202,500
132,203,000
132,203,500
Implanted Embryo_Average RRBS
Embryo (chr14-22)
GENCODE V45lift37
CpG Islands (Islands < 300 Bases are Light Green)
CpG: 104
TR(Selected)
Coordinate
Sperm Average RRBS
Sperm (Chr1-6)
Oocyte Average_RRBS
MII Oocyte (Chr1-2
Zygote Average_RRBS
Zygote (Chr1-16)
2-Cell Embryo_Average RRBS
2-Cell Embryo (chr
4-Cell Embryo_Average_RRBS
4-Cell Embryo (chr
8-cell Embryo Average_RRBS
8-Cell Embryo (chr
Morula Average_RRBS
Morula (chr1-11)
ICM Average_RRBS
ICM (chr1-10)
ICM Average_RRBS
ICM (chr11-22)
Implanted Embryo_Average RRBS
Implanted Embryo (

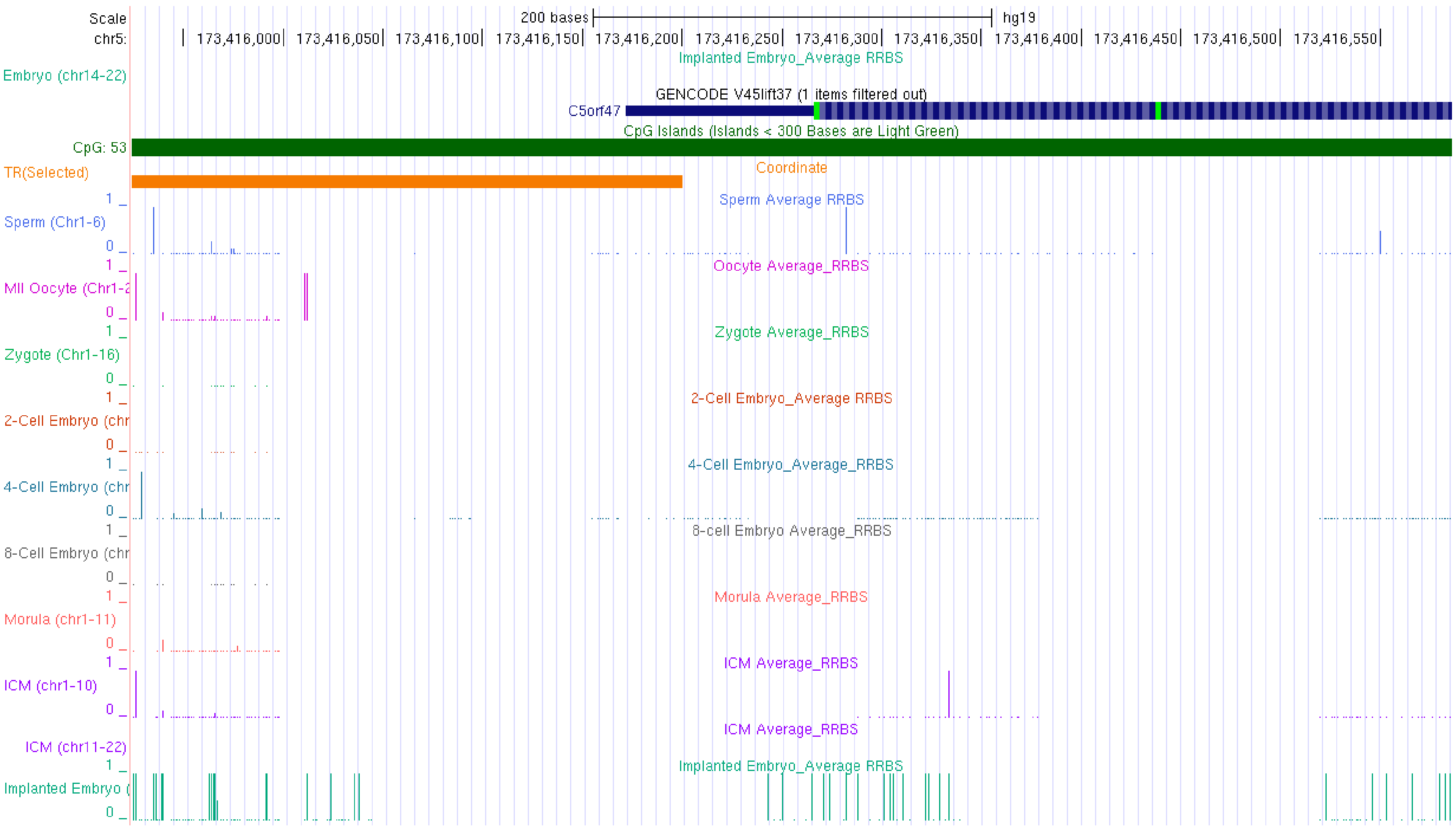

Scale
chr5:
200 bases
hg19
173,416,000
173,416,050
173,416,100
173,416,150
173,416,200
173,416,250
173,416,300
173,416,350
173,416,400
173,416,450
173,416,500
173,416,550
Implanted Embryo_Average RRBS
Embryo (chr14-22)
GENCODE V45lift37 (1 items filtered out)
C5orf47
CpG Islands (Islands < 300 Bases are Light Green)
CpG: 53
TR(Selected)
Coordinate
Sperm Average RRBS
Sperm (Chr1-6)
Oocyte Average_RRBS
MII Oocyte (Chr1-2
Zygote Average_RRBS
Zygote (Chr1-16)
2-Cell Embryo_Average RRBS
2-Cell Embryo (chr
4-Cell Embryo_Average_RRBS
4-Cell Embryo (chr
8-cell Embryo Average_RRBS
8-Cell Embryo (chr
Morula Average_RRBS
Morula (chr1-11)
ICM Average_RRBS
ICM (chr1-10)
ICM Average_RRBS
ICM (chr11-22)
Implanted Embryo_Average RRBS
Implanted Embryo (

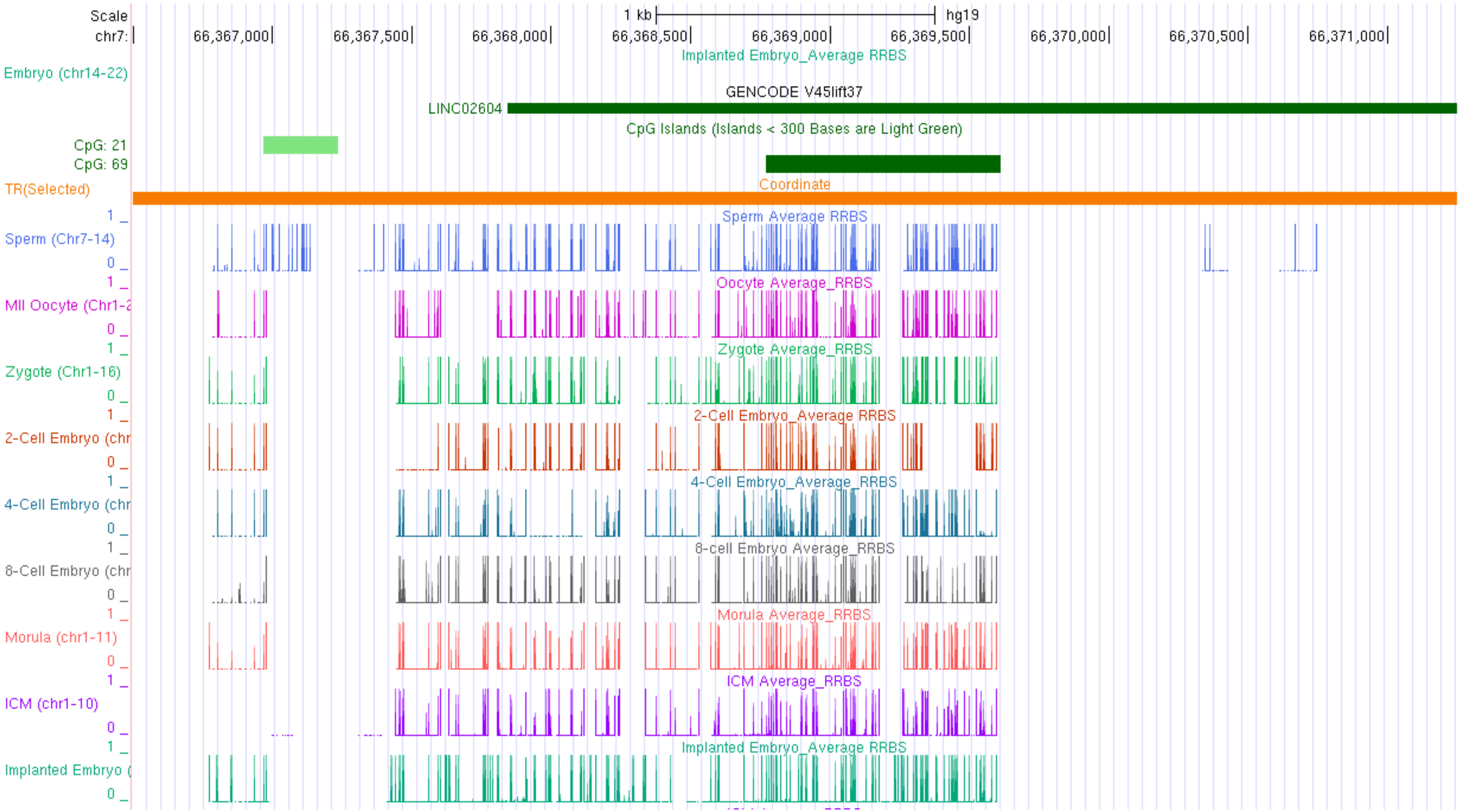
Scale
chr7:
1 kb
hg19
66,367,000
66,367,500
66,368,000
66,368,500
66,369,000
66,369,500
66,370,000
66,370,500
66,371,000
Implanted Embryo_Average RRBS
Embryo (chr14-22)
GENCODE V45lift37
LINC02604
CpG Islands (Islands < 300 Bases are Light Green)
CpG: 21
CpG: 69
TR(Selected)
Coordinate
Sperm Average RRBS
Sperm (Chr7-14)
Oocyte Average_RRBS
MII Oocyte (Chr1-2
Zygote Average_RRBS
Zygote (Chr1-16)
2-Cell Embryo_Average RRBS
2-Cell Embryo (chr
4-Cell Embryo_Average_RRBS
4-Cell Embryo (chr
8-cell Embryo Average_RRBS
8-Cell Embryo (chr
Morula Average_RRBS
Morula (chr1-11)
ICM Average_RRBS
ICM (chr1-10)
Implanted Embryo_Average RRBS
Implanted Embryo (

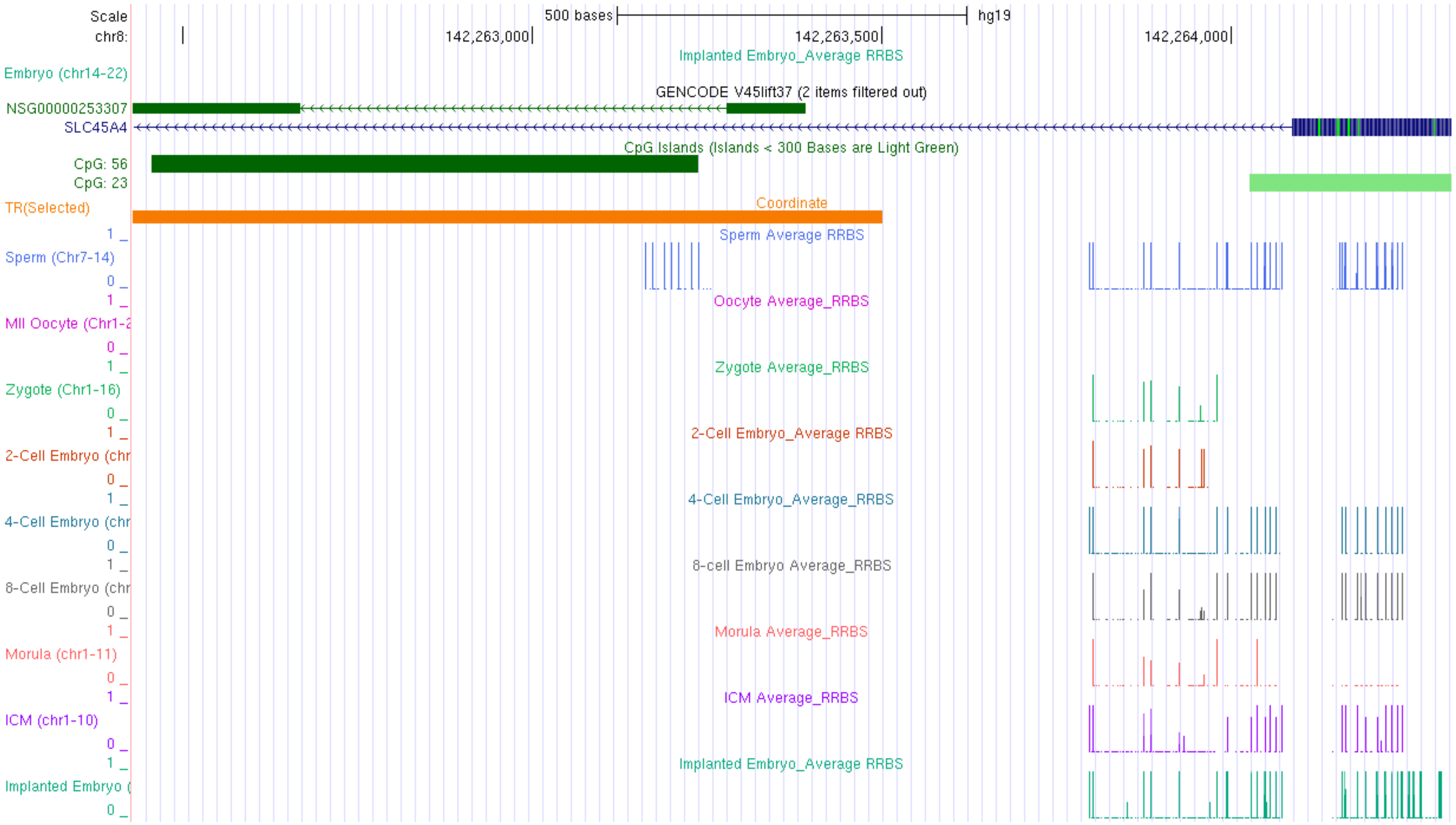
Scale
chr8:
500 bases
hg19
142,263,000
142,263,500
142,264,000
Implanted Embryo_Average RRBS
Embryo (chr14-22)
GENCODE V45lift37 (2 items filtered out)
NSG00000253307
SLC45A4
CpG Islands (Islands < 300 Bases are Light Green)
CpG: 56
CpG: 23
TR(Selected)
Coordinate
Sperm Average RRBS
Sperm (Chr7-14)
Oocyte Average_RRBS
MII Oocyte (Chr1-2
Zygote Average_RRBS
Zygote (Chr1-16)
2-Cell Embryo_Average RRBS
2-Cell Embryo (chr
4-Cell Embryo_Average_RRBS
4-Cell Embryo (chr
8-cell Embryo Average_RRBS
8-Cell Embryo (chr
Morula Average_RRBS
Morula (chr1-11)
ICM Average_RRBS
ICM (chr1-10)
Implanted Embryo_Average RRBS
Implanted Embryo (

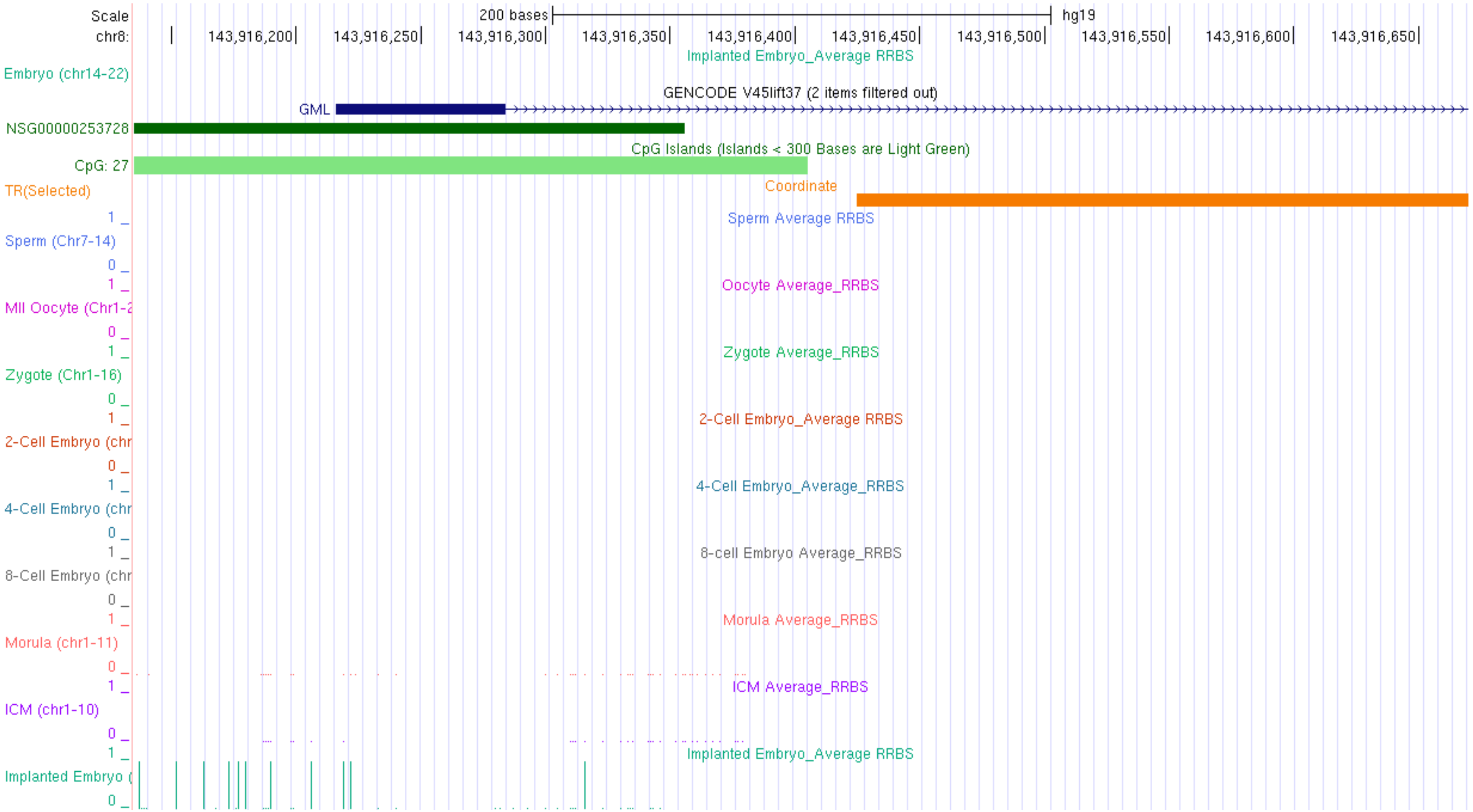
Scale
chr8:
200 bases
hg19
143,916,200
143,916,250
143,916,300
143,916,350
143,916,400
143,916,450
143,916,500
143,916,550
143,916,600
143,916,650
Implanted Embryo_Average RRBS
Embryo (chr14-22)
GENCODE V45lift37 (2 items filtered out)
GML
NSG00000253728
CpG Islands (Islands < 300 Bases are Light Green)
CpG: 27
TR(Selected)
Coordinate
Sperm Average RRBS
Sperm (Chr7-14)
Oocyte Average_RRBS
MII Oocyte (Chr1-2
Zygote Average_RRBS
Zygote (Chr1-16)
2-Cell Embryo_Average RRBS
2-Cell Embryo (chr
4-Cell Embryo_Average_RRBS
4-Cell Embryo (chr
8-cell Embryo Average_RRBS
8-Cell Embryo (chr
Morula Average_RRBS
Morula (chr1-11)
ICM Average_RRBS
ICM (chr1-10)
Implanted Embryo_Average RRBS
Implanted Embryo (

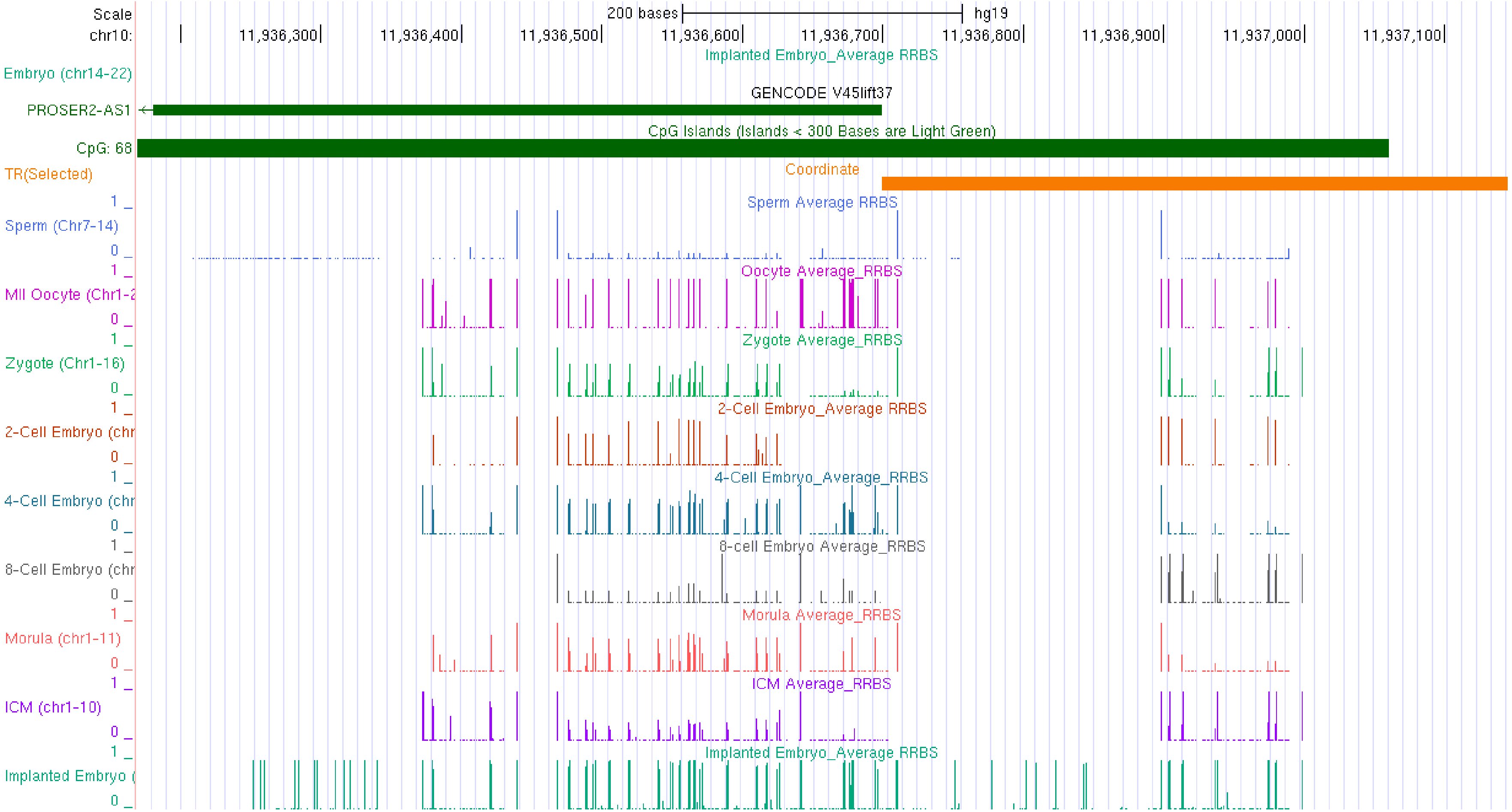

Scale
chr10:
200 bases
hg19
11,936,300
11,936,400
11,936,500
11,936,600
11,936,700
11,936,800
11,936,900
11,937,000
11,937,100
Implanted Embryo_Average RRBS
Embryo (chr14-22)
GENCODE V45lift37
PROSER2-AS1
CpG Islands (Islands < 300 Bases are Light Green)
CpG: 68
Coordinate
TR(Selected)
Sperm Average RRBS
Sperm (Chr7-14)
Oocyte Average_RRBS
MII Oocyte (Chr1-2
Zygote Average_RRBS
Zygote (Chr1-16)
2-Cell Embryo_Average RRBS
2-Cell Embryo (chr
4-Cell Embryo_Average_RRBS
4-Cell Embryo (chr
8-cell Embryo Average_RRBS
8-Cell Embryo (chr
Morula Average_RRBS
Morula (chr1-11)
ICM Average_RRBS
ICM (chr1-10)
Implanted Embryo_Average RRBS
Implanted Embryo (

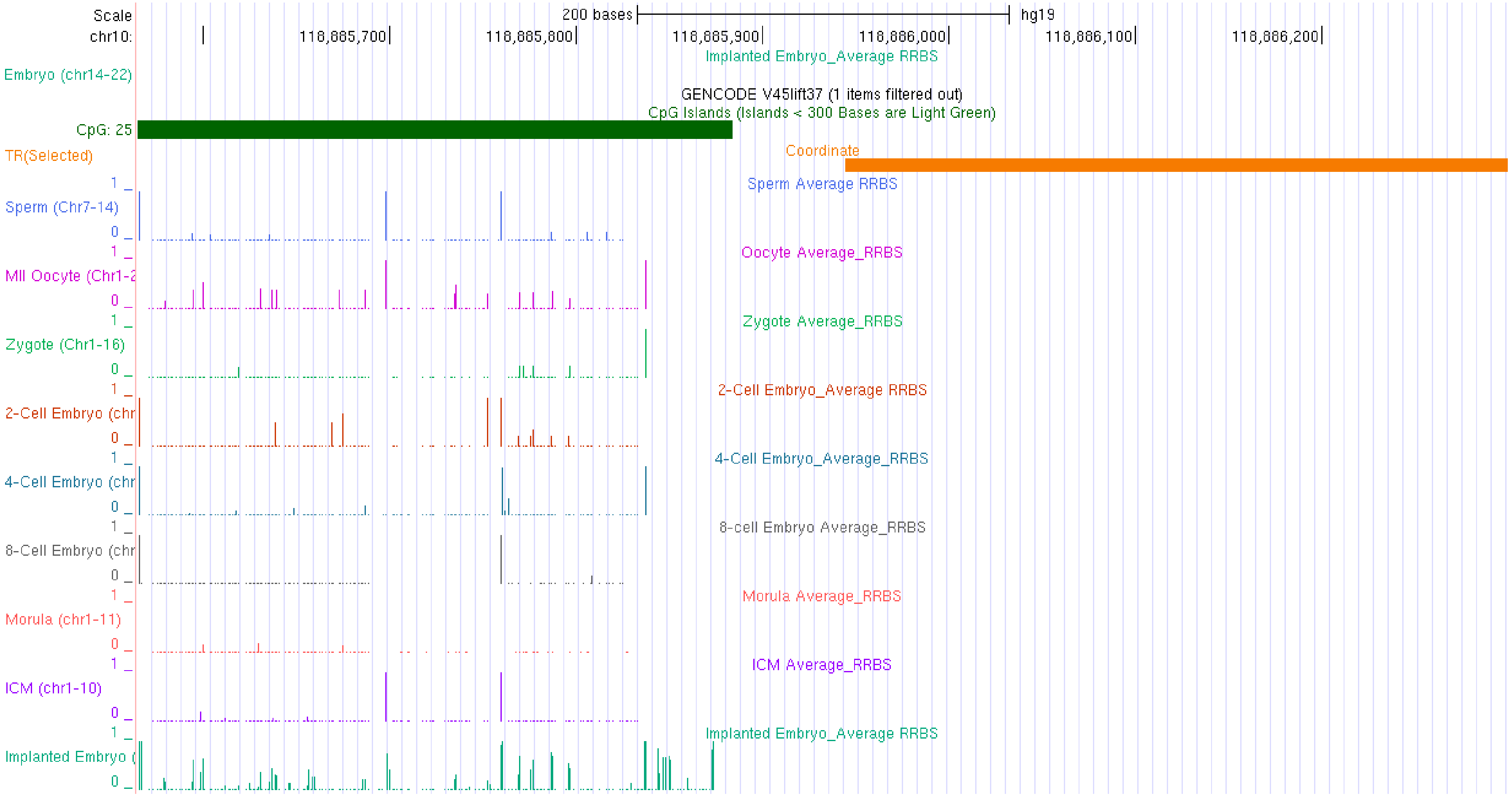
Scale
chr10:
200 bases
hg19
118,885,700
118,885,800
118,885,900
118,886,000
118,886,100
118,886,200
Implanted Embryo_Average RRBS
Embryo (chr14-22)
GENCODE V45lift37 (1 items filtered out)
CpG Islands (Islands < 300 Bases are Light Green)
CpG: 25
Coordinate
TR(Selected)
Sperm Average RRBS
Sperm (Chr7-14)
Oocyte Average_RRBS
MII Oocyte (Chr1-2
Zygote Average_RRBS
Zygote (Chr1-16)
2-Cell Embryo_Average RRBS
2-Cell Embryo (chr
4-Cell Embryo_Average_RRBS
4-Cell Embryo (chr
8-cell Embryo Average_RRBS
8-Cell Embryo (chr
Morula Average_RRBS
Morula (chr1-11)
ICM Average_RRBS
ICM (chr1-10)
Implanted Embryo_Average RRBS
Implanted Embryo (

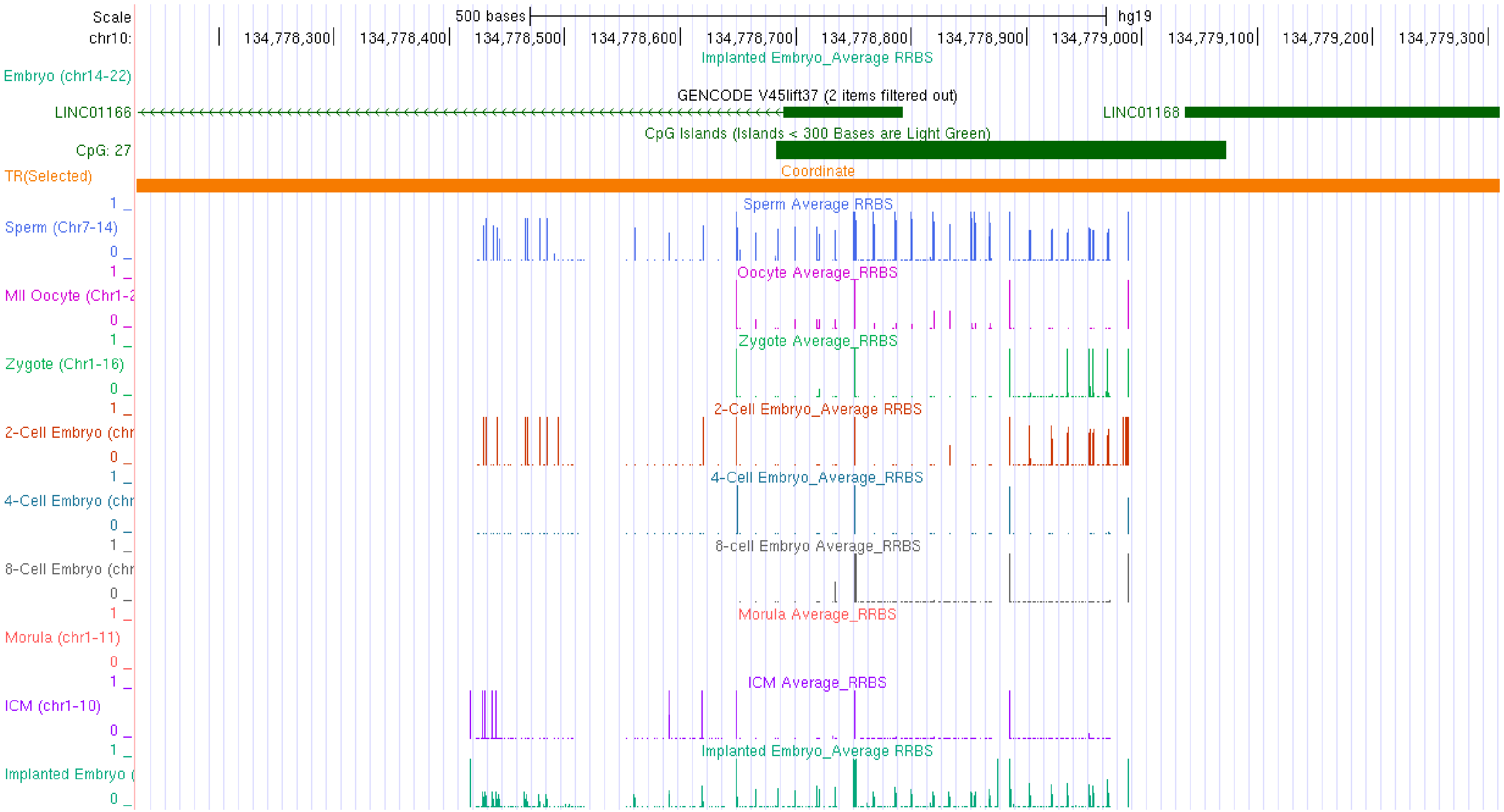

Scale
chr10:
500 bases
hg19
134,778,300
134,778,400
134,778,500
134,778,600
134,778,700
134,778,800
134,778,900
134,779,000
134,779,100
134,779,200
134,779,300
Implanted Embryo_Average RRBS
Embryo (chr14-22)
GENCODE V45lift37 (2 items filtered out)
LINC01166
LINC01168
CpG Islands (Islands < 300 Bases are Light Green)
CpG: 27
Coordinate
TR(Selected)
Sperm Average RRBS
Sperm (Chr7-14)
Oocyte Average_RRBS
MII Oocyte (Chr1-2
Zygote Average_RRBS
Zygote (Chr1-16)
2-Cell Embryo_Average RRBS
2-Cell Embryo (chr
4-Cell Embryo_Average_RRBS
4-Cell Embryo (chr
8-cell Embryo Average_RRBS
8-Cell Embryo (chr
Morula Average_RRBS
Morula (chr1-11)
ICM Average_RRBS
ICM (chr1-10)
Implanted Embryo_Average RRBS
Implanted Embryo (

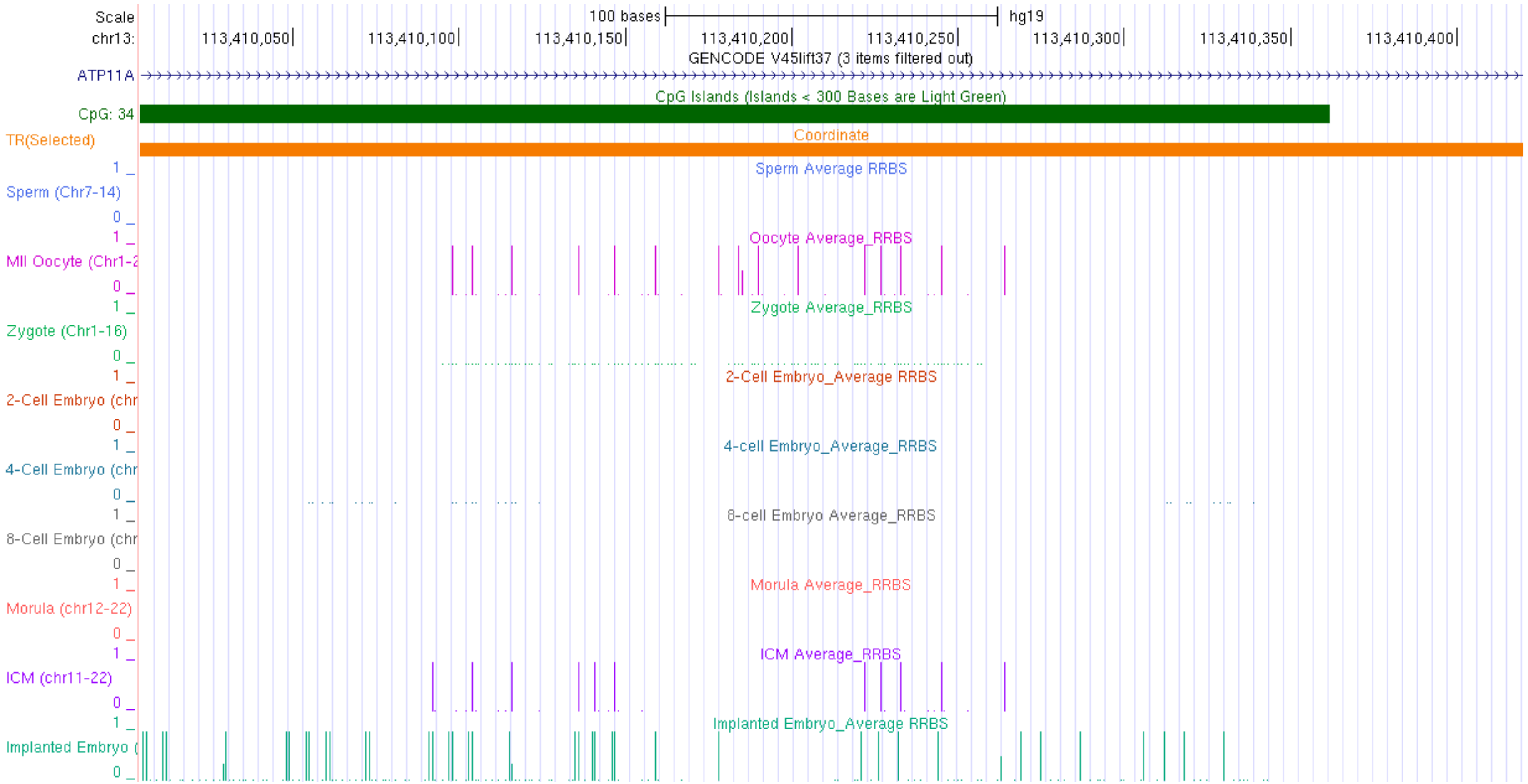

Scale
100 bases
hg19
chr13:
113,410,050
113,410,100
113,410,150
113,410,200
113,410,250
113,410,300
113,410,350
113,410,400
GENCODE V45lift37 (3 items filtered out)
ATP11A
CpG Islands (Islands < 300 Bases are Light Green)
CpG: 34
TR(Selected)
Coordinate
Sperm Average RRBS
Sperm (Chr7-14)
Oocyte Average_RRBS
MII Oocyte (Chr1-2
Zygote Average_RRBS
Zygote (Chr1-16)
2-Cell Embryo_Average RRBS
2-Cell Embryo (chr
4-cell Embryo_Average_RRBS
4-Cell Embryo (chr
8-cell Embryo Average_RRBS
8-Cell Embryo (chr
Morula Average_RRBS
Morula (chr12-22)
ICM Average_RRBS
ICM (chr11-22)
Implanted Embryo_Average RRBS
Implanted Embryo (

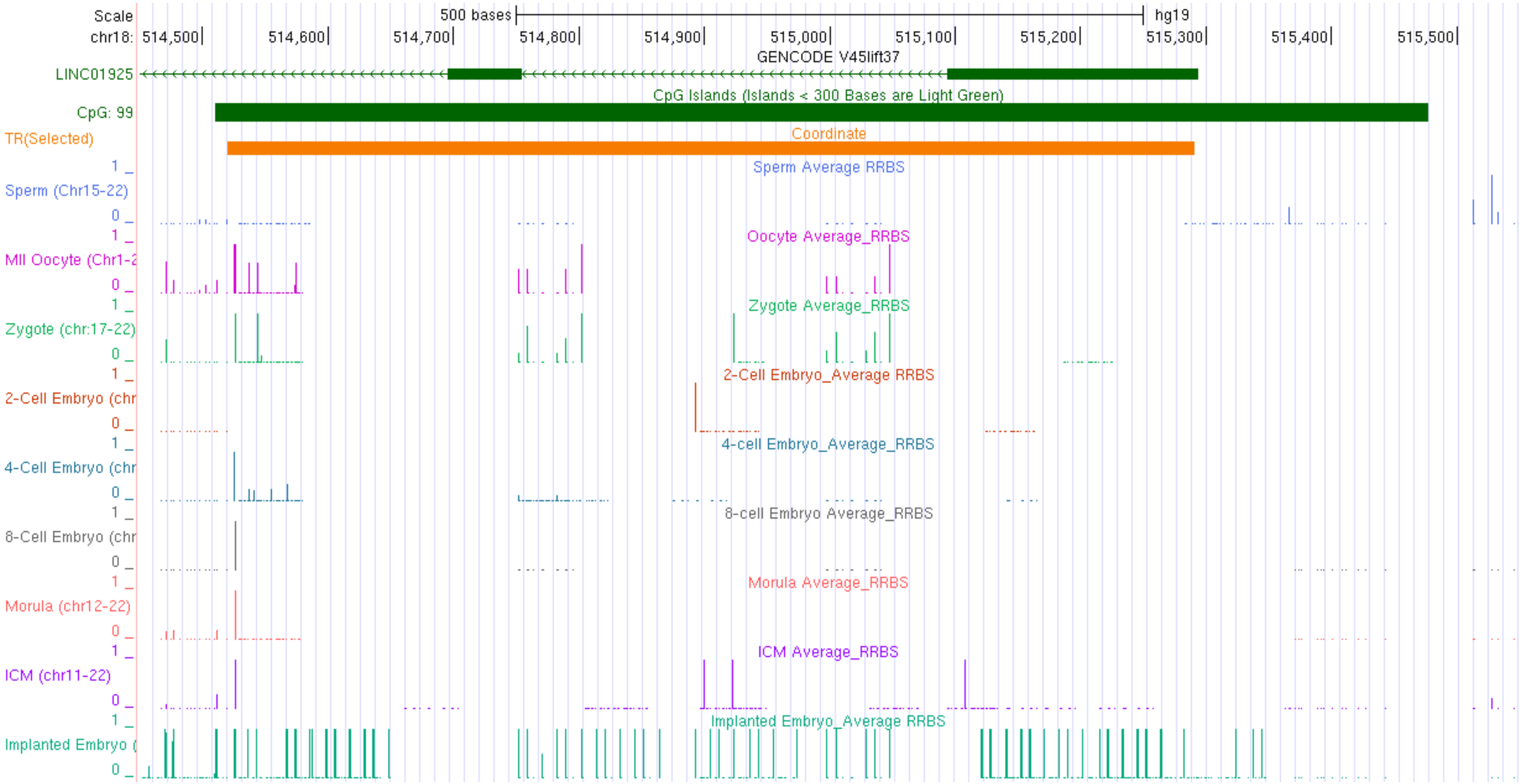

Scale
500 bases
hg19
chr18: 514,500 514,600 514,700 514,800 514,900 515,000 515,100 515,200 515,300 515,400 515,500
GENCODE V45lift37
LINC01925
CpG Islands (Islands < 300 Bases are Light Green)
CpG: 99
TR(Selected)
Coordinate
Sperm Average RRBS
Sperm (Chr15-22)
Oocyte Average_RRBS
MII Oocyte (Chr1-2
Zygote Average_RRBS
Zygote (chr:17-22)
2-Cell Embryo_Average RRBS
2-Cell Embryo (chr
4-cell Embryo_Average_RRBS
4-Cell Embryo (chr
8-cell Embryo Average_RRBS
8-Cell Embryo (chr
Morula Average_RRBS
Morula (chr12-22)
ICM Average_RRBS
ICM (chr11-22)
Implanted Embryo_Average RRBS
Implanted Embryo (

Suppl. Fig. 8

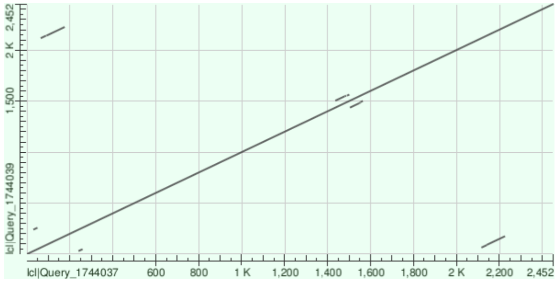


Mouse C5orf47

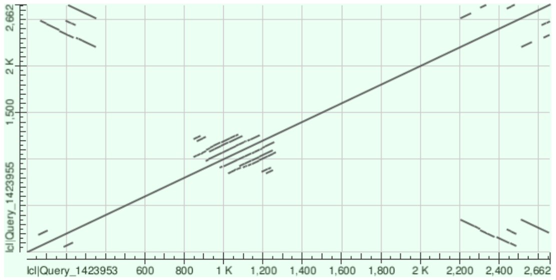


Human C5orf47

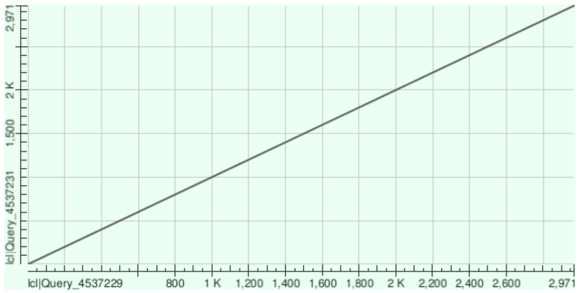


Mouse Dnaaf5

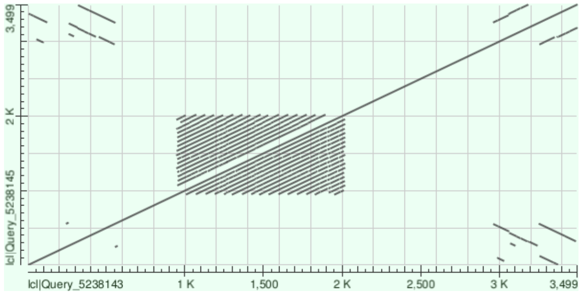


Human DNAAF5

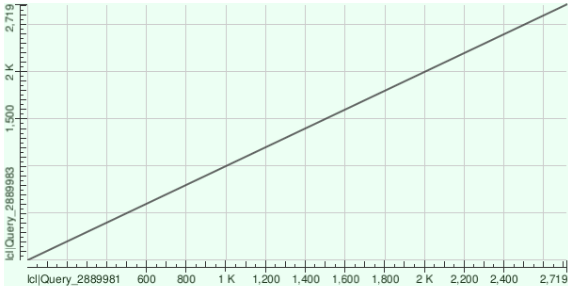


Mouse Slc45a4

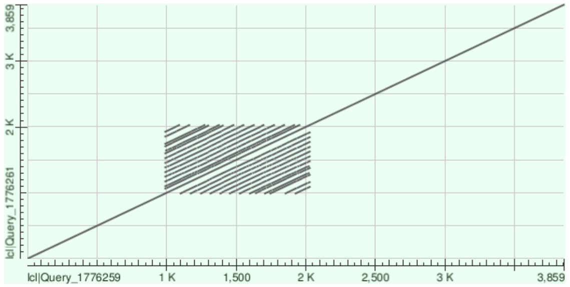


Human SLC45A4

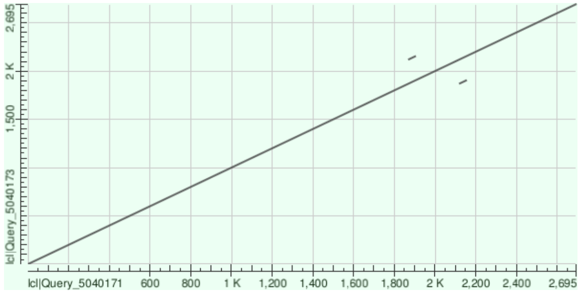

Mouse Proser2-as1

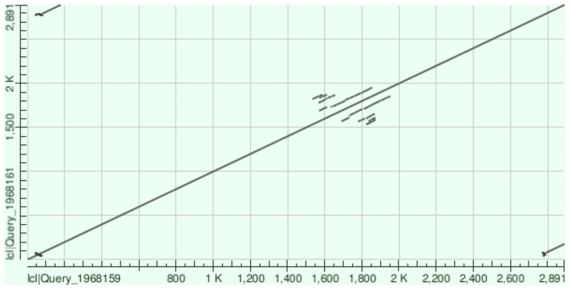

Human PROSER2-AS1

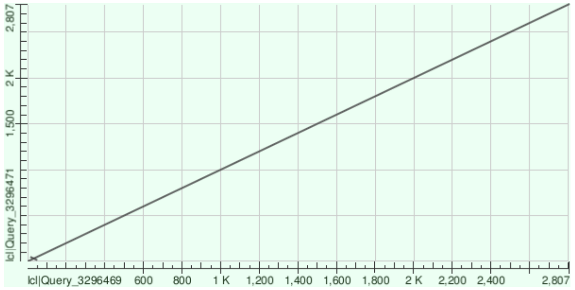

Mouse Shtn1 (5’)

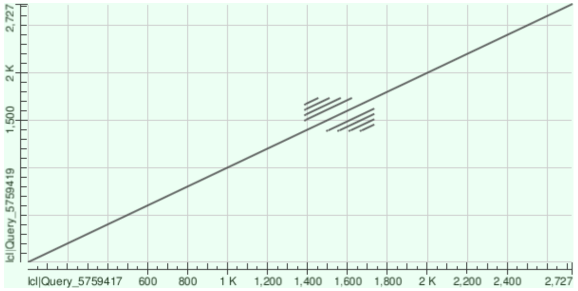

Human SHTN1 (5’)

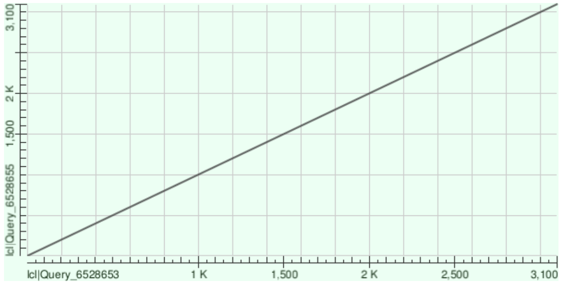

Mouse Kcng2

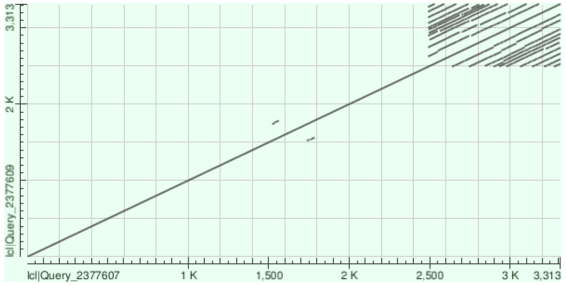

Human KCNG2

### Suppl. Fig. 9

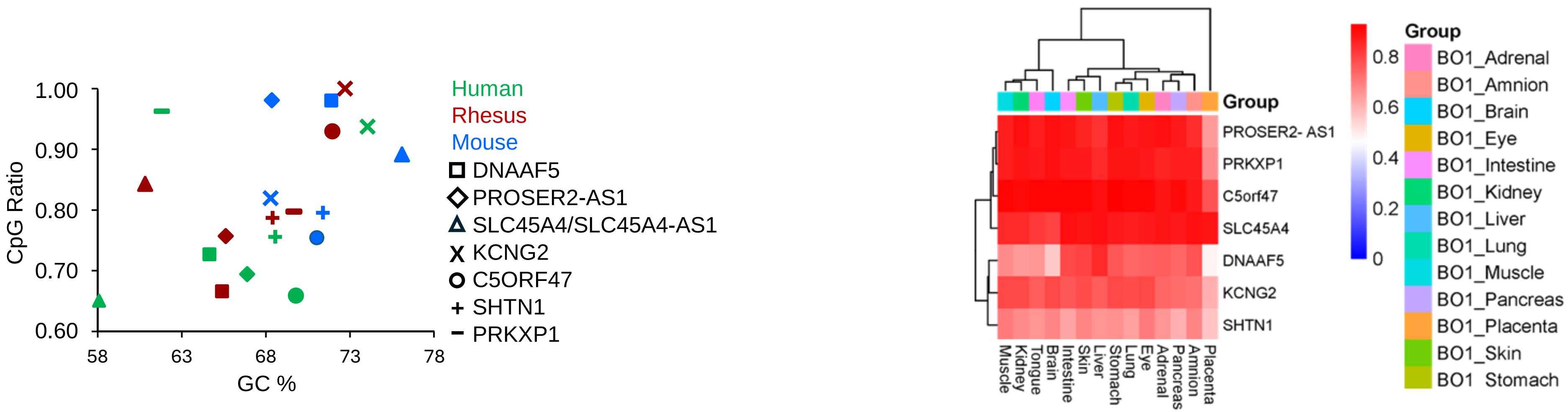
1.00
0.90
0.80
0.70
0.60
CpG Ratio
58
63
68
73
78
GC %
Human
Rhesus
Mouse
DNAAF5
PROSER2-AS1
SLC45A4/SLC45A4-AS1
KCNG2
C5ORF47
SHTN1
PRKXP1
Group
PROSER2- AS1
PRKXP1
C5orf47
SLC45A4
DNAAF5
KCNG2
SHTN1
Muscle
Kidney
Tongue
Brain
Intestine
Skin
Liver
Stomach
Lung
Eye
Adrenal
Pancreas
Amnion
Placenta
0.8
0.6
0.4
0.2
0
Group
BO1_Adrenal
BO1_Amnion
BO1_Brain
BO1_Eye
BO1_Intestine
BO1_Kidney
BO1_Liver
BO1_Lung
BO1_Muscle
BO1_Pancreas
BO1_Placenta
BO1_Skin
BO1 Stomach
